# SIN-3 transcriptional coregulator maintains mitochondrial homeostasis and polyamine flux

**DOI:** 10.1101/2023.08.07.552272

**Authors:** M. Giovannetti, P. Fabrizio, O. Nicolle, C. Bedet, MJ. Rodríguez-Palero, G. Michaux, M. Artal-Sanz, M. Witting, F Palladino

## Abstract

Mitochondrial function relies on the coordinated transcription of mitochondrial and nuclear genomes to assemble respiratory chain complexes. Across species, the SIN3 coregulator influences mitochondrial functions, but how its loss impacts mitochondrial homeostasis and metabolism in the context of a whole organism is unknown. Exploring this link is important because *SIN3* haploinsufficiency causes intellectual disability/autism syndromes and SIN3 plays an important role in tumor biology. Here we show that loss of *C. elegans* SIN-3 results in transcriptional deregulation of mitochondrial- and nuclear encoded mitochondrial genes, potentially leading to mito-nuclear imbalance. Consistent with impaired mitochondrial function, *sin-3* mutants show extensive mitochondrial fragmentation by transmission electron microscopy (TEM) and *in vivo* imaging, and altered oxygen consumption. Metabolomic analysis of *sin-3* mutant animals identifies a signature of mitochondria stress, and deregulation of methionine flux resulting in decreased S-adenosyl methionine (SAM), and increased polyamine levels. Our results identify SIN3 as a key regulator of mitochondrial dynamics and metabolic flux, with important implications for human pathologies.

## Introduction

Mitochondria, the main energy providers within cells, produce ATP through oxidative phosphorylation (OXPHOS) and control the levels of metabolites essential for various cellular functions. The OXPHOS system consists of four multimeric complexes, coenzyme Q and cytochrome *c* that form the mitochondrial respiratory chain (I–IV) and couple redox reactions, creating an electrochemical gradient leading to the creation of ATP through a fifth complex, the F_1_F_0_ ATPase (Signes and Fernandez-Vizarra, 2018). Assembly of functional mitochondria relies on coordinated transcription and translation of mitochondrial and nuclear genomes (Richter-Dennerlein et al., 2016), and both mitochondrial and nuclear DNA mutations affecting the accumulation and function of OXPHOS enzymes are the most common cause of mitochondrial diseases and are associated with neurodegeneration and aging (DiMauro and Schon, 2003; Fernandez-Vizarra and Zeviani, 2021; Reeve et al., 2008). The mitochondrial genome, which is highly conserved among species, comprises 37 genes coding for two ribosomal RNAs, 22 transfer RNAs, and 13 protein subunits (12 for *C. elegans*) of the mitochondrial respiratory chain performing OXPHOS (Taanman, 1999). The rest of the mitochondrial proteome, comprising over a thousand proteins, is encoded in the nucleus (Ali et al., 2019; Barshad et al., 2018; Mercer et al., 2011). The bidirectional regulation between mitochondria and the nucleus, referred to as mito-nuclear communication, maintains homeostasis and regulates stress responses (Quirós et al., 2016). Mitochondrial damage or alterations in mitochondrial function trigger specific quality control mechanisms. Of these, the best characterized is the mitochondrial unfolded protein response (UPR^mt^), a cellular stress response that leads to increased transcription of mitochondrial chaperones and proteases (Anderson and Haynes, 2020). In addition to an imbalance between mitochondrial and nuclear protein quantities (Matilainen et al., 2017), reduced levels of TCA cycle components (Yang et al., 2022), reduction of β-oxidation and lipid biosynthesis (Rolland and Conradt, 2022; Rolland et al., 2019) and defective mitochondrial import (Lionaki et al., 2022), can all trigger UPR^mt^. Recent data has shown that mitochondrial stress leads to extensive chromatin reorganization (Tian et al., 2016; Zhu et al., 2020), and chromatin regulatory factors contribute to mitochondrial gene expression (Matilainen et al., 2017). Conversely, metabolites originating from the mitochondria can initiate modifications in the nucleus (Matilainen et al., 2017).

Depletion experiments in various models have shown that the highly conserved SIN3/HDAC coregulator plays a role in mitochondrial functions and metabolism (Chaubal and Pile, 2018). Yeast *sin3* null mutants grow poorly on non-fermentable carbon sources, have lower ATP levels, and reduced respiration rates (Barnes et al., 2010). In Drosophila cultured cells, reduction of SIN3 levels also resulted in altered ATP levels and deregulation of genes encoded by the mitochondrial genome (Barnes et al., 2010; Pile et al., 2003). Lower levels of ATP and increased sensitivity to oxidative stress were also observed in *C. elegans* animals carrying the *sin-3(tm1276)* partial loss of function allele (Sharma et al., 2018). In mice, Sin3a was detected in a transcriptional complex with the MafA pancreatic β-cell-specific activator (Scoville et al., 2015), and its inactivation reduced the fitness of β-cells and altered glucose production in liver (Yang et al., 2020a). Significantly, in mammalian cells, SIN3 and its associated protein SUDS3 were identified in a screen for modifiers of drug-induced mitochondrial dysfunction (To et al., 2019). Altogether, these data from different systems support a conserved role for SIN3 in mitochondrial homeostasis and metabolism, but how SIN3 affects mitochondrial dynamics and metabolic pathways in the context of a whole organism remains largely unknown. This is particularly important given that heterozygous loss-of-function variants, as well as point mutations in *SIN3* were recently identified as the underlying cause of intellectual disability (ID)/autism syndromes (Balasubramanian et al., 2021; Latypova et al., 2021; Witteveen et al., 2016), and SIN3 levels play an important role in tumor biology (Bansal et al., 2015; Bansal et al., 2016; Farias et al., 2010; Lewis et al., 2016).

Previous studies in *C. elegans* have shown that knock-down of *sin-3*, the single SIN3 homolog in this organism, results in a decreased lifespan, altered mitochondrial membrane potential, enhanced autophagy and increased oxidative stress (Pandey et al., 2018; Sharma et al., 2018). How these changes affect mitochondrial morphology and function was not investigated. Here, using a *sin-3* CRISPR-Cas9 knock-out allele, we show that loss of SIN-3 results in the deregulation of both mitochondria*-* and nuclear-encoded genes. Transmission electron microscopy (TEM) and *in vivo* imaging revealed extensive fragmentation of mitochondria in all tissues examined, including muscle, intestine, hypodermis, and the germline. Consistent with severe defects in mitochondrial function, both basal and maximal oxygen consumption are increased in the absence of SIN3, while spare respiratory capacity is decreased. Metabolomic analysis identified a signature of mitochondria stress and deregulation of methionine flux, resulting in decreased levels of SAM and a shift towards higher polyamine levels. Together our data identify SIN3 as an important regulator of mitochondrial dynamics in an organismal context, and reveal a SIN-3 dependent connection between expression of OXPHOS subunits, mitochondrial homeostasis, and metabolic fluxes.

## Results

### Loss of *sin-3* results in altered expression of genes with mitochondrial functions

Previous transcriptomic profiling of animals carrying the *sin-3(tm1276)* partial loss-of-function allele revealed deregulation of the germline transcriptome, including metabolic genes (Caron et al., 2023). In order to identify high confidence genes regulated by SIN-3, we extended our analysis to genes commonly misregulated in *sin-3(tm1276)* and *sin-3*(*syb2172*) mutant animals that carry a complete loss of function allele obtained by CRISPR-Cas9 (Caron et al., 2023), generating a list of 892 genes (Table S1). Using Worm Cat (Holdorf et al., 2020), within this set we identified a common class of genes with functions related to mitochondria (Figure 1A and Table 1).

**Figure 1.**
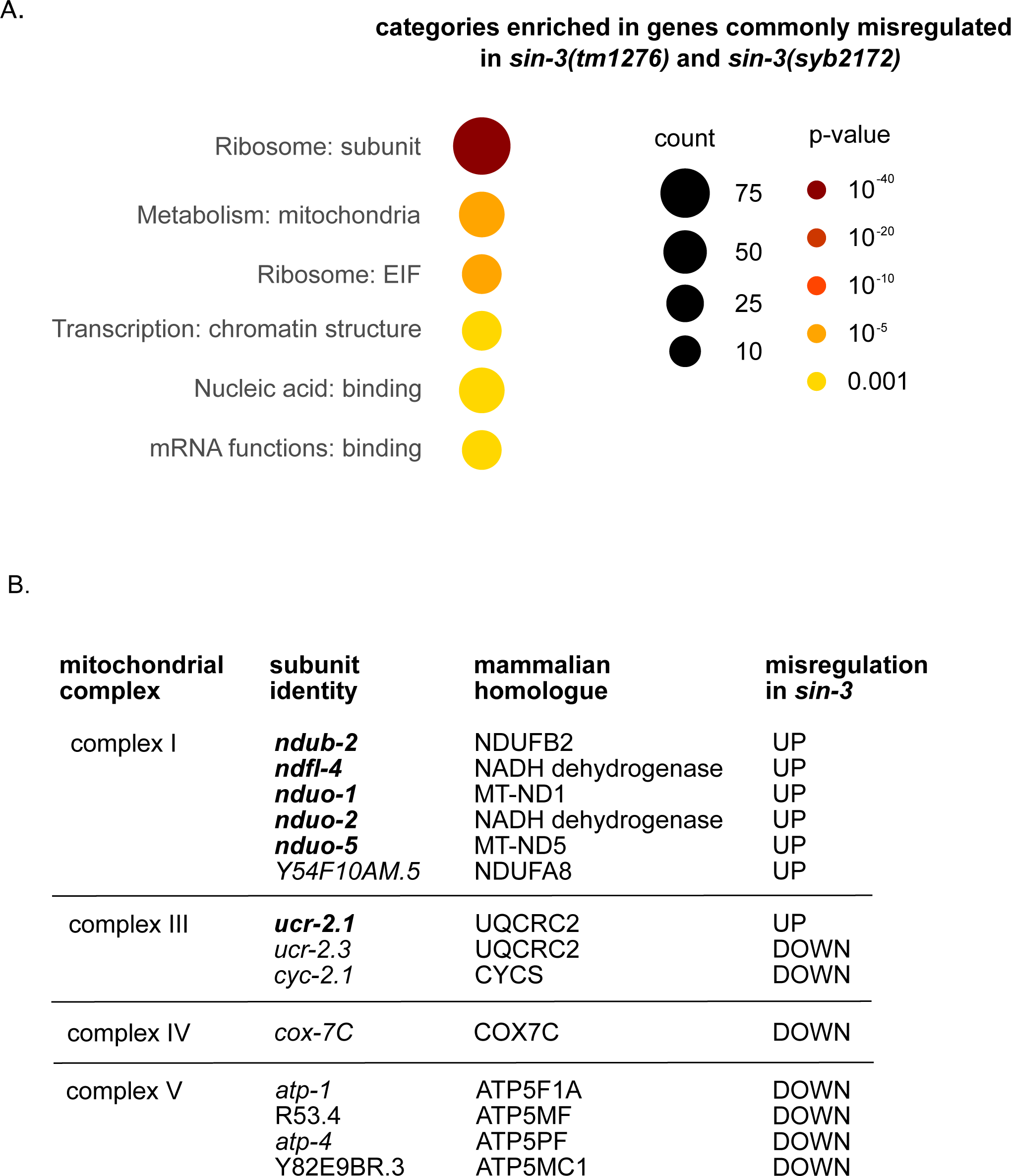
Loss of *sin-3* alters the expression of mitochondrial genes. (A) WormCat visualization of categories enriched in genes commonly misregulated in *sin-3(tm1276)* and *sin-3(syb2172)* mutants. The legend for bubble charts is indicated on the right, with size referring to the number of genes in each category and color referring to the p-value. (B) Respiratory complex subunits commonly misregulated in *sin-3(tm1276)* and *sin-3(syb2172)* mutants are indicated, along with their mammalian counterparts. Subunits encoded by the mitochondrial genome are shown in bold.

**Table 1.** List of genes with mitochondrial function commonly misregulated in both *sin-3(syb2172)* and *sin-3(tm1276)* germlines. The list of all commonly misregulated genes was generated by crossing the set of genes obtained for each allele with a FDR <0.05 following DESeq2 analysis (Table S1), and the list of mitochondrial genes extracted from WormCat.

| sequence_ID | category | misregulation in *sin-3(syb2172)* |
| --- | --- | --- |
| F54C9.6 | Metabolism: mitochondria: chaperone | DOWN |
| C35D10.5 | Metabolism: mitochondria: chaperone | DOWN |
| F25H9.7 | Metabolism: mitochondria: citric acid cycle | DOWN |
| F59B8.2 | Metabolism: mitochondria: citric acid cycle | DOWN |
| C50F7.4 | Metabolism: mitochondria: citric acid cycle | DOWN |
| F23H11.3 | Metabolism: mitochondria: citric acid cycle | DOWN |
| **ndub-2** | **Metabolism: mitochondria: complex I** | **UP** |
| **MTCE.4** | **Metabolism: mitochondria: complex I** | **UP** |
| **MTCE.11** | **Metabolism: mitochondria: complex I** | **UP** |
| **MTCE.16** | **Metabolism: mitochondria: complex I** | **UP** |
| **MTCE.35** | **Metabolism: mitochondria: complex I** | **UP** |
| Y54F10AM.5 | Metabolism: mitochondria: complex I | UP |
| VW06B3R.1 | Metabolism: mitochondria: complex III | UP |
| E04A4.7 | Metabolism: mitochondria: complex III | DOWN |
| T24C4.1 | Metabolism: mitochondria: complex III | DOWN |
| F26E4.6 | Metabolism: mitochondria: complex IV | DOWN |
| H28O16.1 | Metabolism: mitochondria: complex V | DOWN |
| R53.4 | Metabolism: mitochondria: complex V | DOWN |
| T05H4.12 | Metabolism: mitochondria: complex V | DOWN |
| Y82E9BR.3 | Metabolism: mitochondria: complex V | DOWN |
| F13B9.8 | Metabolism: mitochondria: morphology | UP |
| Y71G12B.24 | Metabolism: mitochondria: protease | UP |
| C25A1.13 | Metabolism: mitochondria: ribosome | DOWN |
| T13H5.5 | Metabolism: mitochondria: ribosome | DOWN |
| T21B10.1 | Metabolism: mitochondria: ribosome | DOWN |
| Y48C3A.10 | Metabolism: mitochondria: ribosome | DOWN |
| F56D1.3 | Metabolism: mitochondria: ribosome | DOWN |
| T14B4.2 | Metabolism: mitochondria: ribosome | DOWN |
| T20F5.3 | Metabolism: mitochondria: ribosome | DOWN |
| Y37E3.8 | Metabolism: mitochondria: ribosome | DOWN |
| F32A7.4 | Metabolism: mitochondria: RNA methyltransferase | DOWN |
| C18E9.6 | Metabolism: mitochondria: translocase | DOWN |
| Y67H2A.4 | Metabolism: mitochondria: transporter | DOWN |
| Y57G11C.11 | Metabolism: mitochondria: ubiquinone | DOWN |
| K08E3.7 | Metabolism: mitochondria: unassigned | DOWN |
| F27D4.1 | Metabolism: mitochondria: unassigned | DOWN |
| K12G11.3 | Metabolism: mitochondria: unassigned | DOWN |
| T06D8.6 | Metabolism: mitochondria: unassigned | DOWN |
| ZK1010.2 | Metabolism: mitochondria: unassigned | DOWN |
| T14G11.3 | Metabolism: mitochondria: unassigned | UP |
| C27H6.9 | Metabolism: mitochondria: unassigned | DOWN |
| F32A7.4 | Metabolism: mitochondria: unassigned | DOWN |

Both nuclear and mitochondrial encoded genes contribute to the assembly of mitochondrial respiratory chain complexes (MRC) I-V (Signes and Fernandez-Vizarra, 2018). Five of the 7 MRC complex I subunits encoded by the mitochondrial genome were strongly upregulated in both *sin-3* mutants: *ndub-2*, *ndfl-4, nduo-1, nduo-*2 and *nduo-5* (Figure 1B). By contrast, nuclear encoded MRC subunits identified in our data set were mostly down-regulated (Figure 1B), as were the majority of additional nuclear encoded genes with mitochondria-related functions (31 out of 37, Table 1). These include mitochondrial ribosomal proteins, components of the citric acid cycle, the *tomm-40* translocase and the *coq-3* coenzyme Q3 (Table 1). Comparison of our list of misregulated genes associated with mitochondrial functions to a list of SIN3 targets on chromatin (Beurton et al., 2019) revealed that 23 of these 37 genes have SIN-3 binding at their promoter region, and the majority (18/23) are downregulated in *sin-3* mutants, including ribosomal protein genes, the t*omm-40* mitochondrial translocase, the ATP synthases *atp-1* and *atp-4,* and the cytochrome-C oxidase *cox-7C* (Table S2). SIN-3 may therefore directly promote expression of these genes. Together, our analyses suggest that loss of *sin-3* perturbs the coordinated expression of mitochondrial genes encoded by nuclear and mitochondrial genomes, possibly resulting in mito-nuclear imbalance and affecting mitochondrial homeostasis. Consistent with mitochondrial dysfunction, *sin-3* mutants show reduced fertility, increased sensitivity to oxidative stress, and altered lifespan (Beurton et al., 2019; Caron et al., 2023; Pandey et al., 2018; Sharma et al., 2018), as observed in other mitochondrial mutants (Kropp et al., 2021).

### Transmission electron microscopy reveals extensive mitochondrial fragmentation in *sin-3* mutant animals in all tissues examined

Mitochondrial morphology is tightly linked to mitochondrial function, and its steady state is determined by the balance between fission and fusion events that may be disrupted under conditions of mitochondrial stress (Westermann, 2010). To test whether mitochondrial morphology is affected in *sin-3* mutant animals we performed electron microscopy analyses on wildtype and mutant young adults. Mitochondria in the body wall muscles of wild type animals vary in size reflecting dynamic morphological changes, and tend to be cylindrical- or ovular shaped with regular outer membranes and dense cristae throughout each organelle (Figure 2 panel *a*, white arrows). In animals carrying the loss of function allele *sin-3(tm1276),* or the *sin-3(syb2172)* null allele, mitochondria were instead highly fragmented and appeared more numerous, with a stronger effect in *syb2172* animals (Figure 2 panels *b* and *c*, white arrows). For both mutants, highly fragmented mitochondria were visible in all of the tissues examined, including intestine, pharynx, hypodermis, and germline (Figure 2A and Figure S1A). We also observed the presence of enlarged or “giant” mitochondria, most prominent in muscle and intestinal cells, and often containing electron-dense material (Figure 2A panels *b, c*, and *e*, Figure 2B panels *a-d*, black arrows) that may indicate iron deposits (Puccio et al., 2001; Schober et al., 2021) (Schiavi et al., 2015) or protein aggregates (Chen et al., 2021) (Al Rawi et al., 2011; Wang et al., 2016; Zhou et al., 2016). Selective autophagy of mitochondria, or mitophagy, is a process that results in budding off of damaged mitochondria for targeted breakdown, thereby contributing to mitochondrial homeostasis (Onishi et al., 2021). While this process was never observed in wild type, examples of double-membrane autophagosomes were observed in both the hypodermis and germ cells of mutant animals (Figure S1B). Together, these results show that *sin-3* inactivation dramatically alters mitochondrial dynamics and morphology, and reveal increased mitochondrial fission throughout the body of *sin-3* mutant animals, with the most severe phenotypes observed in the null mutant allele.

**Figure 2.**
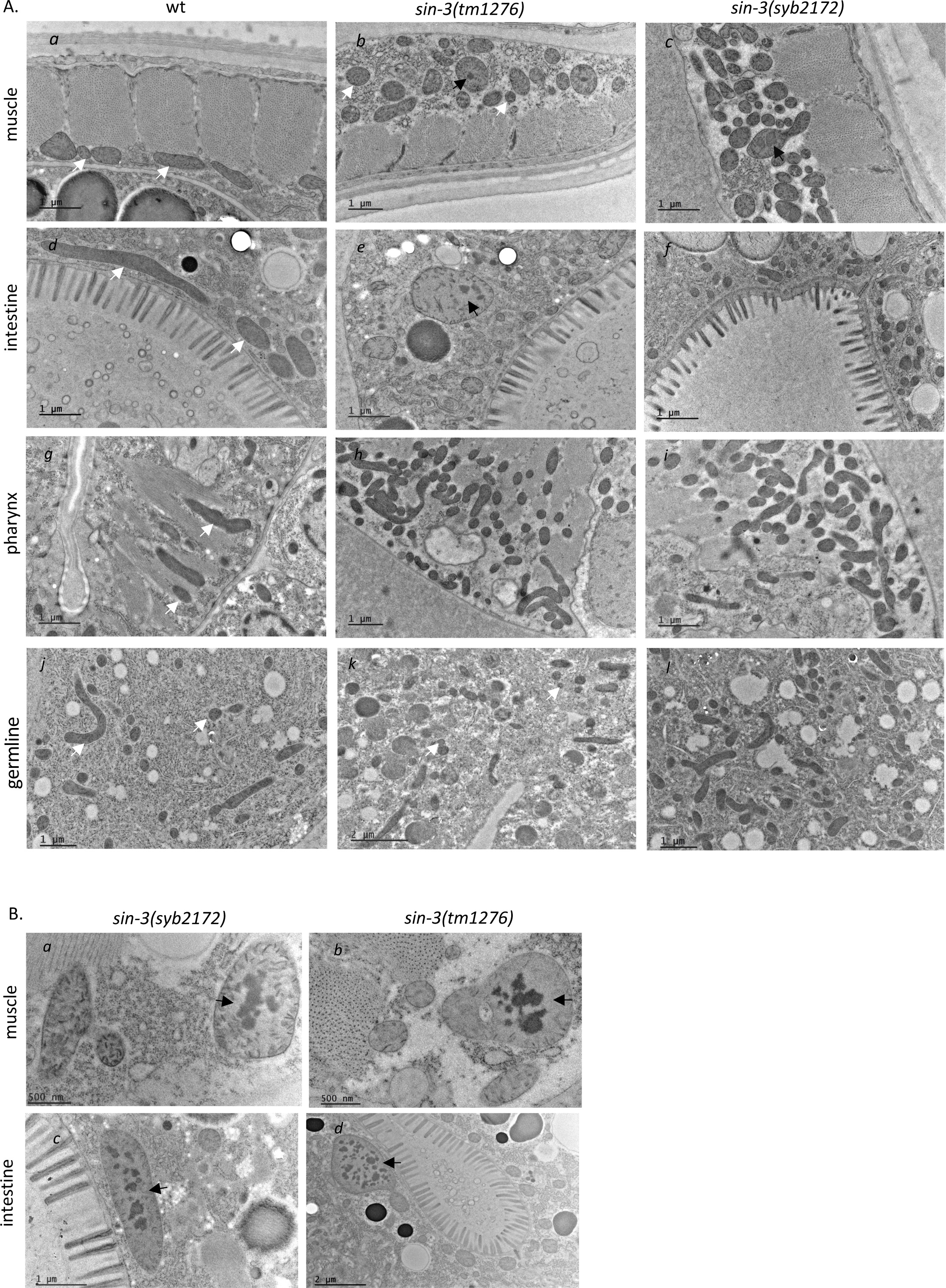
TEM microscopy of mitochondria in *sin-3* mutant animals. Transverse sections captured with electron microscopy of mitochondria from the body wall muscles, intestine, pharynx and germline of young adult wild type, *sin-3(tm1276)* and *sin-3*(*syb2172*) mutant animals. Images are representative of a total of 6 animals imaged for wild type, 4 for *sin-3*(*syb2172*) and 5 for *sin-3*(*syb2172*). Further examples for each genotype are displayed in Supplementary Figure 1. White arrows indicate examples of individual mitochondria, black arrows examples enlarged mitochondria with densely stained aggregates. Scale bars are indicated for each image.

### Time dependent increase in mitochondrial fragmentation in *sin-3* muscle cells

To further examine and quantify changes in mitochondrial morphology in live animals, we used a transgenic strain expressing a red fluorescent protein (RFP) fused at the N terminus to the TOMM-20 translocase of the outer mitochondrial membrane and expressed under the control of a promoter in muscle cells (*myo-3p::20Nter::*wrmScarlet)(Roy et al, 2022), where mitochondria are abundant and easily visible. We used *tm1276* mutant animals because although less fertile than wildtype, they can be maintained as homozygotes, while *syb2172* animals are fully sterile (Caron et al., 2023). In wildtype, the majority of body-wall muscle cells have longitudinally arrayed tubular mitochondria, while a smaller percentage show either elongated mitochondria in an interconnected mesh-like network or fragmented mitochondria (Figure 3A). We arbitrarily classified mitochondrial morphology into three classes-tubular, intermediate, or fragmented-(Regmi et al., 2014), and scored animals falling into each of these at days 1, 6, and 9 of adulthood.

**Figure 3.**
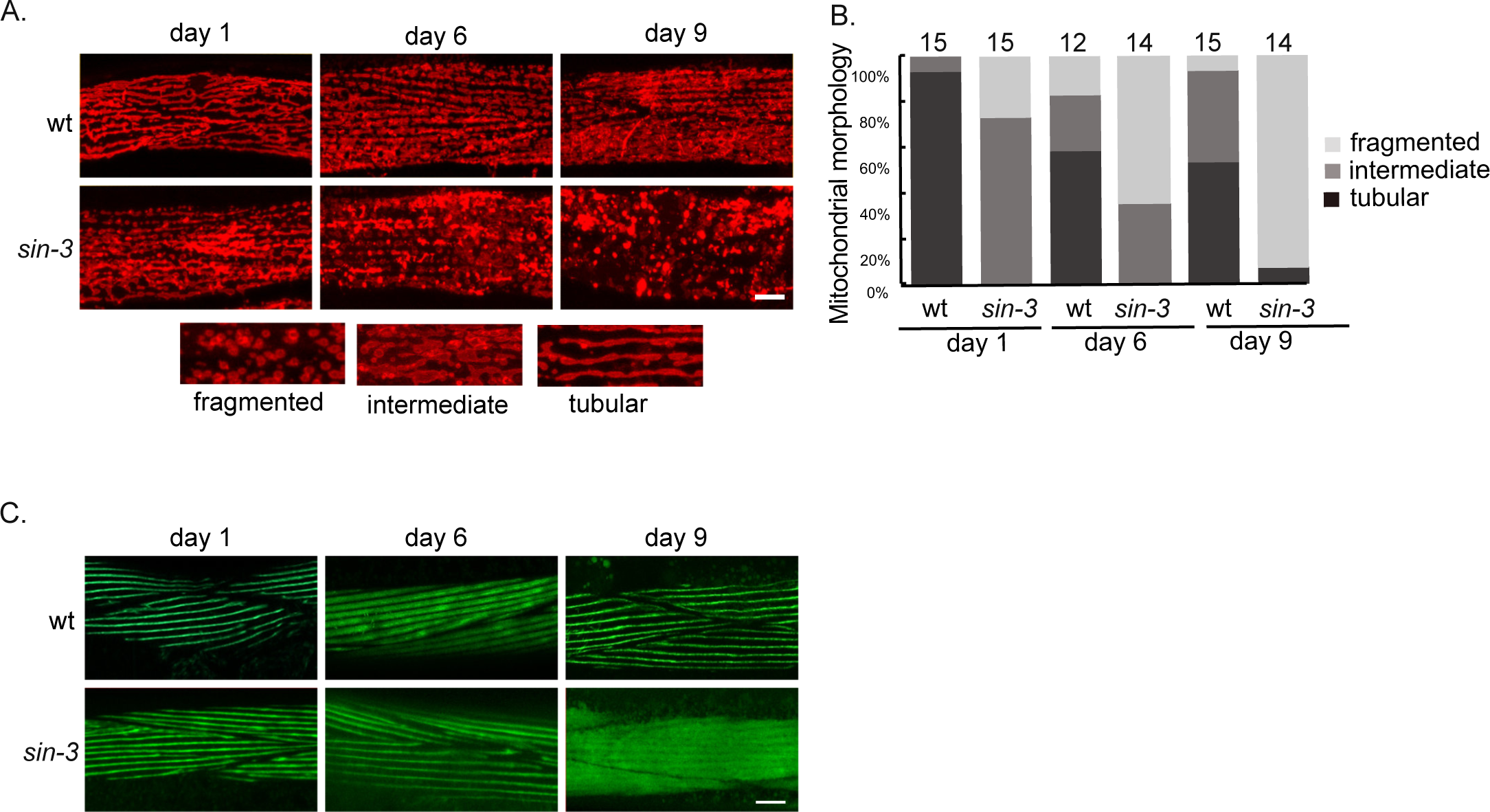
Altered mitochondria morphology and microfilament structure in *sin-3(tm1276)* muscle cells. (A) Representative confocal images of mitochondria morphology in body wall muscles expressing *myo-3p::tomm-20*Nter::wrmScarlet in wildtype and *sin-3(tm1276)* mutant animals. Images were taken at day 1, 6 or 9 of adulthood. Scale bar: 10µm. (B) Quantification of animals with the indicated muscle mitochondria phenotypes at day 1, 6 or 9 of adulthood. Total number of worms scored for each condition is indicated above bars. Phenotypes are described in Materials and Methods. (C) Representative confocal images of myofilament structure in body wall muscles expressing GFP:*:myo-3* in wildtype or *sin-3(tm1276)* mutant animals. Images were taken at day 1, 6 or 9 of adulthood. Scale bar: 10µm.

In wild type day one adults, the large majority of mitochondria showed a tubular morphology (Figure 3A and B). In *sin-3* mutants of the same age, we instead mainly observed “intermediate” mitochondrial morphology, with a significant percentage of highly fragmented mitochondria. At day 6, intermediate and fragmented mitochondria also appeared in wild type, while the majority of mitochondria in *sin-3* mutants were highly fragmented. At day 9, mitochondrial fragmentation further increased in *sin-3(tm1276)* mutants, but remained more or less constant in wildtype (Figure 3B). Using a *mex-5p*::tomm-20::mKate2 reporter expressed in the germline, we also observed changes in mitochondrial morphology in this tissue (Figure S1C). Excessive mitochondrial fragmentation results from an imbalance between fusion and fission that are mediated primarily by two classes of GTPases structurally related to dynamins: mitofusins (MFNs)/Fzo1/FZO-1 and Drp1/DRP-1, respectively (Chen and Chan, 2009; Labrousse et al., 1999; Westermann, 2010). Expression of neither gene was altered in *sin-3* mutants (Caron et al., 2023, Table S1), suggesting that the observed defects in mitochondrial morphology may arise instead as a response to mitochondrial stress.

### Muscle fibers degenerate prematurely in *sin-3* mutants

Mitochondrial dysfunction impairs muscle health and causes subsequent muscle wasting, commonly referred to as sarcopenia (Christian and Benian, 2020). Using a muscle myosin reporter *myo-3::*GFP (Mergoud Dit Lamarche et al., 2018) in wildtype young adults sarcomeres appear as straight lines of GFP, and no obvious change was observed up to day 9 (Figure 3C). By contrast, in *sin-3(tm 1276)* mutants at day 9 we observed diffuse GFP fluorescence, suggesting that muscle integrity is affected prematurely in these mutants (Figure 3C). Degradation of muscle proteins may also occur prematurely in these animals (Liang et al., 2014). Because defects in muscle fibers are observed only in older *sin-3* mutant adults, while mitochondrial defects already appear at day 1, these results are consistent with a decline in mitochondrial network structure preceding muscle decline, as previously reported (Etheridge et al., 2015; Gaffney et al., 2018).

### Mitochondrial UPR is dampened in the absence of SIN-3

In response to mitochondrial stress, cells trigger the mitochondrial unfolded protein response (UPR^mt^), a transcriptional response that transmits mitochondrial stress signals to the nucleus to regulate mitochondrial chaperone genes and other factors necessary for the recovery of dysfunctional mitochondria (Lin and Haynes, 2016). Increased mitochondrial fragmentation in *sin-3* mutants suggests that SIN-3 may be required for mitochondrial homeostasis. We therefore asked whether UPR^mt^ is properly activated in *sin-3* mutant animals, using the *hsp-6* chaperone as a reporter (Heussler et al 2022). *sin-3(tm1276)* mutants carrying *hsp-6*::GFP were sterile, so we used RNAi to deplete *sin-3. sin-3*(RNAi) alone had no reproducible effect on *hsp-6:*:GFP expression (Figure 4A). Induction of mitochondrial stress by RNAi knock-down of the NADH ubiquinone oxidoreductase component *nuo-4*, or the mitochondrial ribosomal protein *mrps-5,* resulted in strong *hsp-6*::GFP expression in the intestine, as expected (Houtkooper et al., 2013). Simultaneous depletion of both *sin-3* and *nuo-4,* or *sin-3* and *mrps-5,* significantly decreased GFP expression, but only in older animals at day 6 (Figure 4A). No significant difference in *hsp-6*::GFP induction between *sin-3* and wildtype was observed at day 3 (Figure S2). These results suggest that SIN-3 is required for a full response to mitochondrial stress depending of the age of the animals. Interestingly components of the NuRD chromatin remodeling complex have been shown to relocate to the nucleus upon mitochondrial stress (Zhu et al., 2020) to influence the UPR^mt^. Using a SIN-3::RFP translational fusion, we observed no obvious change in SIN-3 localization following mitochondrial stress (Figure S3).

**Figure 4.**
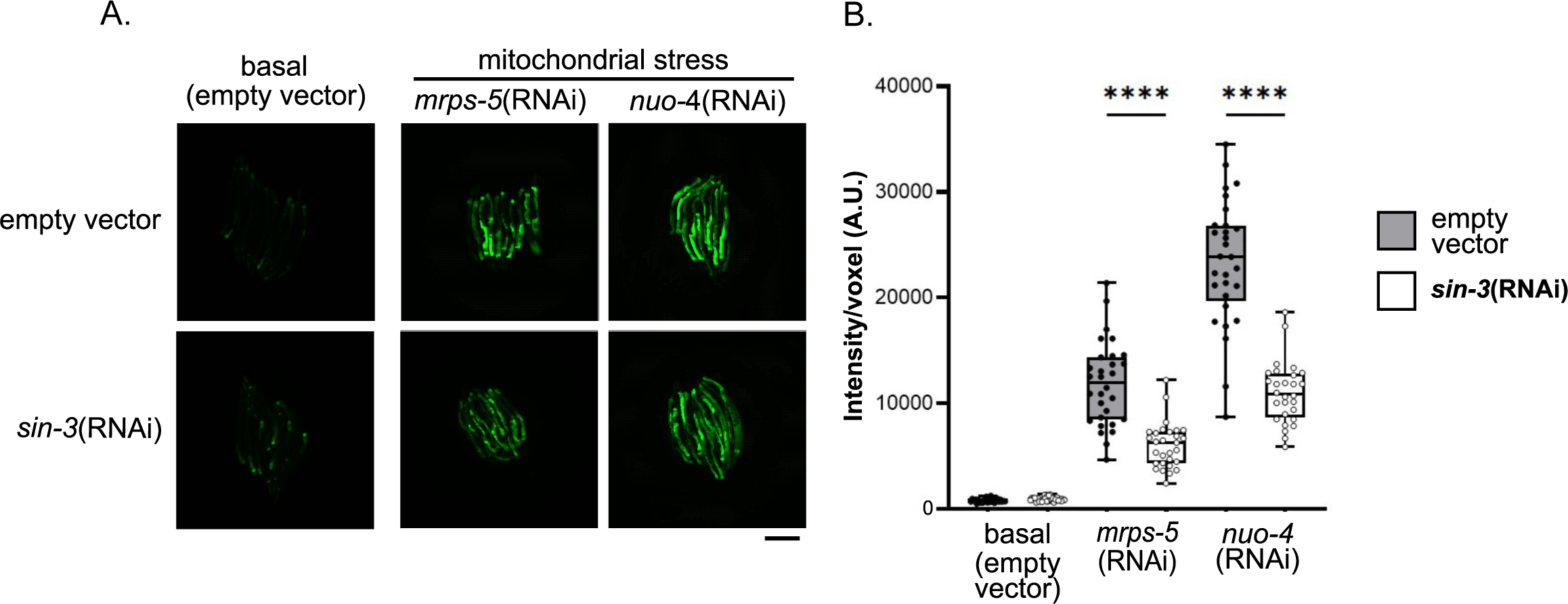
Loss of *sin-3* dampens the response to mitochondrial stress. (A) Images of wildtype or *sin-3*(RNAi) animals expressing *hsp-6p::GFP* reporter under basal conditions, or following mitochondrial stress induced by RNAi knock-down of mitochondrial genes *nuo-4* and *mrps-5*. Parent worms were grown on HT115 control or *sin-3* RNAi from hatch until the adult stage. F1 offspring were then grown from hatch on control, *nuo-4* or *mrps-5* single RNAi, or *sin-3*, *sin-3* + *nuo-4*, and *sin-3* + *mrps-5* double RNAi. GFP expression was measured 5 days after eggs were laid. Scale bar: 300µm. (B) Quantification of *hsp-6p::*GFP expression. Data is representative of one of two independent experiments with 30 animals per condition each. Significance was calculated using Mann-Whitney test, P-value * < 0.05, **** < 0.0001.

### Increased oxygen consumption in aged *sin-3* mutant animals

To understand how altered mitochondrial structure following loss of SIN-3 relates to mitochondrial function, we measured oxygen consumption in *sin-3(tm1276)* mutants. In *C. elegans*, oxygen consumption rates (OCRs) as a measure of ETC function can be accurately assessed using a Seahorse Analyzer with intact animals (Koopman et al., 2016; Kropp et al., 2021). Wildtype animals maintained a constant basal OCR from the young adult stage to day 6 of adulthood, although a tendency to reduce basal respiration during aging was apparent, as previously reported (Figure 5A and B, compare YA to D6) (Higashitani et al., 2022; Huang and Lin, 2018). By contrast, as *sin-3* mutants aged, basal OCR dramatically increased, so that at day six of adulthood it was more than 2-fold higher in *sin-3* mutants compared to wild type (Figure 5A, D6).

**Figure 5.**
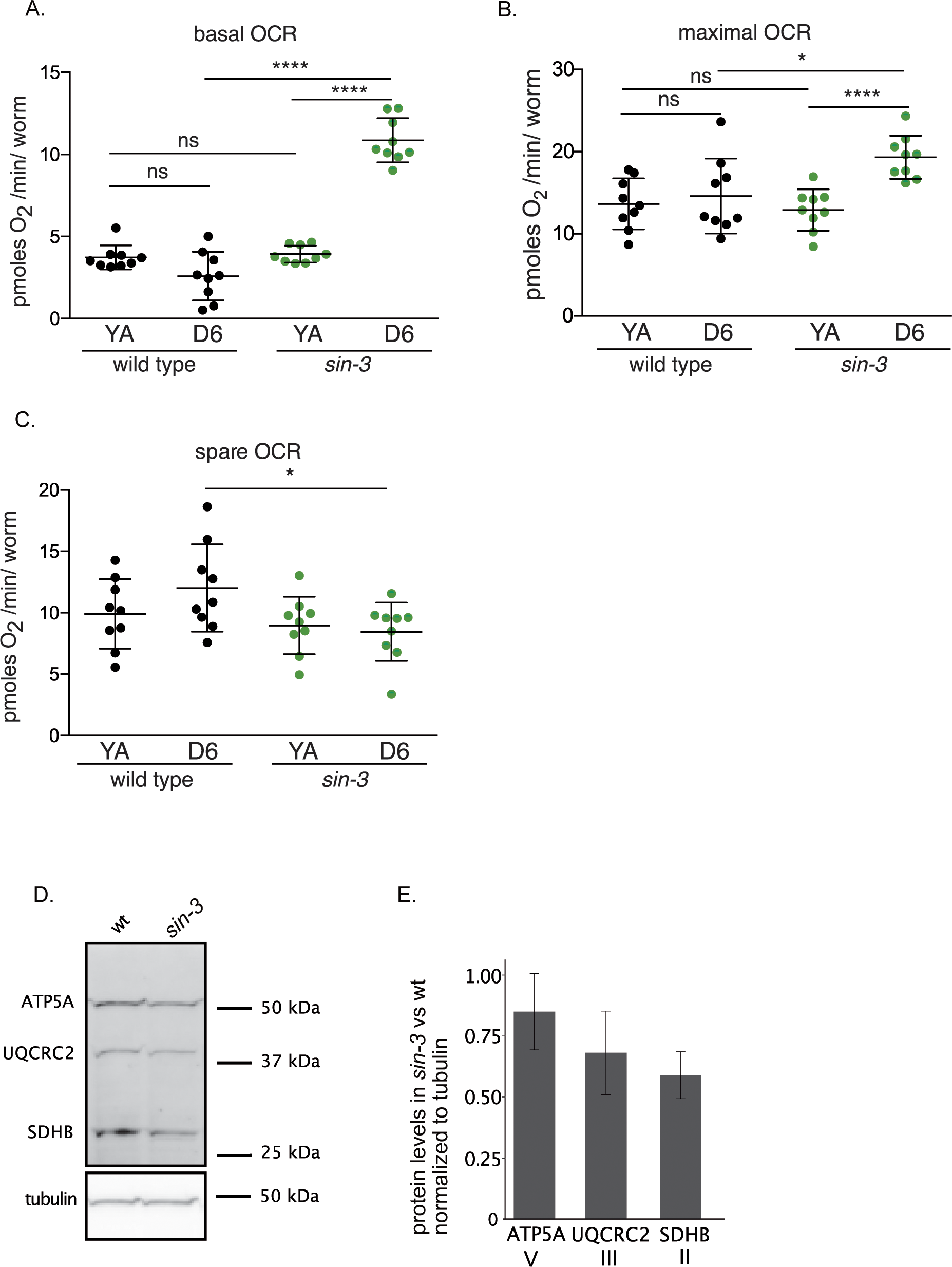
Oxygen consumption rate (OCR) and abundance of mitochondrial respiratory chain subunits in *sin-3(tm1276)* mutants. (A) Basal OCR of wild type and *sin-3* mutants at the young adult (YA) stage and at day 6 of adulthood (D6). (B) Maximal OCR of wild type and *sin-3* mutants at the YA stage and at D6 of adulthood. (C) Spare respiratory capacity of wild type and *sin-3* mutants at the YA stage and at D6 of adulthood. Significance was calculated using unpaired *t*-test or, when appropriate, Mann-Whitney test, P-value * < 0.05, **** < 0.0001. (D) Western blot showing SDHB-1 (complex II), CTC-1 (complex IV), and ATP-2 (complex V) protein levels in total protein extracts from wildtype and *sin-3(tm1276)* L4-young adult worms detected by fluorescence. (F) Levels of each protein were quantified with Image J and normalized to tubulin levels. Bar plot represents the mean of two to three independent biological replicates; error bars correspond to SD.

We next measured maximal respiratory capacity following mitochondrial uncoupling by the addition of carbonyl cyanide-p-trifluoromethoxyphenylhydrazone (FCCP). In *sin-3* young adults, maximal OCR was similar to wild type, but we again observed a significant increase in aged *sin-3* mutants compared to age-matched wildtype (Figure 5B). Moreover, in old *sin-3* mutants the observed increase in basal respiration was proportionally larger than the increase in maximal respiratory capacity (Figure 5A and B). Consequently mitochondrial spare capacity, which characterizes the ability of mitochondria to meet extra energy requirements beyond the basal level (Marchetti et al., 2020), was reduced compared to old wildtype worms (Figure 5C). Increased OCR could be a side effect of partial mitochondrial uncoupling (Demine et al., 2019). We note however that the single mitochondrial uncoupling protein in *C. elegans, ucp-4,* is down-rather than upregulated in both *sin-3* mutants (Table S1), as is the expression of the *ant-1.1* and *ant-1.3* ADP/ATP translocators whose activity can also result in mitochondrial uncoupling (Berry et al., 2022; Bertholet et al., 2019). Consistent with depletion of SIN-3 affecting mitochondrial integrity, we observed that levels of conserved electron transport proteins, detected using antibodies against the highly conserved mammalian complex III subunit UQCRC2 and complex II subunit SDHB1, were reproducibly decreased in extracts from *sin-3(tm1276)* animals. Protein abundance of the ATP synthase ATP5 was less affected (Figure 5D and E).

### *sin-3* inactivation does not alter levels of TCA cycle metabolites, but results in a signature of mitochondrial stress

The above results clearly establish that *sin-3* knock-down or deletion leads to defective mitochondrial function. Because mitochondria are a hub for biosynthetic processes, we used metabolomic analysis to probe for SIN-3-dependent metabolic changes in animals carrying the *sin-3(tm1276)* loss of function allele. We chose this allele because the sterility of *sin-3(syb2172)* null mutants precluded growing the large number of animals required for this type of analysis. We observed no significant change in the abundance of TCA cycle intermediates detected, including succinyl-CoA, alpha-keto-glutaric acid, succinic acid, fumaric acid, and malic acid (Tables S4 and S5, Figures S4 A and B). Although two peaks putatively annotated as glycolytic intermediates glucose-6-P and glucose-1-P showed a significant reduction in *sin-3* mutants, the downstream metabolite fructose-1,6-bis-P showed no difference (Table S4 and Figure S4B). Glucose-1-P is a metabolite of gluconeogenesis and starch and sucrose metabolism. *C. elegans* can store excess energy in the form of trehalose and glycogen (Hanover et al., 2005). However, we detected no significant change in UDP-glucose (an intermediary for glycogen and trehalose synthesis), trehalose-6-P, or trehalose in *sin-3* mutant animals (Table S4 and Figure S4B). Together, these results suggest that the absence of SIN-3 has little or no influence on energy metabolism. This was further corroborated by only minor changes in lipid metabolism intermediates including short-chain acyl-CoA and short-chain acyl-carnitine species (Table S5 and Figure S4A). Nonetheless, it is important to note that metabolite levels are only indicative of changes in metabolic flux, defined as the turnover rate of metabolites through enzymatically controlled pathways (Nielsen, 2003). Increased mitochondrial fragmentation may lead to changes or redirection of metabolic fluxes without altering metabolite levels.

In contrast to the above results, significant changes in amino acid metabolism were detected: aspartic acid levels significantly decreased in *sin-3* mutants, while proline, threonine, lysine, serine, and citrulline increased (Figure 6A and Table S5 and Figure S4C). Interestingly, decreased aspartic acid and increased serine have been shown to be a signature of mitochondrial stress (Quirós et al., 2017). No changes in glutathione (GSH) and glutathione disulfide (GSSG), both associated with oxidative stress, were detected (Table S5 and Figure S4C). An increase in sarcosine and dimethylarginine was also detected (Table S5 and Figure S4C).

**Figure 6.**
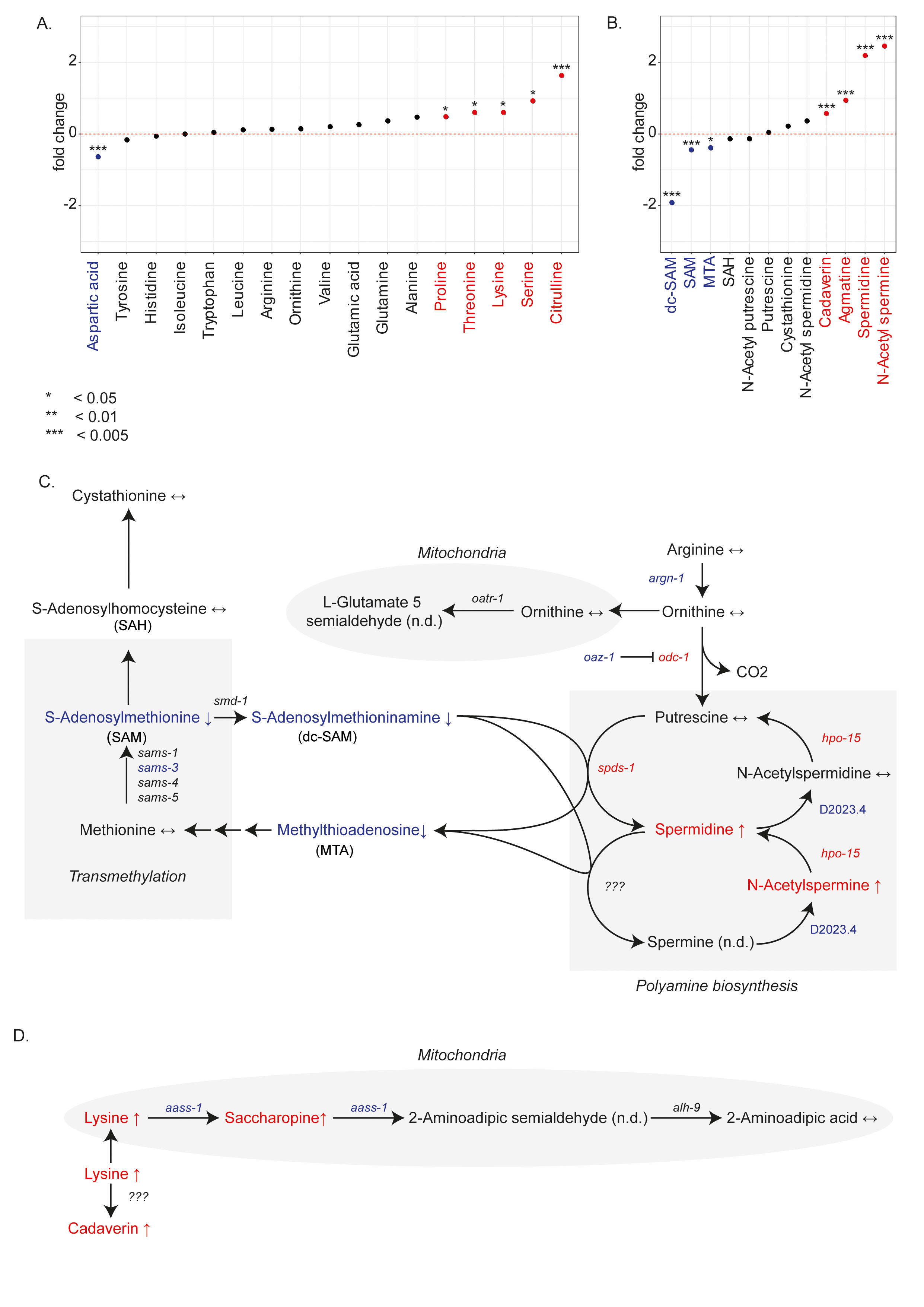
Metabolic alterations in *sin-3(tm1276)* mutant animals and their related pathways. Changes in amino acid metabolites (A) and polyamine-related metabolites (B) in *sin-3(tm1276)* mutants at the young adult stage. Welch test, P-value * < 0.05, ** < 0.01, *** < 0.005. (C) Schematic representation of biosynthetic pathway for arginine-derived polyamines. (D) Biosynthetic pathway of cadaverine and lysine degradation. Arrows and colors in (C) and (D) indicate significant changes for metabolites and metabolic genes (blue = down, red = up). Specific pathways and reactions that take place in the mitochondria are shaded in gray.

### SIN-3 is required to maintain polyamine homeostasis

Both serine and sarcosine are connected to one-carbon metabolism, which encompasses both the folate and methionine cycles to generate one-carbon units (methyl groups) that are used for the biosynthesis of important precursors and for methylation reactions (Locasale, 2013). While no changes in folate cycle intermediates folate, 5-methyl tetrahydrofolic (mTHF) or dihydrofolic (DHF) acid were found (Table S5 and figure S4D), a significant decrease in the methionine cycle metabolite S-adenosylmethionine (SAM) was observed (Figure 6B, Table S5 and Figure S4D).

SAM is the major methyl donor in the cell and is critical for the synthesis of phosphatidylcholine, an important component of cellular membranes (Ye et al., 2017), as well as for the methylation of DNA, RNA, and histones through transmethylation (Locasale, 2013). SAM is also connected to polyamine synthesis through the activity of *S*-adenosylmethionine decarboxylase (adoMETDC/*smd-1*) that results in the production of decarboxylated *S*-adenosylmethioninamine (dc-SAM) (Figure 6C). Spermidine synthase (SPDS/*spds-1*), a key enzyme in the pathway, then catalyzes the transfer of an aminopropyl moiety of dcSAM to putrescine, resulting in the formation of spermidine and 5′-methylthioadenosine (MTA). Putrescine is provided by another key enzyme, ornithine decarboxylase (ODC/*odc-1*) (Figure 6C). Interestingly while the abundance of downstream metabolites of the transmethylation pathway, S-adenosyl homocysteine (SAH) and cystathionine, was not altered in *sin-3* mutants (Figure 6B, Table S5 and Figure S4E), both dcSAM and MTA were strongly reduced (Table S5 and Figure S4E). Levels of neither arginine nor ornithine, two precursors of putrescine, nor putrescine itself was altered in s*in-3* mutants (Figure 6B, Table S5 and Figure S4C), while we detected a large increase in both spermidine and N-Acetyl-spermine, the degradation products of spermine and spermidine, respectively (Figure 6C, Table S5 and Figure S4E). Spermine could not be detected in our analysis. Together, these results suggest that in the absence of SIN-3, the metabolic flux is shifted from transmethylation to polyamine biosynthesis. Consistent with a shift in metabolic flux toward the synthesis of polyamines, the abundance of N-acetylputrescine, a degradation product of putrescine, was unaltered in our analyses (Table S5 and Figure S4D). The abundance of another polyamine, cadaverine, was also increased in *sin-3* mutants (Figure 6D, Table S5 and Figure S4E). Cadaverine is produced by the decarboxylation of lysine, also more abundant in *sin-3* mutants (Figure 6D, Table S5 and Figure S4C). No dedicated lysine decarboxylase has been annotated in *C. elegans* to date, but *odc-1* could potentially fulfill this function (Schaeffer and Donatelli, 1990).

Another lysine catabolite whose abundance increased in *sin-3* mutants is saccharopine, while aminoadipic acid (L-2-Aminoadipate), the metabolite further downstream, showed a slight but not significant increase (Figure 6D, Table S5 and Figure S4C). Because lysine degradation via the formation of saccharopine is confined to the mitochondria (Leandro and Houten, 2020), a potential blockage in its degradation due to defective mitochondria could increase its abundance (Figure 6D).

### Transcriptomics analysis reveals deregulation of metabolic pathways

The above metabolic analysis was carried out on *sin-3(tm1276)* mutant animals. Close inspection of the previously published list of genes misregulated in these mutants animals (Beurton et al., 2019) identified candidates whose misexpression correlates with the observed metabolic changes (Figure 6C). The ornithine decarboxylase *odc-1*, which converts ornithine to putrescine (Macrae et al., 1995), was strongly upregulated in *sin-3(tm1276)* mutants (log2FC 1.15, p = 4.09e-06), as was the spermidine synthase *spds-1* (log2FC 2.29, p = 2.88e-19). Regulation of polyamine synthesis is mainly achieved by controlling the activity of ornithine decarboxylase through its inhibitor antizyme, the binding of which disrupts ODC enzymatic activity and targets it for ubiquitin-independent degradation (Coffino, 2001). Significantly, the *C. elegans* homologue of antizyme, *oaz-1*, was significantly downregulated (log2FC −0.86, p = 4.29e-32), potentially leading to increased ODC-1 activity and spermidine biosynthesis in these mutants. Because *odc-1* may also decarboxylate lysine to produce cadaverin (Schaeffer and Donatelli, 1990), its increased activity could contribute to the observed increase in cadaverin levels. The polyamine oxidase (PAO) *hpo-15* is instead upregulated in *sin-3(tm1276)* mutants (log2FC 1.74, p = 6.56e-08). N-acetylspermine and N-acetylspermidine, metabolites of spermine and spermidine, respectively, are used as substrates by PAO to produce H_2_O_2_ and either spermidine or putrescine, depending on the starting substrate. An increased *hpo-15* expression may therefore also contribute to both the increase in spermidine levels and increased sensitivity to oxidative stress in *sin-3* mutant animals, as observed in mammalian cells (Jeong et al., 2022). Interestingly, spermidine serves as the sole biosynthetic precursor for hypusination, a post-translational modification that is an integral component of eukaryotic translation initiation factor 5A (eIF5A)(Park et al., 2010). Genes encoding the two enzymes required for hypusination, the deoxyhypusine synthase *dhps-1* and the deoxyhypusine hydroxylase *dohh-1*, were both upregulated in *sin-3(tm1276)* mutants (log2FC 0.7390, p = 7.89e-27 and 0.44, p = 2.60e-18 respectively). Of these genes only *oaz-1* and *hpo-15* are included in the list of genes commonly misregulated in the two *sin-3* alleles (Table 1), most likely reflecting experimental differences between the two transcriptomics analyses (Caron et al., 2023). Altogether our data is consistent with deregulating of specific metabolic genes in *sin-3(tm1276)* mutants contributing to the observed changes in polyamine flux. Whether these changes are a cause or consequence of the observed changes in mitochondrial dynamics and respiration remains to be established.

## Discussion

The SIN3 corepressor has been linked to mitochondrial function in several models, but how its loss alters mitochondrial dynamics and metabolism in the context of an entire organism has not been explored. Here, using two different *sin-3* mutant alleles, we show that in *C. elegans* adult animals partial or complete loss of *sin-3* function results in increased expression of genes encoded by the mitochondrial genome and an overall decrease in the expression of nuclear encoded mitochondrial genes, potentially leading to nuclear-mitochondrial imbalance. We show that the altered expression of mitochondrial genes is associated with increased mitochondrial fragmentation in all tissues examined, increased oxygen consumption, and metabolic changes, including a pronounced shift in metabolic flux from methionine to polyamine biosynthesis. Together our data identify SIN-3 as a key regulator of mitochondrial dynamics and polyamine homeostasis.

Knock-down of SIN3 in different systems results in strikingly similar phenotypes. In yeast, Drosophila cultured cells and *C. elegans,* reduced SIN3 activity is associated with lower ATP levels (Barnes et al., 2010; Pandey et al., 2018), and increased oxidative stress in both Drosophila and *C. elegans* adult animals (Barnes et al., 2014; Sharma et al., 2018). Deregulation of mitochondrial genes was reported following either knock-down or overexpression of SIN3 in cultured Drosophila cells (Barnes et al., 2010; Pile et al., 2003), or conditional *mSin3A* knock-out in primary cell culture (Dannenberg et al., 2005). Our transcriptomics analysis on *sin-3(tm1276)* and *sin-3(syb2178)* mutant animals revealed that expression of respiratory complex I subunits encoded by the mitochondrial genome is strongly increased in both mutants, while expression of nuclear encoded subunits of additional complexes is downregulated. The expression of additional genes with important roles in mitochondrial function, including mitochondrial ribosomal proteins and the *tomm-1* translocase is also significantly decreased in *sin-3* mutants. Importantly, SIN-3 binding on a number of downregulated genes suggests that SIN-3 may positively regulates their expression, as observed for SIN3 in other contexts (Caron et al., 2023; Baymaz et al., 2015; van Oevelen et al., 2010; Zhu et al., 2018).

The nucleus and mitochondria constantly communicate to adjust their activities in order to ensure cellular homeostasis (Ryan and Hoogenraad, 2007), and reducing the expression of a single major ribosomal nuclear protein (MRP) is sufficient to induce a stoichiometric imbalance between nuclear and mitochondrial-DNA encoded ETC subunits, referred to as mitonuclear protein imbalance (Houtkooper 2013). Mitonuclear imbalance leads to mitochondrial stress and activates the UPR^mt^ (Melber and Haynes, 2018). Interestingly, in *C. elegans,* the histone deacetylase *hda-1*, a conserved component of SIN3 complexes (Beurton et al., 2019; Caron et al., 2023), was shown to be required for this stress response (Shao et al., 2020). We observed that expression of the UPR^mt^ reporter *hsp-6* was dampened following induction of mitochondrial stress in *sin-3* knockdown animals, but only in old adults. Therefore, it is likely that HDA-1 acts in a context other than SIN-3 in the UPR^mt^, most likely the NuRD histone deacetylase complex (Zhu et al., 2020).

One of the most striking phenotypes we observed is that either reduced SIN-3 function or its complete absence results in a dramatic increase in mitochondrial fragmentation, observed by TEM in all tissues examined, including muscle, intestine, pharynx, hypodermis and germ cells, and through live imaging in muscle cells. The ability to transition between fission/fusion states is essential for mitochondrial function in cellular bioenergetics, regulation of intracellular Ca^2+^, and cellular stress responses (Wai and Langer, 2016). Fusion allows damaged mitochondria to mitigate stress by mixing contents, while fission contributes to quality control of damaged mitochondria by budding off deteriorating components for targeted breakdown via autophagy or mitophagy. The increased fragmentation we observe is consistent with a defect in the maintenance of mitochondrial homeostasis. We also observed examples of enlarged mitochondria, another hallmark of mitochondrial damage (Vincent et al., 2016), and the accumulation of electron-dense material in mitochondria that may represent iron deposits or misfolded proteins. Similar aggregates were observed by TEM in *C. elegans* following acute heat stress (Chen et al., 2021), in sperm-derived mitochondria before their elimination by mitophagy (Al Rawi et al., 2011; Al Rawi et al., 2011; Wang et al., 2016), and in the mitochondria of worms containing mutations in *eat-3/*OPA1 and *fzo-1/*mfn1 fusion genes (Byrne et al., 2019), supporting a link between their formation and mitochondrial morphological defects. Alternatively, these aggregates may represent iron deposits such as those observed in the mitochondria of a mouse model of Friedrich ataxia (Puccio et al., 2001; Whitnall et al., 2012) and congenital sideroblastic anemia (Ducamp and Fleming, 2019). Interestingly mitochondrial iron overload promotes ROS production, as observed in *sin-3* mutants (Pandey et al., 2018), and is associated with mitochondrial dysfunction and oxidative damage (Ward and Cloonan, 2019). Regardless of the underlying cause, disruption of mitochondria homeostasis, as observed in *sin-3* mutant animals, is likely to have major consequences for mitochondrial health and function.

The rate of oxygen consumption is another measure of mitochondrial activity, and we observed significant differences in respiration in *sin-3(tm1276)* mutants. While neither basal nor maximal oxygen consumption rates were affected in mutant young adults, both were significantly elevated in older mutants compared to wild type. Conversely, mitochondrial spare respiratory capacity was reduced in these mutants. Spare respiratory capacity can be viewed as a determinant of mitochondrial fitness and correlates with level of mitochondrial plasticity, allowing bioenergetic adaptability in response to pathopsysiological stress conditions (Marchetti et al., 2020). Its decrease in older *sin-3(tm1276)* mutants is consistent with mitochondrial dysfunction in these animals.

Increased basal and maximal oxygen consumption in old *sin-3(tm1276)* animals contrasts with wildtype, in which oxygen consumption has a tendency to decrease with age (Huang and Lin, 2018; Koopman et al., 2016). Interestingly, a similar increase during aging was observed in worms lacking the mitochondrial prohibitin (PHB) complex subunits *phb-1* and *phb-2* (de la Cruz-Ruiz et al., 2021). Like *sin-3* mutants, prohibitin mutants are also short-lived and show severely fragmented mitochondria (Artal-Sanz and Tavernarakis, 2009; Artal-Sanz et al., 2003). PHB plays a prominent role in the response to mitochondrial stress, quality control, biogenesis, and degradation (Hernando-Rodríguez and Artal-Sanz, 2018), and its expression is induced by metabolic stress resulting from mito-nuclear imbalance, but not other cellular stresses (Yoneda et al., 2004). It was suggested that PHB may act as chaperone in respiratory complex assembly (Nijtmans, 2000), with its loss affecting complex integrity. Likewise, imbalance between nuclear- and mitochondrial-encoded subunits in *sin-3* mutants could result in defects in respiratory complex assembly, proton leakage, and dissipation of the proton gradient. In this context, increased oxygen consumption may reflect compensatory activity from properly assembled supercomplexes, or mitochondrial adaptation resulting in an increase in mitochondrial mass (Bennett et al., 2022).

Consistent with mitochondrial fragmentation being generally associated with metabolic dysfunction and disease (Wai and Langer, 2016), metabolomic analysis revealed dramatic changes in the levels of several metabolites in *sin-3(tm1276)* mutants. One of the most striking differences we observed is an increase in the polyamines spermine and cadaverine. Polyamine biosynthesis is tightly controlled, and changes in polyamine levels can have dramatic consequences on physiology, with established links between polyamine metabolism and human diseases including cancer and diabetes (Madeo et al., 2018). Spermine is a potent free radical scavenger and an important antioxidant (Murray Stewart et al., 2018; Pegg, 2013), and in *C. elegans,* spermidine has been shown to inhibit neurodegeneration and have pro-longevity effects (Yang et al., 2020b). However, spermidine or spermine in excess can also have deleterious consequences on animal physiology (Kumar et al., 2022; Nakanishi and Cleveland, 2021; Pegg, 2013; Til et al., 1997), and SIN-3 inactivation or knock-down in *C. elegans* is associated with sterility, decreased longevity, and increased oxidative stress (Sharma et al., 2018). Our expression profiling suggests that decreased expression of the *oaz-1* antizyme, a critical regulator of polyamine biosynthesis (Coffino, 2001), may be a major contributor to the effect of SIN3 knock-down on polyamine homeostasis.

Among a broad range of functions, spermidine serves as the sole biosynthetic precursor for hypusination, a post-translational modification that is an integral component of eukaryotic translation initiation factor 5A (eIF5A) (Barba-Aliaga and Alepuz, 2022). Polyamine levels are elevated in most cancers, and hypusinated eIF5A is a critical regulator of cell growth (Nakanishi and Cleveland, 2016). More rescently, hypusinated eIF5A was shown to promote expression of mitochondrial proteins in macrophages (Puleston et al., 2019), suggesting that it may also alter mitochondrial activity in both normal and cancer cells. This raises the intriguing possibility that increased spermidine biosynthesis and eIF5A hypusylation in *sin-3* mutants may be part of a mechanism to increase expression of mitochondrial proteins in response to mitochondrial dysfunction.

Further evidence for a shift in metabolic flux towards polyamine biosynthesis comes from the analysis of downstream metabolites in the transmethylation pathway. Conversion of SAM to dcSAM provides necessary amino-propyl groups to sustain polyamine synthesis (Nakanishi and Cleveland, 2021). The methionine salvage pathway (MSP) recycles one carbon unit lost during polyamine synthesis back to the methionine cycle for SAM replenishment. In addition to showing a strong decrease in dcSAM, and a smaller but significant decrease in SAM, *sin-3* mutants also show reduced activity of the salvage pathway, as illustrated by a decrease in levels of the MSP intermediate methyladenosine (Figure 6C). Interestingly, in Prostate cancer (PCa) cells, which have an intrinsically high polyamine metabolic flux and therefore rely heavily on the methionine salvage pathway, MSP inhibition while maintaining high polyamine flux was shown to be an effective cancer therapy by blocking the ability of the cell to mitigate this stress, leading to cell death (Affronti et al., 2020). In other tumor cell types, by contrast, defects in the MSP were found to increase polyamine levels, highlighting how the stress that is generated by metabolic perturbations is often context-dependent (Nakanishi and Cleveland, 2021). Our study may provide novel insight on the interconnected links between SIN3, methionine metabolism, polyamine biosynthesis, and mitochondrial homeostasis, and point to new lines of investigation.

## Materials and Methods

### Strains and culture

Strains were cultured on Nematode Growth Medium (NGM) agarose plates with *Escherichia coli* OP50 and incubated at 20°C. The following strains were used (*created for this study, (** from (Beurton et al., 2019);***from (Caron et al., 2023): wild type N2 Bristol, **PFR590 *sin-3(tm1276*) I, ***PHX2172 *sin-3*(syb2172)/hT2[bli-4(e937) let-?(q782) qIs48] I, KAG420 kagIs4[gfp::*myo-3*]V (Mergoud Dit Lamarche et al., 2018); *PFR750 *sin-3(tm1276)* I, kagIs4[*gfp*::*myo-3*]V; EN7714 krSi134[*myo-3p*::*tomm-20*Nter::wrmScarlet] (Roy et al., 2022); PFR754 *sin-3(tm1276)* I; krSi134[*myo-3*p::*tomm-20*Nter::wrmScarlet]; EGD629 egxSi155[*mex-5p::tomm-20*::mKate2::*pie-1* 3’UTR + *unc-119*(+)] II; PFR758 *sin-3(tm1276)*, egxSi155[*mex-5p::tomm-20*::mKate2::pie-1 3’UTR + *unc-119*(+)]II; *PFR669 *sin-3:*:mCherry(*syb521*); oxIs279[*pie-1*p::GFP::H2B + *unc-119*(+)]II; SJ4100 *zcIs13* [*hsp-6p*::GFP + *lin-15*(+)]. PFR750 and PFR754 were obtained by crossing *sin-3(tm1276)* hermaphrodites with GFP::*myo-3* or *myo-3*p::tomm-20Nter::wrmScarlet males, respectively.

### Transmission electron microscopy

Worms (young adults) were fixed by HPF with EMPACT-2 (Leica Microsystems) and then freeze substituted in anhydrous acetone containing 1% OsO4, 0.5% glutaraldehyde and 0.25% uranyl acetate for 60 h in a FS system (AFS-2, Leica Microsystems). Larvae were embedded in Epon-Araldite mix (EMS hard formula). To gain better anteroposterior orientation and sectioning, adhesive frames were used (11560294 GENE-FRAME 65 μl, Thermo Fisher Scientific) for flat-embedding, as previously described (Kolotuev et al., 2012). Ultrathin sections were cut on an ultramicrotome (UC7; Leica Microsystems) and collected on formvar-coated slot grids (FCF2010-CU, EMS). Each larva was sectioned in five different places with ≥10 μm between each grid to ensure that different cells were observed. Each grid contained at least 5-10 consecutive sections of 70 nm. TEM grids were observed using a JEM-1400 transmission electron microscope (JEOL) operated at 120 kV, equipped with a Gatan Orius SC200 camera (Gatan) and piloted by the Digital Micrograph program.

### Quantification of muscle mitochondria and imaging of myofilament structure

For mitochondria and myofilament observation in muscle cells, staged worms were mounted on agarose pads in M9 solution containing 10% sodium azide. Muscles in the posterior part of worms were imaged. Z-stack images were acquired using a Zeiss LSM980 inverted confocal microscope with 63× oil immersion objective. Z-stacks were acquired every 0.27μm to image mitochondria and every 0.23μm to image myofilaments. Images shown are projections using max intensity method in Fiji. Quantification of mitochondrial morphology in body wall muscle cells was performed in double blind experiments according to the following criteria: cells containing long interconnected mitochondrial networks were classified as tubular, cells containing a combination of interconnected mitochondrial networks along with smaller fragmented mitochondria were classified as intermediate, and cells with sparse small round mitochondria were classified as fragmented (Regmi et al., 2014).

### Activation of UPR^mt^ by RNAi treatment

RNAi bacteria were induced by directly adding 1 mM IPTG directly onto the bacteria lawn 2 hours before the transfer of worms (Shamalnasab et al., 2017). Transgenic animals expressing *hsp-6::*GFP were synchronized by bleaching, and the L1 stage animals obtained transferred to empty vector HT115 or *sin-3(*RNAi) 35mm plates, and allowed to develop to egg-laying adults (P0). 4-6 P0 adults from each RNAi treatment were then individually transferred onto: 1) empty vector control; 2) single RNAi (*sin-3*, *nuo-4*, *mrps-5)* or double RNAi plates (sin-3/*nuo-4*, *sin-3*/*mrps-5)* to induce mitochondrial stress in F1 progeny. *sin-3* RNAi treatment was started in P0 animals to ensure depletion of maternal SIN-3 protein in scored animals. GFP expression was assessed in F1 progeny 5 days after egg lay. For imaging, worms were immobilized using 2.5mM Levamisole in M9 buffer (22mM KH2PO4, 42mM Na2HPO4, 86mM NaCl, 1mM MgSO4). Images were acquired with a Nikon AZ100M microscope.

### Oxygen consumption measurements

Worm oxygen consumption was measured using the Agilent Seahorse XFp Analyzer. Animals were synchronized by allowing gravid mothers to lay eggs for 2-3 hours before removing them from plate. Worms (20-30 per well) at the young adult stage or at day 6 of adulthood were transferred into M9-filled Seahorse plates. Working solutions were diluted in M9 at the following final concentrations: Carbonyl cyanide 4-(trifluoromethoxy) phenylhydrazone (FCCP) (Sigma-Aldrich) 250 μM, NaN_3_ (Sigma-Aldrich) 400 mM. Oxygen Consumption Rate (OCR) measurements were repeated 8 times in basal conditions, 10 times after FCPP injection (for maximal respiration), and 4 times after NaN_3_ injection (no mitochondrial respiration). Each cycle consisted of 3 min mix, 30 second wait, and 3 min measure. Values were normalized per worm. Three independent assays were carried out and the combined data was analyzed by Unpaired t-test, or, when appropriate, Mann-Whitney test, using GraphPad Prism software.

### Western blot analysis

Young adult worms were collected in M9 buffer, washed 3 times, pelleted and frozen in dry ice. Pellets were resuspended in TNET buffer (50 mM Tris·HCl (pH 8), 300 mM NaCl, 1 mM EDTA, 0,5% Triton X-100 and cOmplete™ Protease Inhibitor Cocktail [Merck # 11697498001]) and lysed with zirconium beads [Lysing Matrix Y, MP Biomedicals #116960050] using a Precellys24 homogenizer [Ozyme] with the following parameters : 6000 rpm 2×20sec. Homogenates were centrifuged and supernatants aliquoted and frozen at −80◦C. Total protein amount was quantified by the Bradford assay [Bio-Rad]. 10,20,30,40,50 and 80 ug of protein extracts were loaded either on 12% NuPage Novex or 12 % SDS PAGE gels for western blot analysis. After transfer, membranes were incubated overnight with total OXPHOS Rodent WB Antibody Cocktail [Abcam #ab110413] diluted at 1:500 or anti-alpha-tubulin [Abcam #ab18251] diluted at 1:5,000. The next day, membranes were incubated for 1 hour with goat anti-rabbit DyLight™ 800 [Invitrogen # SA5-10036] and IRDye® 680RD goat anti-mouse [LiCOR #926-68070] diluted at 1:10,000. Aquisition was performed on a ChemiDoc MP apparatus [Bio-Rad]. Quantification was carried out using Image J, and each protein signal was normalized to the level of tubulin. Two to three independent biological replicates and three to four technical replicates were used for quantification.

### Metabolite extraction and profiling

Embryos derived from bleaching were transferred onto 90 mm plates seeded with 0P50 bacteria and left to develop to the young adult stage. Animals were collected in M9 medium, washed 3 times in 10 ml M9, and pellets quick frozen in liquid nitrogen (Spanier et al., 2021) prior to processing. 8 replicas, each containing approximately 2000 worms, were processed per genotype. All chemicals and solvents were of LC-MS or analytical grade. *C. elegans* samples were thawed on ice and 1 ml of ice-cold H_2_O/MeOH/CHCl_3_ (1/3/1, v/v/v). After suspension in solvent, *C. elegans* animals were transferred to 2 ml MN Bead Tubes Type A (Macherey-Nagel, Düren, Germany) and lysed using a Precellys Bead Beating system with an additional Cryolys cooling module (Bertin Instruments, Montigny-le-Bretonneux, France). After lysis, samples were incubated for 10 minutes in an ice-cold ultrasonic bath, followed by centrifugation at 4°C and 13,000 rpm for 15 minutes. The supernatant was transferred to a fresh reaction tube and evaporated to dryness using a centrifugal evaporator. Proteins were extracted from residue debris pellets and quantified using a BCA kit (Sigma-Aldrich, Taufkirchen, Germany). Metabolite profiling was performed using a Sciex ExionLC AD coupled to a Sciex ZenoTOF 7600 under the control of Sciex OS 3.0 (Sciex, Darmstadt, Germany). Separation was achieved on an Agilent InfinityLab Poroshell 120 HILIC-Z column (2.1 mm x 150 mm, 2.7 µm particle size, PEEK-lined) (Agilent Technologies, Waldbronn, Germany). Different eluents and gradients were applied for positive and negative ionization modes. In positive ionization mode, eluent A consisted of 100% H_2_O + 10 mM ammonium formate / 0.1% formic acid, and eluent B consisted of 10% H_2_O / 90% ACN + 10 mM ammonium formate / 0.1% formic acid. In negative ionization mode, eluent A consisted of 100% H_2_O + 10 mM ammonium acetate / 2.5 µM medronic acid, pH = 9, and eluent A consisted of 15% H_2_O / 85% ACN + 10 mm ammonium acetate / 2.5 µM medronic acid, pH = 9. The column temperature was set to 25°C and 50°C for positive and negative ionization modes, respectively, and the flow rate was 0.25 mL/min in both cases. Gradients for both ionization modes are summarized in Table S2. Dried samples were re-dissolved in 50 µL H_2_O/ACN (1/3, v/v), and 40 µL were transferred to an autosampler vial and 10 µL to a pooled QC sample. Autosampler temperature was set to 5°C and 5 µl were injected for analysis. In MS^1,^ ions in the m/z range 70 to 1500 were accumulated for 0.1s, and information-dependent acquisition (IDA) of MS^2^ was used with a maximum number of 6 candidate ions and a collision energy of 35 eV with a spread of 15 eV. Accumulation time for MS2 was set to 0.025s yielding a total cycle time of 0.299s. ZenoTrapping was enabled with a value of 80000. QC samples were used for conditioning the column and were also injected every ten samples. Automatic calibration of the mass spectrometer in MS^1^ and MS^2^ mode was performed every five injections using the ESI positive Calibration Solution for the Sciex X500 system or the ESI negative Calibration Solution for the Sciex X500 system (Sciex, Darmstadt, Germany).

### Metabolite data analysis

Data analysis was performed in a targeted fashion for identified metabolites (see Tables S4 and S5). Metabolites were identified by comparison to in-house reference standards, publicly available reference spectra, and manual interpretation of fragmentation spectra. Data analysis was performed in Sciex OS 3.0.0.3339 (Sciex, Darmstadt, Germany). Peaks for all metabolites indicated in Tables S4 and S5 were integrated with an XIC width of 0.02 Da and a Gaussian smooth width of 3 points using the MQ4 peak-picking algorithm. Peak areas were exported to a .txt file and normalized according to the protein content of the respective sample. All further processing was performed in R 4.2.1 within RStudio using the following packages: tidyverse (1.3.2), readxl (1.4.1), ggsignif (0.6.4), gghalves (0.1.4), scales (1.2.1) and viridis (0.6.2). Data were plotted using ggplot with gghalves using half box- and half dot-plots. Significance was tested using a Welch test within ggsignif.

## Acknowledgments

FP was supported by ANR (Agence Nationale de la Recherche) grant N° 19-CE12-0025-01 and the Centre National de la Recherche Scientifique. MAS was funded through grants from the Ministerio de Ciencia, Innovación y Universidades, the Agencia Estatal de Investigación (AEI) PID 2019-104145GB-I00. GM laboratory receives institutional funding from the Centre National De La Recherche Scientifique and the Université de Rennes. TEM imaging was performed at the Microscopy Rennes imaging Center (Biosit, Rennes, France), a member of the national infrastructure France-BioImaging supported by the French National Research Agency (ANR-10-INBS-04).

## Author contributions

PF, FP, GM, MAS and MW conceived and designed experiments, MG, MJRP, ON and MW and CB performed experiments and analyzed the data, FP, MAS and MW wrote the paper.

## References

Affronti, H.C., A.M. Rowsam, A.J. Pellerite, S.R. Rosario, M.D. Long, J.J. Jacobi, A. Bianchi-Smiraglia, C.S. Boerlin, B.M. Gillard, E. Karasik, B.A. Foster, M. Moser, J.H. Wilton, K. Attwood, M.A. Nikiforov, G. Azabdaftari, R. Pili, J.G. Phillips, R.A. Casero, and D.J. Smiraglia. 2020. Pharmacological polyamine catabolism upregulation with methionine salvage pathway inhibition as an effective prostate cancer therapy. Nat Commun. 11:52. doi:10.1038/s41467-019-13950-4.

Al Rawi, S., S. Louvet-Vallée, A. Djeddi, M. Sachse, E. Culetto, C. Hajjar, L. Boyd, R. Legouis, and V. Galy. 2011. Postfertilization Autophagy of Sperm Organelles Prevents Paternal Mitochondrial DNA Transmission. Science. 334:1144–1147. doi:10.1126/science.1211878.

Ali, A.T., L. Boehme, G. Carbajosa, V.C. Seitan, K.S. Small, and A. Hodgkinson. 2019. Nuclear genetic regulation of the human mitochondrial transcriptome. Elife. 8:e41927. doi:10.7554/eLife.41927.

Anderson, N.S., and C.M. Haynes. 2020. Folding the Mitochondrial UPR into the Integrated Stress Response. Trends in Cell Biology. 30:428–439. doi:10.1016/j.tcb.2020.03.001.

Artal-Sanz, M., and N. Tavernarakis. 2009. Prohibitin couples diapause signalling to mitochondrial metabolism during ageing in C. elegans. Nature. 461:793–797. doi:10.1038/nature08466.

Artal-Sanz, M., W.Y. Tsang, E.M. Willems, L.A. Grivell, B.D. Lemire, H. van der Spek, and L.G.J. Nijtmans. 2003. The mitochondrial prohibitin complex is essential for embryonic viability and germline function in Caenorhabditis elegans. J Biol Chem. 278:32091–32099. doi:10.1074/jbc.M304877200.

Balasubramanian, M., A.J.M. Dingemans, S. Albaba, R. Richardson, T.M. Yates, H. Cox, S. Douzgou, R. Armstrong, F.H. Sansbury, K.B. Burke, A.E. Fry, N. Ragge, S. Sharif, A. Foster, A. De Sandre-Giovannoli, S. Elouej, P. Vasudevan, S. Mansour, K. Wilson, H. Stewart, S. Heide, C. Nava, B. Keren, S. Demirdas, A.S. Brooks, M. Vincent, B. Isidor, S. Küry, M. Schouten, E. Leenders, W.K. Chung, A. van Haeringen, T. Scheffner, F.-G. Debray, S.M. White, M.I.V. Palafoll, R. Pfundt, R. Newbury-Ecob, and T. Kleefstra. 2021. Comprehensive study of 28 individuals with SIN3A-related disorder underscoring the associated mild cognitive and distinctive facial phenotype. Eur J Hum Genet. 29:625–636. doi:10.1038/s41431-020-00769-7.

Bansal, N., G. David, E. Farias, and S. Waxman. 2016. Emerging Roles of Epigenetic Regulator Sin3 in Cancer. Adv. Cancer Res. 130:113–135. doi:10.1016/bs.acr.2016.01.006.

Bansal, N., K. Petrie, R. Christova, C.-Y. Chung, B.A. Leibovitch, L. Howell, V. Gil, Y. Sbirkov, E. Lee, J. Wexler, E.V. Ariztia, R. Sharma, J. Zhu, E. Bernstein, M.-M. Zhou, A. Zelent, E. Farias, and S. Waxman. 2015. Targeting the SIN3A-PF1 interaction inhibits epithelial to mesenchymal transition and maintenance of a stem cell phenotype in triple negative breast cancer. Oncotarget. 6:34087–34105. doi:10.18632/oncotarget.6048.

Barba-Aliaga, M., and P. Alepuz. 2022. Role of eIF5A in Mitochondrial Function. IJMS. 23:1284. doi:10.3390/ijms23031284.

Barnes, V.L., A. Bhat, A. Unnikrishnan, A.R. Heydari, R. Arking, and L.A. Pile. 2014. SIN3 is critical for stress resistance and modulates adult lifespan. Aging. 6:645–660. doi:10.18632/aging.100684.

Barnes, V.L., B.S. Strunk, I. Lee, M. Hüttemann, and L.A. Pile. 2010. Loss of the SIN3 transcriptional corepressor results in aberrant mitochondrial function. BMC Biochem. 11:26. doi:10.1186/1471-2091-11-26.

Barshad, G., A. Blumberg, T. Cohen, and D. Mishmar. 2018. Human primitive brain displays negative mitochondrial-nuclear expression correlation of respiratory genes. Genome Res. 28:952–967. doi:10.1101/gr.226324.117.

Baymaz, H.I., I.D. Karemaker, and M. Vermeulen. 2015. Perspective on unraveling the versatility of ‘co-repressor’ complexes. Biochimica et Biophysica Acta (BBA) - Gene Regulatory Mechanisms. 1849:1051–1056. doi:10.1016/j.bbagrm.2015.06.012.

Bennett, C.F., P. Latorre-Muro, and P. Puigserver. 2022. Mechanisms of mitochondrial respiratory adaptation. Nat Rev Mol Cell Biol. 23:817–835. doi:10.1038/s41580-022-00506-6.

Berry, B.J., E. Mjelde, F. Carreno, K. Gilham, E.J. Hanson, E. Na, and M. Kaeberlein. 2022. Preservation of Mitochondrial Membrane Potential is Necessary for Lifespan Extension from Dietary Restriction. Physiology.

Bertholet, A.M., E.T. Chouchani, L. Kazak, A. Angelin, A. Fedorenko, J.Z. Long, S. Vidoni, R. Garrity, J. Cho, N. Terada, D.C. Wallace, B.M. Spiegelman, and Y. Kirichok. 2019. H+ transport is an integral function of the mitochondrial ADP/ATP carrier. Nature. 571:515–520. doi:10.1038/s41586-019-1400-3.

Beurton, F., P. Stempor, M. Caron, A. Appert, Y. Dong, R.A.-J. Chen, D. Cluet, Y. Couté, M. Herbette, N. Huang, H. Polveche, M. Spichty, C. Bedet, J. Ahringer, and F. Palladino. 2019. Physical and functional interaction between SET1/COMPASS complex component CFP-1 and a Sin3S HDAC complex in C. elegans. Nucleic Acids Res. 47:11164–11180. doi:10.1093/nar/gkz880.

Byrne, J.J., M.S. Soh, G. Chandhok, T. Vijayaraghavan, J.-S. Teoh, S. Crawford, A.E. Cobham, N.M.B. Yapa, C.K. Mirth, and B. Neumann. 2019. Disruption of mitochondrial dynamics affects behaviour and lifespan in Caenorhabditis elegans. Cell. Mol. Life Sci. 76:1967–1985. doi:10.1007/s00018-019-03024-5.

Caron, M., V. Robert, L. Gely, A. Adrait, V. Pakulska, Y. Couté, M. Chevalier, C.G. Riedel, C. Bedet, and F. Palladino. 2023. SIN3 acts in distinct complexes to regulate the germline transcriptional program in *C. elegans*. Genetics.

Chaubal, A., and L.A. Pile. 2018. Same agent, different messages: insight into transcriptional regulation by SIN3 isoforms. Epigenetics Chromatin. 11:17. doi:10.1186/s13072-018-0188-y.

Chen, H., and D.C. Chan. 2009. Mitochondrial dynamics-fusion, fission, movement, and mitophagy-in neurodegenerative diseases. Human Molecular Genetics. 18:R169–R176. doi:10.1093/hmg/ddp326.

Chen, Y., R. Leboutet, C. Largeau, S. Zentout, C. Lefebvre, A. Delahodde, E. Culetto, and R. Legouis. 2021. Autophagy facilitates mitochondrial rebuilding after acute heat stress via a DRP-1– dependent process. Journal of Cell Biology. 220:e201909139. doi:10.1083/jcb.201909139.

Christian, C.J., and G.M. Benian. 2020. Animal models of sarcopenia. Aging Cell. 19. doi:10.1111/acel.13223.

Coffino, P. 2001. Regulation of cellular polyamines by antizyme. Nat Rev Mol Cell Biol. 2:188–194. doi:10.1038/35056508.

de la Cruz-Ruiz, P., B. Hernando-Rodríguez, M.M. Pérez-Jiménez, M.J. Rodríguez-Palero, M.D. Martínez-Bueno, A. Pla, R. Gatsi, and M. Artal-Sanz. 2021. Prohibitin depletion extends lifespan of a TORC2/SGK-1 mutant through autophagy and the mitochondrial UPR. Aging Cell. 20:e13359. doi:10.1111/acel.13359.

Dannenberg, J.-H., G. David, S. Zhong, J. van der Torre, W.H. Wong, and R.A. Depinho. 2005. mSin3A corepressor regulates diverse transcriptional networks governing normal and neoplastic growth and survival. Genes Dev. 19:1581–1595. doi:10.1101/gad.1286905.

Demine, Renard, and Arnould. 2019. Mitochondrial Uncoupling: A Key Controller of Biological Processes in Physiology and Diseases. Cells. 8:795. doi:10.3390/cells8080795.

DiMauro, S., and E.A. Schon. 2003. Mitochondrial Respiratory-Chain Diseases. N Engl J Med. 348:2656– 2668. doi:10.1056/NEJMra022567.

Ducamp, S., and M.D. Fleming. 2019. The molecular genetics of sideroblastic anemia. Blood. 133:59–69. doi:10.1182/blood-2018-08-815951.

Etheridge, T., M. Rahman, C.J. Gaffney, D. Shaw, F. Shephard, J. Magudia, D.E. Solomon, T. Milne, J. Blawzdziewicz, D. Constantin-Teodosiu, P.L. Greenhaff, S.A. Vanapalli, and N.J. Szewczyk. 2015. The integrin-adhesome is required to maintain muscle structure, mitochondrial ATP production, and movement forces in *Caenorhabditis elegans*. FASEB j. 29:1235–1246. doi:10.1096/fj.14-259119.

Farias, E.F., K. Petrie, B. Leibovitch, J. Murtagh, M.B. Chornet, T. Schenk, A. Zelent, and S. Waxman. 2010. Interference with Sin3 function induces epigenetic reprogramming and differentiation in breast cancer cells. Proc. Natl. Acad. Sci. U.S.A. 107:11811–11816. doi:10.1073/pnas.1006737107.

Fernandez-Vizarra, E., and M. Zeviani. 2021. Mitochondrial disorders of the OXPHOS system. FEBS Lett. 595:1062–1106. doi:10.1002/1873-3468.13995.

Gaffney, C.J., A. Pollard, T.F. Barratt, D. Constantin-Teodosiu, P.L. Greenhaff, and N.J. Szewczyk. 2018. Greater loss of mitochondrial function with ageing is associated with earlier onset of sarcopenia in C. elegans. Aging. 10:3382–3396. doi:10.18632/aging.101654.

Hanover, J.A., M.E. Forsythe, P.T. Hennessey, T.M. Brodigan, D.C. Love, G. Ashwell, and M. Krause. 2005. A Caenorhabditis elegans model of insulin resistance: altered macronutrient storage and dauer formation in an OGT-1 knockout. Proc Natl Acad Sci U S A. 102:11266–11271. doi:10.1073/pnas.0408771102.

Haeussler, S., and B. Conradt. 2022. Methods to Study the Mitochondrial Unfolded Protein Response (UPRmt) in Caenorhabditis elegans. *In* The Unfolded Protein Response. R. Pérez-Torrado, editor. Springer US, New York, NY. 249–259.

Hernando-Rodríguez, B., and M. Artal-Sanz. 2018. Mitochondrial Quality Control Mechanisms and the PHB (Prohibitin) Complex. Cells. 7:238. doi:10.3390/cells7120238.

Higashitani, A., M. Teranishi, Y. Nakagawa, Y. Itoh, S. Sudevan, N.J. Szewczyk, Y. Kubota, T. Abe, and T. Kobayashi. 2022. Increased mitochondrial Ca ^2+^ contributes to health decline with age and Duchene muscular dystrophy in *C. elegans*. Cell Biology.

Holdorf, A.D., D.P. Higgins, A.C. Hart, P.R. Boag, G.J. Pazour, A.J.M. Walhout, and A.K. Walker. 2020. WormCat: An Online Tool for Annotation and Visualization of *Caenorhabditis elegans* Genome-Scale Data. Genetics. 214:279–294. doi:10.1534/genetics.119.302919.

Houtkooper, R.H., L. Mouchiroud, D. Ryu, N. Moullan, E. Katsyuba, G. Knott, R.W. Williams, and J. Auwerx. 2013. Mitonuclear protein imbalance as a conserved longevity mechanism. Nature. 497:451–457. doi:10.1038/nature12188.

Huang, S.-H., and Y.-W. Lin. 2018. Bioenergetic Health Assessment of a Single Caenorhabditis elegans from Postembryonic Development to Aging Stages via Monitoring Changes in the Oxygen Consumption Rate within a Microfluidic Device. Sensors. 18:2453. doi:10.3390/s18082453.

Jeong, H.D., J.H. Kim, G.E. Kwon, and S.-T. Lee. 2022. Expression of Polyamine Oxidase in Fibroblasts Induces MMP-1 and Decreases the Integrity of Extracellular Matrix. IJMS. 23:10487. doi:10.3390/ijms231810487.

Kolotuev, I., D.J. Bumbarger, M. Labouesse, and Y. Schwab. 2012. Targeted Ultramicrotomy. In Methods in Cell Biology. Elsevier. 203–222.

Koopman, M., H. Michels, B.M. Dancy, R. Kamble, L. Mouchiroud, J. Auwerx, E.A.A. Nollen, and R.H. Houtkooper. 2016. A screening-based platform for the assessment of cellular respiration in Caenorhabditis elegans. Nat Protoc. 11:1798–1816. doi:10.1038/nprot.2016.106.

Kropp, P.A., J. Wu, M. Reidy, S. Shrestha, K. Rhodehouse, P. Rogers, M.N. Sack, and A. Golden. 2021. Allele-specific mitochondrial stress induced by Multiple Mitochondrial Dysfunctions Syndrome 1 pathogenic mutations modeled in Caenorhabditis elegans. PLoS Genet. 17:e1009771. doi:10.1371/journal.pgen.1009771.

Kumar, V., R.K. Mishra, D. Ghose, A. Kalita, P. Dhiman, A. Prakash, N. Thakur, G. Mitra, V.D. Chaudhari, A. Arora, and D. Dutta. 2022. Free spermidine evokes superoxide radicals that manifest toxicity. eLife. 11:e77704. doi:10.7554/eLife.77704.

Labrousse, A.M., M.D. Zappaterra, D.A. Rube, and A.M. van der Bliek. 1999. C. elegans Dynamin-Related Protein DRP-1 Controls Severing of the Mitochondrial Outer Membrane. Molecular Cell. 4:815–826. doi:10.1016/S1097-2765(00)80391-3.

Latypova, X., M. Vincent, A. Mollé, O.A. Adebambo, C. Fourgeux, T.N. Khan, A. Caro, M. Rosello, C. Orellana, D. Niyazov, D. Lederer, M. Deprez, Y. Capri, P. Kannu, A.C. Tabet, J. Levy, E. Aten, N. den Hollander, M. Splitt, J. Walia, L.L. Immken, P. Stankiewicz, K. McWalter, S. Suchy, R.J. Louie, S. Bell, R.E. Stevenson, J. Rousseau, C. Willem, C. Retiere, X.-J. Yang, P.M. Campeau, F. Martinez, J.A. Rosenfeld, C. Le Caignec, S. Küry, S. Mercier, K. Moradkhani, S. Conrad, T. Besnard, B. Cogné, N. Katsanis, S. Bézieau, J. Poschmann, E.E. Davis, and B. Isidor. 2021. Haploinsufficiency of the Sin3/HDAC corepressor complex member SIN3B causes a syndromic intellectual disability/autism spectrum disorder. The American Journal of Human Genetics. 108:929–941. doi:10.1016/j.ajhg.2021.03.017.

Leandro, J., and S.M. Houten. 2020. The lysine degradation pathway: Subcellular compartmentalization and enzyme deficiencies. Molecular Genetics and Metabolism. 131:14–22. doi:10.1016/j.ymgme.2020.07.010.

Lewis, M.J., J. Liu, E.F. Libby, M. Lee, N.P.S. Crawford, and D.R. Hurst. 2016. SIN3A and SIN3B differentially regulate breast cancer metastasis. Oncotarget. 7:78713–78725. doi:10.18632/oncotarget.12805.

Liang, V., M. Ullrich, H. Lam, Y.L. Chew, S. Banister, X. Song, T. Zaw, M. Kassiou, J. Götz, and H.R. Nicholas. 2014. Altered proteostasis in aging and heat shock response in C. elegans revealed by analysis of the global and de novo synthesized proteome. Cell. Mol. Life Sci. 71:3339–3361. doi:10.1007/s00018-014-1558-7.

Lin, Y.-F., and C.M. Haynes. 2016. Metabolism and the UPR mt. Molecular Cell. 61:677–682. doi:10.1016/j.molcel.2016.02.004.

Lionaki, E., I. Gkikas, I. Daskalaki, M.-K. Ioannidi, M.I. Klapa, and N. Tavernarakis. 2022. Mitochondrial protein import determines lifespan through metabolic reprogramming and de novo serine biosynthesis. Nat Commun. 13:651. doi:10.1038/s41467-022-28272-1.

Locasale, J.W. 2013. Serine, glycine and one-carbon units: cancer metabolism in full circle. Nat Rev Cancer. 13:572–583. doi:10.1038/nrc3557.

Luz, A.L., J.P. Rooney, L.L. Kubik, C.P. Gonzalez, D.H. Song, and J.N. Meyer. 2015. Mitochondrial Morphology and Fundamental Parameters of the Mitochondrial Respiratory Chain Are Altered in Caenorhabditis elegans Strains Deficient in Mitochondrial Dynamics and Homeostasis Processes. PLoS One. 10:e0130940. doi:10.1371/journal.pone.0130940.

Macrae, M., R.H. Plasterk, and P. Coffino. 1995. The ornithine decarboxylase gene of Caenorhabditis elegans: cloning, mapping and mutagenesis. Genetics. 140:517–525. doi:10.1093/genetics/140.2.517.

Madeo, F., T. Eisenberg, F. Pietrocola, and G. Kroemer. 2018. Spermidine in health and disease. Science. 359:eaan2788. doi:10.1126/science.aan2788.

Marchetti, P., Q. Fovez, N. Germain, R. Khamari, and J. Kluza. 2020. Mitochondrial spare respiratory capacity: Mechanisms, regulation, and significance in non-transformed and cancer cells. FASEB j. 34:13106–13124. doi:10.1096/fj.202000767R.

Matilainen, O., P.M. Quirós, and J. Auwerx. 2017. Mitochondria and Epigenetics - Crosstalk in Homeostasis and Stress. Trends Cell Biol. 27:453–463. doi:10.1016/j.tcb.2017.02.004.

Melber, A., and C.M. Haynes. 2018. UPRmt regulation and output: a stress response mediated by mitochondrial-nuclear communication. Cell Res. 28:281–295. doi:10.1038/cr.2018.16.

Mercer, T.R., S. Neph, M.E. Dinger, J. Crawford, M.A. Smith, A.-M.J. Shearwood, E. Haugen, C.P. Bracken, O. Rackham, J.A. Stamatoyannopoulos, A. Filipovska, and J.S. Mattick. 2011. The human mitochondrial transcriptome. Cell. 146:645–658. doi:10.1016/j.cell.2011.06.051.

Mergoud Dit Lamarche, A., L. Molin, L. Pierson, M.-C. Mariol, J.-L. Bessereau, K. Gieseler, and F. Solari. 2018. UNC-120/SRF independently controls muscle aging and lifespan in Caenorhabditis elegans. Aging Cell. 17:e12713. doi:10.1111/acel.12713.

Murray Stewart, T., T.T. Dunston, P.M. Woster, and R.A. Casero. 2018. Polyamine catabolism and oxidative damage. Journal of Biological Chemistry. 293:18736–18745. doi:10.1074/jbc.TM118.003337.

Nakanishi, S., and J.L. Cleveland. 2021. Polyamine Homeostasis in Development and Disease. Medical Sciences. 9:28. doi:10.3390/medsci9020028.

Nielsen, J. 2003. It Is All about MetabolicFluxes. J Bacteriol. 185:7031–7035. doi:10.1128/JB.185.24.7031-7035.2003.

Nijtmans, L.G.J. 2000. Prohibitins act as a membrane-bound chaperone for the stabilization of mitochondrial proteins. The EMBO Journal. 19:2444–2451. doi:10.1093/emboj/19.11.2444.

van Oevelen, C., C. Bowman, J. Pellegrino, P. Asp, J. Cheng, F. Parisi, M. Micsinai, Y. Kluger, A. Chu, A. Blais, G. David, and B.D. Dynlacht. 2010. The Mammalian Sin3 Proteins Are Required for Muscle Development and Sarcomere Specification. Molecular and Cellular Biology. 30:5686– 5697. doi:10.1128/MCB.00975-10.

Onishi, M., K. Yamano, M. Sato, N. Matsuda, and K. Okamoto. 2021. Molecular mechanisms and physiological functions of mitophagy. EMBO J. 40:e104705. doi:10.15252/embj.2020104705.

Pandey, R., M. Sharma, and D. Saluja. 2018. SIN-3 as a key determinant of lifespan and its sex dependent differential role on healthspan in Caenorhabditis elegans. Aging. 10:3910–3937. doi:10.18632/aging.101682.

Park, M.H., K. Nishimura, C.F. Zanelli, and S.R. Valentini. 2010. Functional significance of eIF5A and its hypusine modification in eukaryotes. Amino Acids. 38:491–500. doi:10.1007/s00726-009-0408-7.

Pegg, A.E. 2013. Toxicity of Polyamines and Their Metabolic Products. Chem. Res. Toxicol. 26:1782– 1800. doi:10.1021/tx400316s.

Pile, L.A., P.T. Spellman, R.J. Katzenberger, and D.A. Wassarman. 2003. The SIN3 Deacetylase Complex Represses Genes Encoding Mitochondrial Proteins. Journal of Biological Chemistry. 278:37840– 37848. doi:10.1074/jbc.M305996200.

Puccio, H., D. Simon, M. Cossée, P. Criqui-Filipe, F. Tiziano, J. Melki, C. Hindelang, R. Matyas, P. Rustin, and M. Koenig. 2001. Mouse models for Friedreich ataxia exhibit cardiomyopathy, sensory nerve defect and Fe-S enzyme deficiency followed by intramitochondrial iron deposits. Nat Genet. 27:181–186. doi:10.1038/84818.

Puleston, D.J., M.D. Buck, R.I. Klein Geltink, R.L. Kyle, G. Caputa, D. O’Sullivan, A.M. Cameron, A. Castoldi, Y. Musa, A.M. Kabat, Y. Zhang, L.J. Flachsmann, C.S. Field, A.E. Patterson, S. Scherer, F. Alfei, F. Baixauli, S.K. Austin, B. Kelly, M. Matsushita, J.D. Curtis, K.M. Grzes, M. Villa, M. Corrado, D.E. Sanin, J. Qiu, N. Pällman, K. Paz, M.E. Maccari, B.R. Blazar, G. Mittler, J.M. Buescher, D. Zehn, S. Rospert, E.J. Pearce, S. Balabanov, and E.L. Pearce. 2019. Polyamines and eIF5A Hypusination Modulate Mitochondrial Respiration and Macrophage Activation. Cell Metabolism. 30:352–363.e8. doi:10.1016/j.cmet.2019.05.003.

Quirós, P.M., A. Mottis, and J. Auwerx. 2016. Mitonuclear communication in homeostasis and stress. Nat Rev Mol Cell Biol. 17:213–226. doi:10.1038/nrm.2016.23.

Quirós, P.M., M.A. Prado, N. Zamboni, D. D’Amico, R.W. Williams, D. Finley, S.P. Gygi, and J. Auwerx. 2017. Multi-omics analysis identifies ATF4 as a key regulator of the mitochondrial stress response in mammals. J Cell Biol. 216:2027–2045. doi:10.1083/jcb.201702058.

Reeve, A.K., K.J. Krishnan, and D. Turnbull. 2008. Mitochondrial DNA Mutations in Disease, Aging, and Neurodegeneration. Annals of the New York Academy of Sciences. 1147:21–29. doi:10.1196/annals.1427.016.

Regmi, S.G., S.G. Rolland, and B. Conradt. 2014. Age-dependent changes in mitochondrial morphology and volume are not predictors of lifespan. Aging. 6:118–130. doi:10.18632/aging.100639.

Richter-Dennerlein, R., S. Oeljeklaus, I. Lorenzi, C. Ronsör, B. Bareth, A.B. Schendzielorz, C. Wang, B. Warscheid, P. Rehling, and S. Dennerlein. 2016. Mitochondrial Protein Synthesis Adapts to Influx of Nuclear-Encoded Protein. Cell. 167:471–483.e10. doi:10.1016/j.cell.2016.09.003.

Rolland, S., and B. Conradt. 2022. Genetic screen identifies non-mitochondrial proteins involved in the maintenance of mitochondrial homeostasis. MicroPubl Biol. 2022. doi:10.17912/micropub.biology.000562.

Rolland, S.G., S. Schneid, M. Schwarz, E. Rackles, C. Fischer, S. Haeussler, S.G. Regmi, A. Yeroslaviz, B. Habermann, D. Mokranjac, E. Lambie, and B. Conradt. 2019. Compromised Mitochondrial Protein Import Acts as a Signal for UPRmt. Cell Reports. 28:1659–1669.e5. doi:10.1016/j.celrep.2019.07.049.

Ryan, M.T., and N.J. Hoogenraad. 2007. Mitochondrial-nuclear communications. Annu Rev Biochem. 76:701–722. doi:10.1146/annurev.biochem.76.052305.091720.

Schaeffer, J.M., and M.R. Donatelli. 1990. Characterization of a high-affinity membrane-associated ornithine decarboxylase from the free-living nematode Caenorhabditis elegans. Biochem J. 270:599–604. doi:10.1042/bj2700599.

Schiavi, A., S. Maglioni, K. Palikaras, A. Shaik, F. Strappazzon, V. Brinkmann, A. Torgovnick, N. Castelein, S. De Henau, B.P. Braeckman, F. Cecconi, N. Tavernarakis, and N. Ventura. 2015. Iron-Starvation-Induced Mitophagy Mediates Lifespan Extension upon Mitochondrial Stress in C. elegans. Current Biology. 25:1810–1822. doi:10.1016/j.cub.2015.05.059.

Schober, F.A., D. Moore, I. Atanassov, M.F. Moedas, P. Clemente, Á. Végvári, N.E. Fissi, R. Filograna, A.-L. Bucher, Y. Hinze, M. The, E. Hedman, E. Chernogubova, A. Begzati, R. Wibom, M. Jain, R. Nilsson, L. Käll, A. Wedell, C. Freyer, and A. Wredenberg. 2021. The one-carbon pool controls mitochondrial energy metabolism via complex I and iron-sulfur clusters. Sci. Adv. 7:eabf0717. doi:10.1126/sciadv.abf0717.

Scoville, D.W., H.A. Cyphert, L. Liao, J. Xu, A. Reynolds, S. Guo, and R. Stein. 2015. MLL3 and MLL4 Methyltransferases Bind to the MAFA and MAFB Transcription Factors to Regulate Islet β-Cell Function. Diabetes. 64:3772–3783. doi:10.2337/db15-0281.

Shamalnasab, M., M. Dhaoui, M. Thondamal, E.B. Harvald, N.J. Færgeman, H. Aguilaniu, and P. Fabrizio. 2017. HIF-1–dependent regulation of lifespan in Caenorhabditis elegans by the acyl-CoA–binding protein MAA-1. Aging. 9:1745–1769. doi:10.18632/aging.101267.

Shao, L.-W., Q. Peng, M. Dong, K. Gao, Y. Li, Y. Li, C.-Y. Li, and Y. Liu. 2020. Histone deacetylase HDA-1 modulates mitochondrial stress response and longevity. Nat Commun. 11:4639. doi:10.1038/s41467-020-18501-w.

Sharma, M., R. Pandey, and D. Saluja. 2018. ROS is the major player in regulating altered autophagy and lifespan in sin-3 mutants of C. elegans. Autophagy. 14:1239–1255. doi:10.1080/15548627.2018.1474312.

Signes, A., and E. Fernandez-Vizarra. 2018. Assembly of mammalian oxidative phosphorylation complexes I–V and supercomplexes. Essays in Biochemistry. 62:255–270. doi:10.1042/EBC20170098.

Spanier, B., A. Laurençon, A. Weiser, N. Pujol, S. Omi, A. Barsch, A. Korf, S.W. Meyer, J.J. Ewbank, F. Palladino, S. Garvis, H. Aguilaniu, and M. Witting. 2021. Correction to: Comparison of lipidome profiles of Caenorhabditis elegans—results from an inter-laboratory ring trial. Metabolomics. 17:33. doi:10.1007/s11306-021-01784-5.

Taanman, J.-W. 1999. The mitochondrial genome: structure, transcription, translation and replication. Biochimica et Biophysica Acta (BBA) - Bioenergetics. 1410:103–123. doi:10.1016/S0005-2728(98)00161-3.

Tian, Y., G. Garcia, Q. Bian, K.K. Steffen, L. Joe, S. Wolff, B.J. Meyer, and A. Dillin. 2016. Mitochondrial Stress Induces Chromatin Reorganization to Promote Longevity and UPR mt. Cell. 165:1197–1208. doi:10.1016/j.cell.2016.04.011.

Til, H.P., H.E. Falke, M.K. Prinsen, and M.I. Willems. 1997. Acute and subacute toxicity of tyramine, spermidine, spermine, putrescine and cadaverine in rats. Food and Chemical Toxicology. 35:337–348. doi:10.1016/S0278-6915(97)00121-X.

To, T.-L., A.M. Cuadros, H. Shah, W.H.W. Hung, Y. Li, S.H. Kim, D.H.F. Rubin, R.H. Boe, S. Rath, J.K. Eaton, F. Piccioni, A. Goodale, Z. Kalani, J.G. Doench, D.E. Root, S.L. Schreiber, S.B. Vafai, and V.K. Mootha. 2019. A Compendium of Genetic Modifiers of Mitochondrial Dysfunction Reveals Intra-organelle Buffering. Cell. 179:1222–1238.e17. doi:10.1016/j.cell.2019.10.032.

Vincent, A.E., Y.S. Ng, K. White, T. Davey, C. Mannella, G. Falkous, C. Feeney, A.M. Schaefer, R. McFarland, G.S. Gorman, R.W. Taylor, D.M. Turnbull, and M. Picard. 2016. The Spectrum of Mitochondrial Ultrastructural Defects in Mitochondrial Myopathy. Sci Rep. 6:30610. doi:10.1038/srep30610.

Wai, T., and T. Langer. 2016. Mitochondrial Dynamics and Metabolic Regulation. Trends Endocrinol Metab. 27:105–117. doi:10.1016/j.tem.2015.12.001.

Wang, Y., Y. Zhang, L. Chen, Q. Liang, X.-M. Yin, L. Miao, B.-H. Kang, and D. Xue. 2016. Kinetics and specificity of paternal mitochondrial elimination in Caenorhabditis elegans. Nat Commun. 7:12569. doi:10.1038/ncomms12569.

Ward, D.M., and S.M. Cloonan. 2019. Mitochondrial Iron in Human Health and Disease. Annu Rev Physiol. 81:453–482. doi:10.1146/annurev-physiol-020518-114742.

Wei, W., and G. Ruvkun. 2020. Lysosomal activity regulates *Caenorhabditis elegans* mitochondrial dynamics through vitamin B12 metabolism. Proc. Natl. Acad. Sci. U.S.A. 117:19970–19981. doi:10.1073/pnas.2008021117.

Westermann, B. 2010. Mitochondrial fusion and fission in cell life and death. Nat Rev Mol Cell Biol. 11:872–884. doi:10.1038/nrm3013.

Whitnall, M., Y. Suryo Rahmanto, M.L.-H. Huang, F. Saletta, H.C. Lok, L. Gutiérrez, F.J. Lázaro, A.J. Fleming, T.G. St Pierre, M.R. Mikhael, P. Ponka, and D.R. Richardson. 2012. Identification of nonferritin mitochondrial iron deposits in a mouse model of Friedreich ataxia. Proc Natl Acad Sci U S A. 109:20590–20595. doi:10.1073/pnas.1215349109.

Witteveen, J.S., M.H. Willemsen, T.C.D. Dombroski, N.H.M. van Bakel, W.M. Nillesen, J.A. van Hulten, E.J.R. Jansen, D. Verkaik, H.E. Veenstra-Knol, C.M.A. van Ravenswaaij-Arts, J.S.K. Wassink-Ruiter, M. Vincent, A. David, C. Le Caignec, J. Schieving, C. Gilissen, N. Foulds, P. Rump, T. Strom, K. Cremer, A.M. Zink, H. Engels, S.A. de Munnik, J.E. Visser, H.G. Brunner, G.J.M. Martens, R. Pfundt, T. Kleefstra, and S.M. Kolk. 2016. Haploinsufficiency of MeCP2-interacting transcriptional co-repressor SIN3A causes mild intellectual disability by affecting the development of cortical integrity. Nat Genet. 48:877–887. doi:10.1038/ng.3619.

Yang, R., Y. Li, Y. Wang, J. Zhang, Q. Fan, J. Tan, W. Li, X. Zou, and B. Liang. 2022. NHR-80 senses the mitochondrial UPR to rewire citrate metabolism for lipid accumulation in Caenorhabditis elegans. Cell Reports. 38:110206. doi:10.1016/j.celrep.2021.110206.

Yang, X., S.M. Graff, C.N. Heiser, K.-H. Ho, B. Chen, A.J. Simmons, A.N. Southard-Smith, G. David, D.A. Jacobson, I. Kaverina, C.V.E. Wright, K.S. Lau, and G. Gu. 2020a. Coregulator Sin3a Promotes Postnatal Murine β-Cell Fitness by Regulating Genes in Ca2+ Homeostasis, Cell Survival, Vesicle Biosynthesis, Glucose Metabolism, and Stress Response. Diabetes. 69:1219– 1231. doi:10.2337/db19-0721.

Yang, X., M. Zhang, Y. Dai, Y. Sun, Y. Aman, Y. Xu, P. Yu, Y. Zheng, J. Yang, and X. Zhu. 2020b. Spermidine inhibits neurodegeneration and delays aging via the PINK1-PDR1-dependent mitophagy pathway in C. elegans. Aging. 12:16852–16866. doi:10.18632/aging.103578.

Ye, C., B.M. Sutter, Y. Wang, Z. Kuang, and B.P. Tu. 2017. A Metabolic Function for Phospholipid and Histone Methylation. Molecular Cell. 66:180–193.e8. doi:10.1016/j.molcel.2017.02.026.

Yoneda, T., C. Benedetti, F. Urano, S.G. Clark, H.P. Harding, and D. Ron. 2004. Compartment-specific perturbation of protein handling activates genes encoding mitochondrial chaperones. Journal of Cell Science. 117:4055–4066. doi:10.1242/jcs.01275.

Zhou, Q., H. Li, H. Li, A. Nakagawa, J.L.J. Lin, E.-S. Lee, B.L. Harry, R.R. Skeen-Gaar, Y. Suehiro, D. William, S. Mitani, H.S. Yuan, B.-H. Kang, and D. Xue. 2016. Mitochondrial endonuclease G mediates breakdown of paternal mitochondria upon fertilization. Science. 353:394–399. doi:10.1126/science.aaf4777.

Zhu, D., X. Wu, J. Zhou, X. Li, X. Huang, J. Li, J. Wu, Q. Bian, Y. Wang, and Y. Tian. 2020. NuRD mediates mitochondrial stress-induced longevity via chromatin remodeling in response to acetyl-CoA level. Sci Adv. 6:eabb2529. doi:10.1126/sciadv.abb2529.

Zhu, F., Q. Zhu, D. Ye, Q. Zhang, Y. Yang, X. Guo, Z. Liu, Z. Jiapaer, X. Wan, G. Wang, W. Chen, S. Zhu, C. Jiang, W. Shi, and J. Kang. 2018. Sin3a–Tet1 interaction activates gene transcription and is required for embryonic stem cell pluripotency. Nucleic Acids Research. 46:6026–6040. doi:10.1093/nar/gky347.

