## Supplementary figures and images for "SIN-3 transcriptional coregulator maintains mitochondrial homeostasis and polyamine flux"

### Supplemental Figure 1

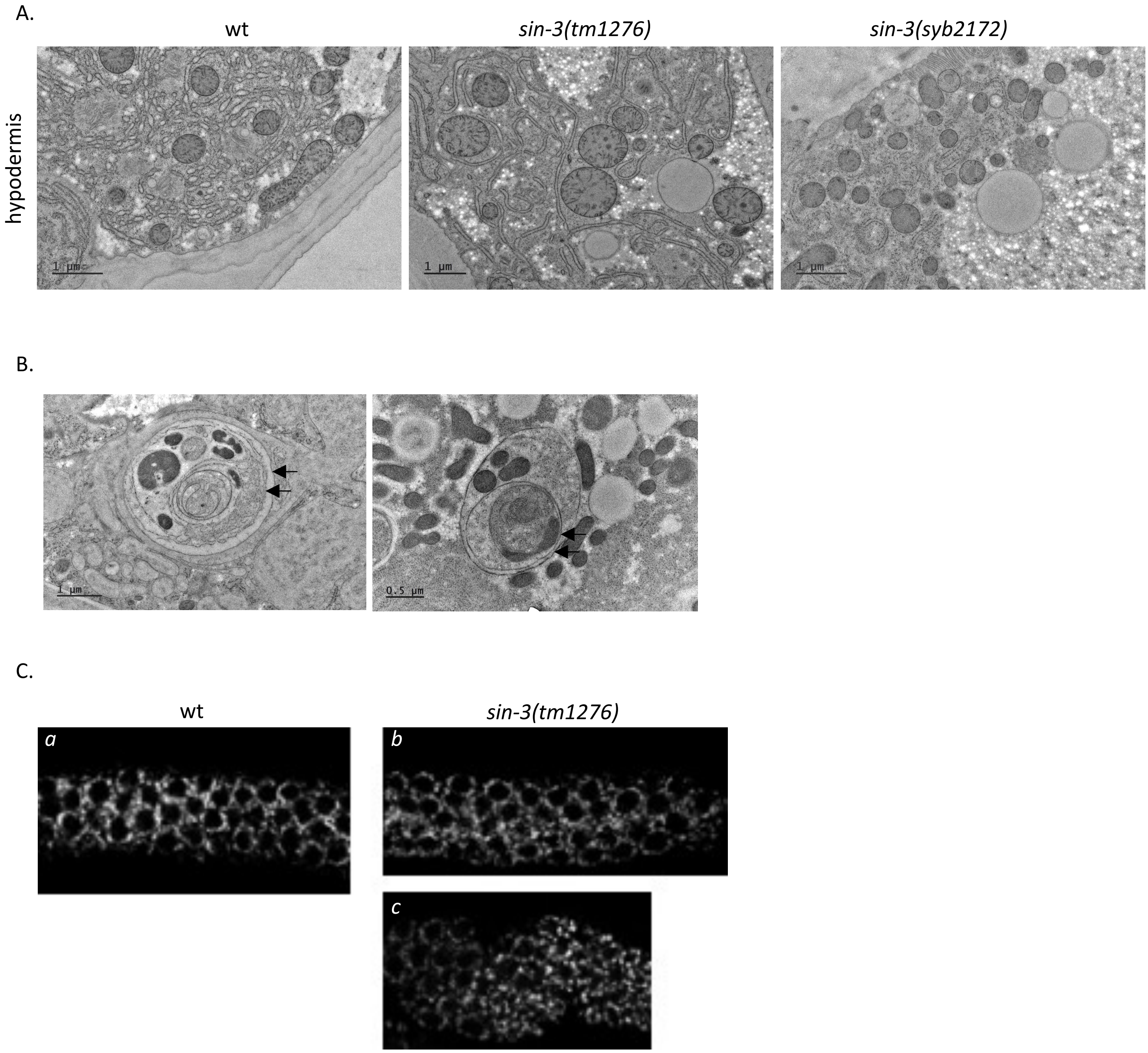

### Supplemental Figure 2

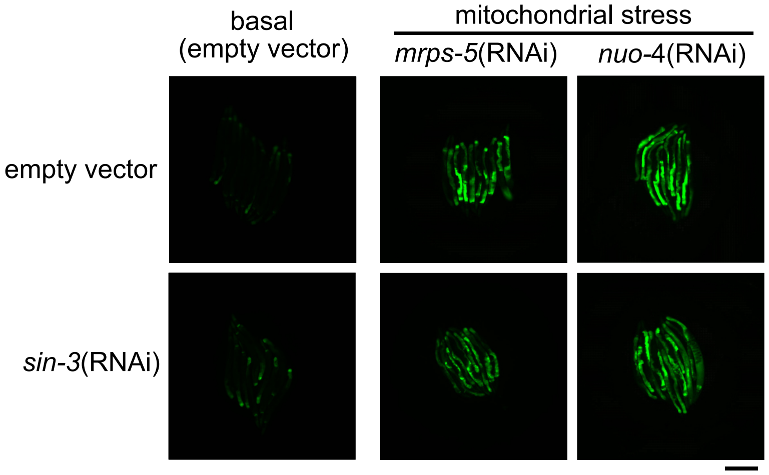

### Supplemental Figure 3

Figure S3

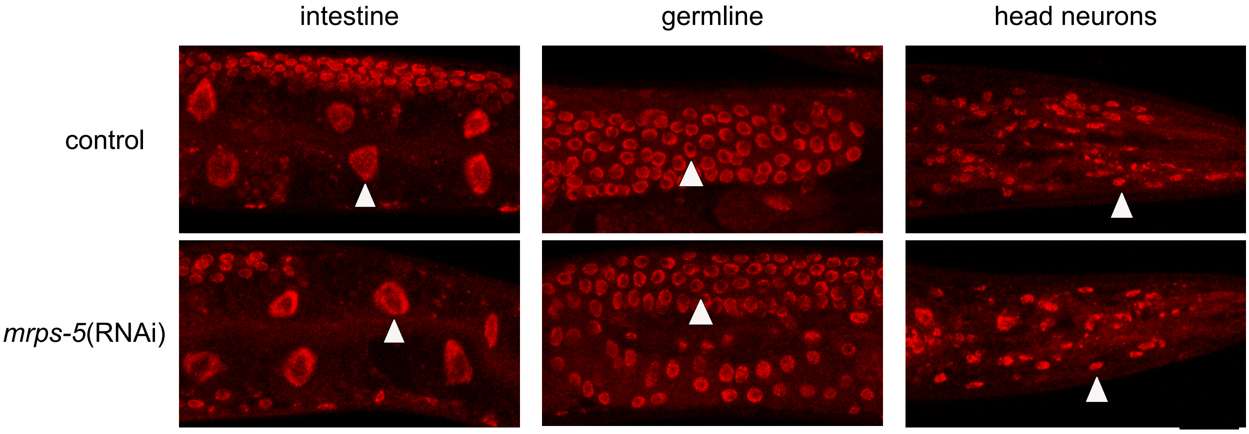

### Supplemental Figure 4A

Figure S4A; CoAs and carnitines

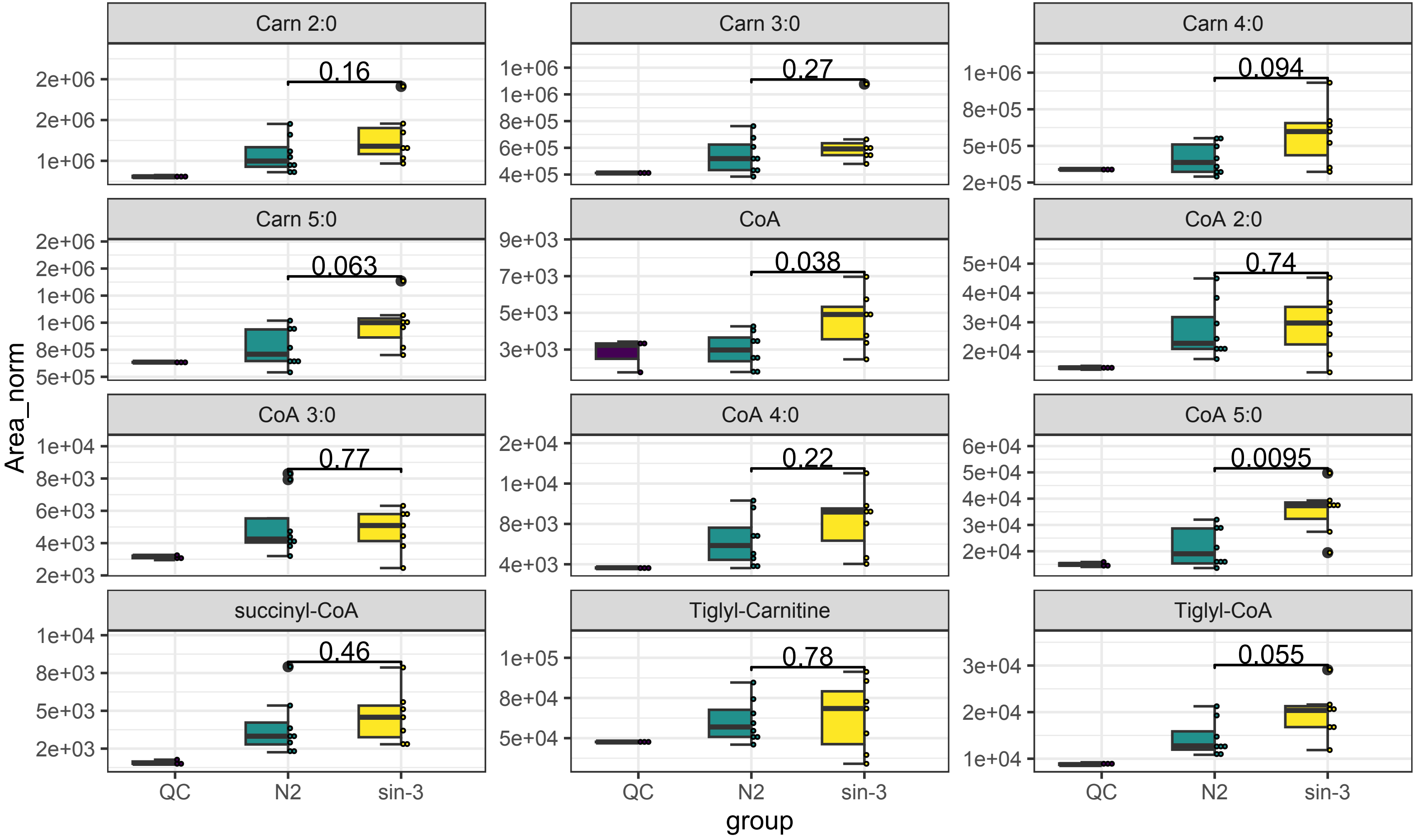

### Supplemental Figure 4B

Figure S4B; TCA and others

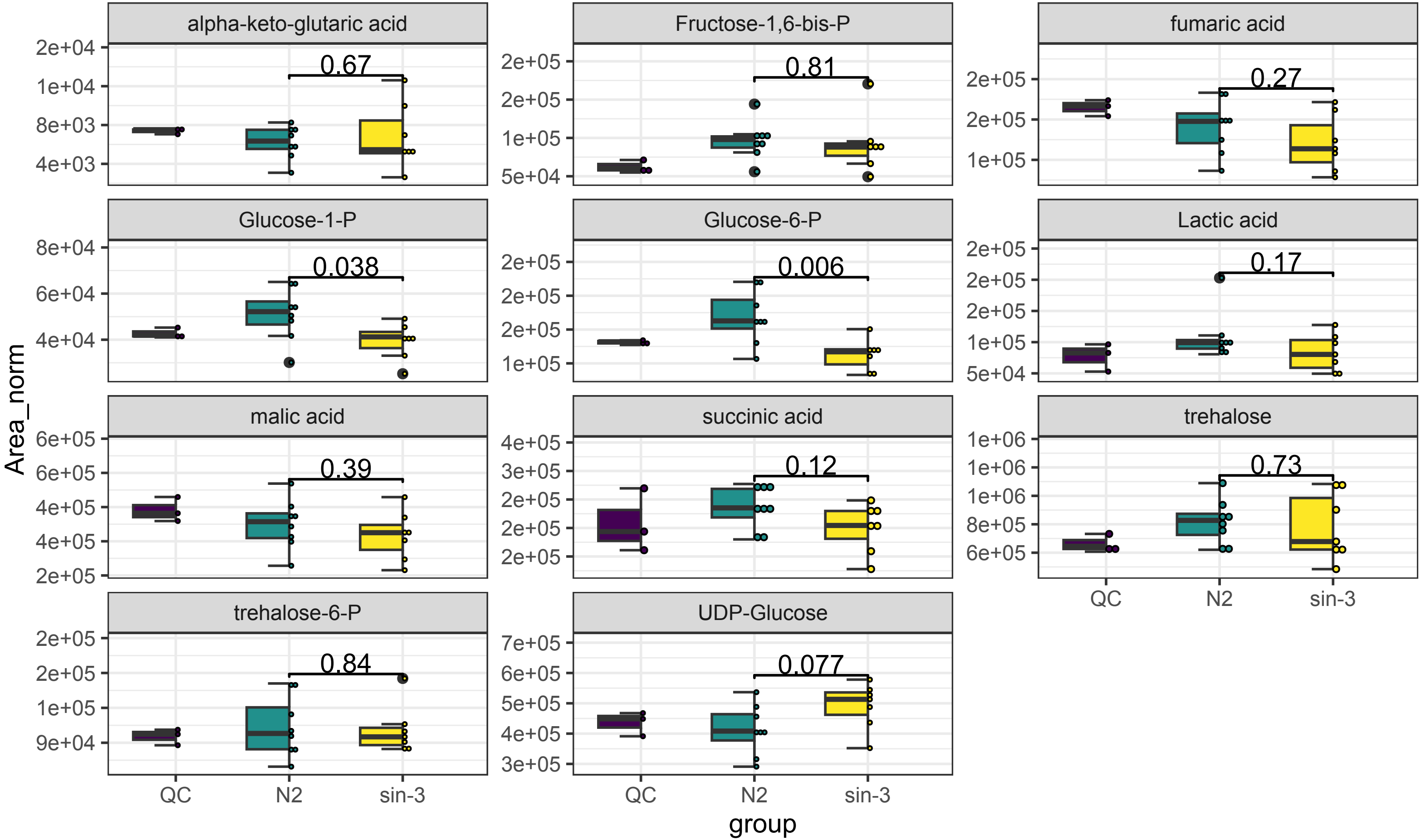

### Supplemental Figure 4C

Figure S4C; amino acid and related

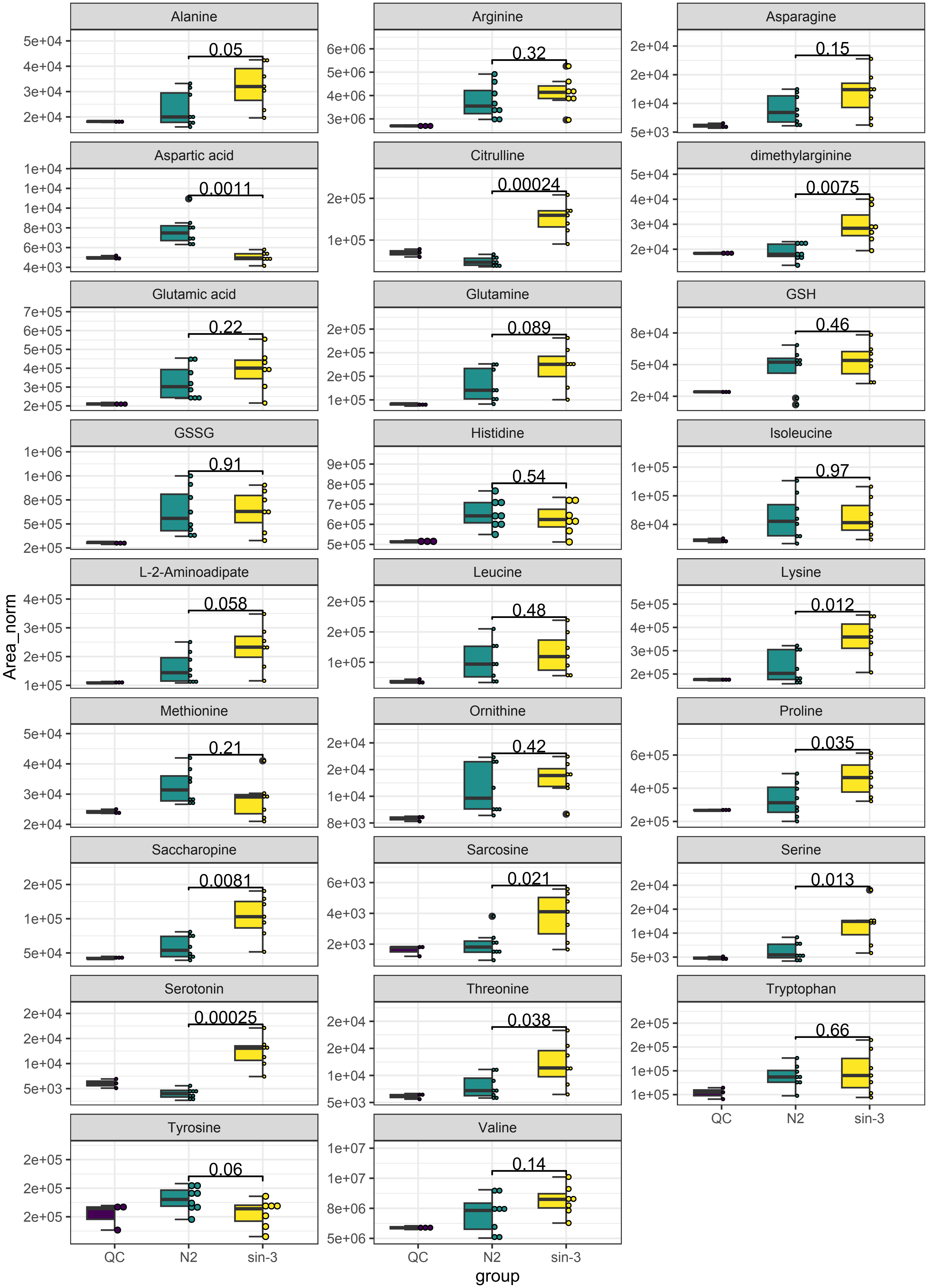

### Supplemental Figure 4D

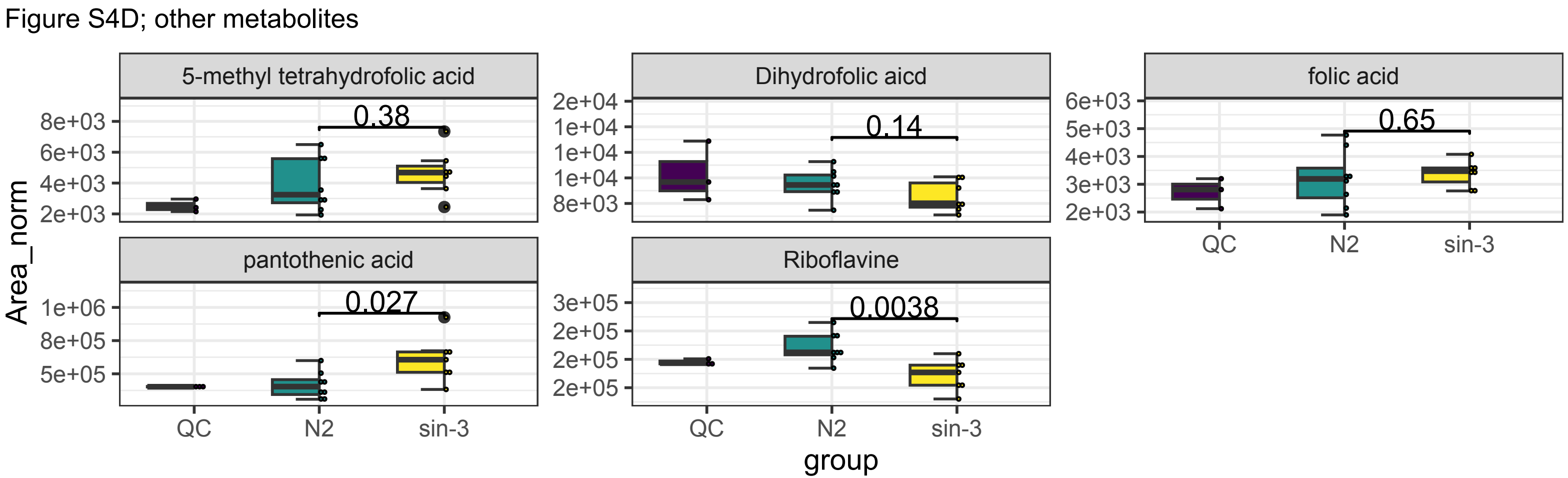

### Supplemental Figure 4E

Figure S4E; SAM and polyamines

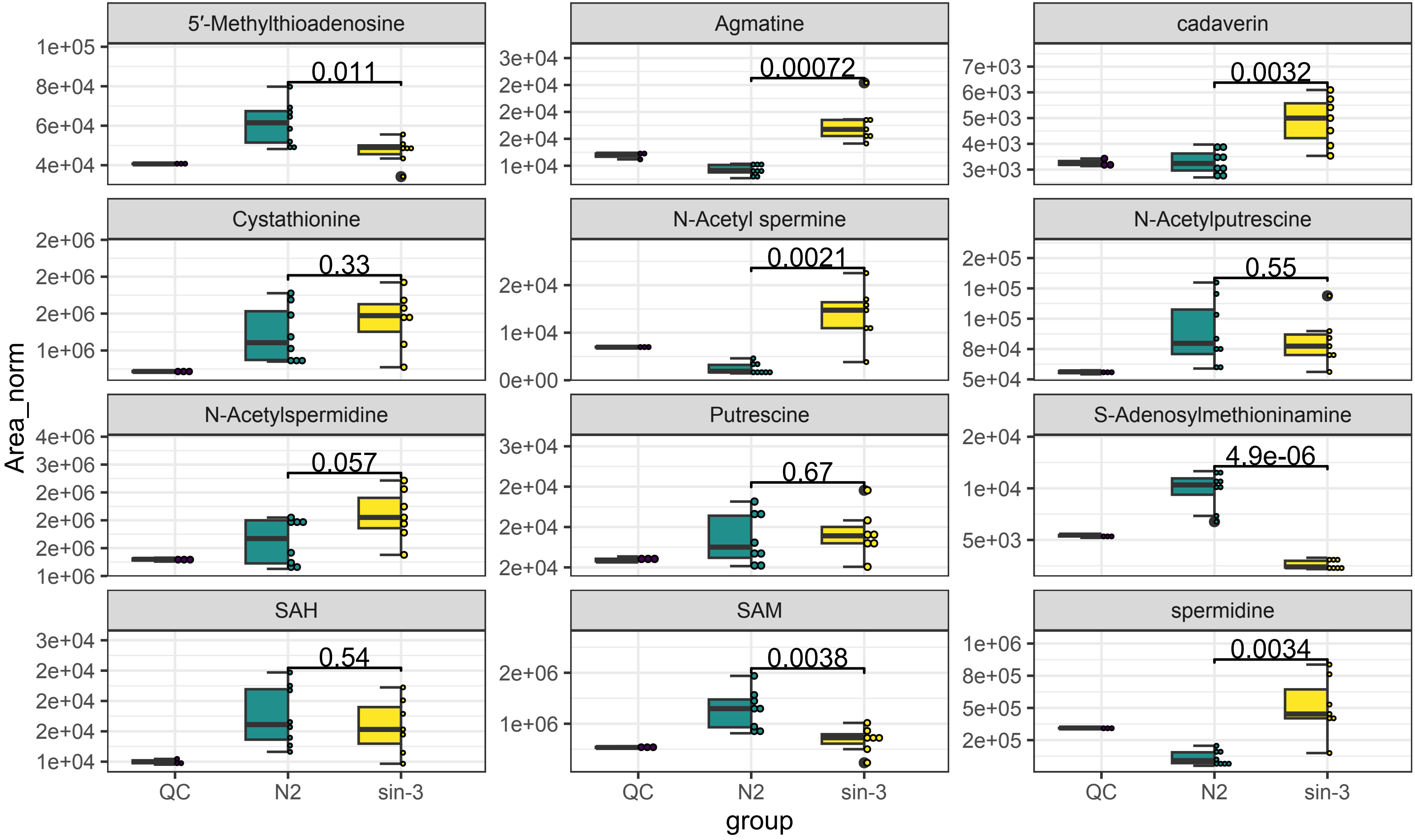
