## Supplemental Table1 for "SIN-3 transcriptional coregulator maintains mitochondrial homeostasis and polyamine flux"

**Table S1 : List of significant commonly misregulated genes in *sin-3(tm1276)* and *sin-3(syb21 72)* compared to wild-type**

| id_gene | gene_name | Log2.FC_tm1 | padj_tm1276 | Log2.FC_syb | padj_syb2172 |
|---|---|---|---|---|---|
| WBGene00000 | act-2 | -0.7337908935 | 4.6777629160 | -1.3402953390 | 0.00388591688508462 |
| WBGene00000 | agt-1 | -0.5074901861 | 2.2471986508 | -1.3241776459 | 0.000298497174038359 |
| WBGene00000 | agt-2 | -0.5693187693 | 3.2229899386 | -0.8925756648 | 0.0363039359761594 |
| WBGene00000 | alh-8 | -0.8871072606 | 3.2512867891 | -1.0480043824 | 0.000941745667591152 |
| WBGene00000 | apc-10 | -1.3101957701 | 4.3810443309 | -2.0494584607 | 2.78971557688454e-07 |
| WBGene00000 | apc-11 | -0.3589899004 | 1.7461620344 | -1.1518369501 | 0.00378024821546395 |
| WBGene00000 | arf-3 | -0.2073248196 | 4.2385913681 | -0.7832594046 | 0.0396665340979832 |
| WBGene00000 | arx-7 | -0.4237311310 | 1.4169793737 | -1.4853415957 | 0.000629068370943816 |
| WBGene00000 | asp-1 | -1.1439227516 | 0.0070392156 | 1.9633212522 | 0.0402998074998023 |
| WBGene00000 | asp-6 | -1.3206931081 | 0.0065410198 | 2.9998061931 | 0.0119879695899209 |
| WBGene00000 | bli-4 | -0.8585983354 | 1.9027740050 | -1.2224593886 | 0.000129397405920671 |
| WBGene00000 | bpl-1 | -1.3999098793 | 5.5058664536 | -2.3117290683 | 1.97362331745253e-10 |
| WBGene00000 | ccf-1 | -0.3599930022 | 1.5459843331 | -0.6978171117 | 0.0442907004844807 |
| WBGene00000 | cdd-2 | -0.8113385697 | 6.3865066050 | -1.6386297869 | 3.31185720168559e-05 |
| WBGene00000 | cdk-1 | -0.1592949382 | 0.0002992521 | -0.7680852667 | 0.0337165143815552 |
| WBGene00000 | cdk-5 | -0.5833238832 | 1.3801606288 | -1.0913610946 | 0.000903237471222072 |
| WBGene00000 | cey-2 | -0.6733215617 | 1.5700492576 | -1.9271667090 | 1.0420981813283e-06 |
| WBGene00000 | cey-3 | -0.3387835175 | 4.9310470973 | -1.3713276095 | 0.000179216214333118 |
| WBGene00000 | cgh-1 | -0.3717875748 | 0.0057237951 | -1.2954968683 | 8.00021391507808e-05 |
| WBGene00000 | chp-1 | -0.1511256723 | 0.0116949242 | -0.9872204511 | 0.00754173239372364 |
| WBGene00000 | clh-6 | -0.2427904167 | 1.8712661756 | -0.8702924870 | 0.0465138372032017 |
| WBGene00000 | coq-3 | -0.2246779888 | 0.0015724725 | -1.0493901119 | 0.0233759444726421 |
| WBGene00000 | cpb-1 | -1.8109856009 | 5.0314156873 | -1.7551213789 | 0.000292439353591756 |
| WBGene00000 | cpn-1 | -0.9708363551 | 7.8863110980 | -1.3373542433 | 0.00274369066672906 |
| WBGene00000 | crn-2 | -0.4353830356 | 8.6080892246 | -0.9538047627 | 0.0434236144179205 |
| WBGene00000 | cul-1 | -0.9128845631 | 6.9424928069 | -0.7292892218 | 0.043906502119357 |
| WBGene00000 | cul-5 | -0.8863130371 | 3.2195335421 | -1.1204560253 | 0.0033531951019102 |
| WBGene00000 | cyn-5 | -0.3182618676 | 1.9909270866 | -1.0295372387 | 0.00723497380251161 |
| WBGene00000 | cyn-7 | -0.4210380440 | 7.4749138189 | -1.1067966082 | 0.02375596895563 |
| WBGene00000 | cyn-10 | -0.7261455461 | 1.1899076571 | -1.3073272412 | 0.00219544124564106 |
| WBGene00000 | cyn-12 | -0.6042960662 | 1.2483490299 | -1.0876181795 | 0.0076207331370829 |
| WBGene00000 | hsp-90 | -0.6388899726 | 3.6320870766 | -1.1007139090 | 0.0173374876563467 |
| WBGene00000 | daf-25 | -0.4194945366 | 6.8361972336 | -0.8186805715 | 0.0465138372032017 |
| WBGene00000 | dbr-1 | -0.8294941257 | 3.7644594303 | -1.0684207604 | 0.00242437861275097 |
| WBGene00000 | dhs-11 | -0.6805876527 | 2.8716528845 | -1.3896285706 | 0.000268972261359105 |

| | | | | | |
|---|---|---|---|---|---|
| WBGene00000 | dhs-12 | -1.4379477228 | 2.7960223819 | -2.1503922821 | 9.70332770424679e-07 |
| WBGene00001 | dnc-2 | -0.6674191931 | 2.2272846123 | -1.1069358404 | 0.00262843922352046 |
| WBGene00001 | dnj-17 | -1.0315362601 | 8.3385682268 | -0.8155077036 | 0.0474925382769167 |
| WBGene00001 | dnj-19 | -0.5613004052 | 2.0171113408 | -0.7680322304 | 0.0430134412966845 |
| WBGene00001 | eef-2 | -0.4388652776 | 2.1634932679 | -1.3580958772 | 0.000789757293431072 |
| WBGene00001 | eef-1A.1 | -0.6240433084 | 3.4319756153 | -1.4473476432 | 0.00356305659172655 |
| WBGene00001 | eif-3.G | -0.3782749036 | 3.9686728579 | -0.9846433199 | 0.0107665712442118 |
| WBGene00001 | eif-3.K | -0.3485575364 | 2.4520370689 | -1.0353506253 | 0.00355483757279993 |
| WBGene00001 | elb-1 | -0.4605590273 | 1.4231022422 | -0.8868568727 | 0.029792358284338 |
| WBGene00001 | emo-1 | -0.6941068448 | 5.8126929036 | -1.0475027280 | 0.00536349951241789 |
| WBGene00001 | qars-1 | -0.5048482349 | 3.9842759685 | -1.0314160255 | 0.00316729511205931 |
| WBGene00001 | exl-1 | -1.2593020110 | 4.2664734374 | -1.4494594178 | 0.00089446647563869 |
| WBGene00001 | fbp-1 | -0.4604296898 | 2.0758425313 | -1.2440536630 | 0.00162191545186003 |
| WBGene00001 | fem-2 | -0.5543916935 | 1.5779667829 | -0.9963951934 | 0.00330240458741758 |
| WBGene00001 | fkb-1 | -0.9165824389 | 2.8595502654 | -1.0931924668 | 0.0321753128310218 |
| WBGene00001 | frm-10 | -1.0454608694 | 8.7209391281 | -1.3366946651 | 0.00262843922352046 |
| WBGene00001 | fum-1 | -1.5729436556 | 5.6763212785 | -1.7551863864 | 1.0420981813283e-06 |
| WBGene00001 | gln-6 | -0.6173531983 | 9.6868572065 | -1.1989267488 | 0.00112019944442587 |
| WBGene00001 | gna-2 | -0.5041856027 | 2.0886056703 | -0.8275668483 | 0.0116603908719766 |
| WBGene00001 | gpd-4 | -0.9806284855 | 1.0894149170 | -1.1805851953 | 0.0218364924533932 |
| WBGene00001 | gst-1 | -0.3377534117 | 2.8227854668 | -1.2472131894 | 0.000390481151327125 |
| WBGene00001 | hil-4 | -0.1250670382 | 0.0149370295 | -1.1345489792 | 0.00809816817866127 |
| WBGene00001 | hil-5 | -0.5148861366 | 1.3117087075 | -1.5101345023 | 5.32190669710519e-05 |
| WBGene00001 | him-5 | -0.9643073232 | 1.0539857312 | -1.3069079294 | 0.0145394758785685 |
| WBGene00001 | him-6 | -0.7432588933 | 2.3051074668 | -0.9426179148 | 0.00622535664943638 |
| WBGene00001 | him-14 | -1.0609457528 | 5.6588696803 | -1.2521427405 | 0.00350810133964749 |
| WBGene00001 | his-35 | -0.9651596104 | 5.4422261131 | -1.7065463547 | 7.16906857710512e-05 |
| WBGene00001 | C50F4.6 | -0.8067511484 | 1.5980197957 | -1.5907415455 | 1.04848973078758e-06 |
| WBGene00001 | his-37 | -0.8155011572 | 1.2050953930 | -1.7201528837 | 1.7408386620663e-06 |
| WBGene00001 | his-48 | -0.5571205497 | 6.1163611875 | -1.7386376088 | 0.0178821293999925 |
| WBGene00001 | his-59 | -0.8935714087 | 3.1405607809 | -2.0966537003 | 1.53330988297995e-06 |
| WBGene00001 | his-60 | -0.8866623142 | 5.7805917329 | -2.7282152167 | 1.41501011006282e-13 |
| WBGene00001 | his-61 | -0.9038038228 | 5.7248314144 | -2.4433133862 | 9.03744198071171e-05 |
| WBGene00001 | his-62 | -0.9172401706 | 1.6070908179 | -2.9157638696 | 1.65204800580944e-11 |
| WBGene00001 | his-63 | -0.2180722657 | 0.0147780938 | -1.9787675305 | 0.00231355529884879 |
| WBGene00002 | icd-1 | -0.2288583599 | 5.6235155193 | -1.0107304741 | 0.0433529837227188 |
| WBGene00002 | ife-3 | -0.4076657628 | 1.2880294737 | -1.0748219721 | 0.00162191545186003 |
| WBGene00002 | iff-1 | -0.3462929443 | 5.7613774166 | -1.7258843845 | 9.88795158767315e-05 |

| | | | | | |
|---|---|---|---|---|---|
| WBGene00002 | ile-1 | -0.865440081€ | 1.810972325‡ | -1.020704288ℇ | 0.00136329489293447 |
| WBGene00002 | inx-14 | -0.8988255879 | 1.0407711066 | -2.1124269707 | 1.61990188885901e-12 |
| WBGene00002 | klp-10 | -0.1464900257 | 0.0147885897 | -1.2056839658 | 0.000622639997514217 |
| WBGene00002 | klp-16 | -0.5885222183 | 5.4558422659 | -1.1566351277 | 0.00437687905729372 |
| WBGene00002 | kars-1 | -0.3167119557 | 7.8485025541 | -0.914220563C | 0.0144548481156194 |
| WBGene00002 | lap-2 | -1.4053324887 | 1.0750228588 | -1.8229049964 | 2.17325367568778e-09 |
| WBGene00002 | lat-1 | -0.1967085498 | 1.8093003056 | 0.9396027736 | 0.0112437461973556 |
| WBGene00002 | lig-1 | -0.6079772488 | 1.3537598196 | -0.8235740368 | 0.0116603908719766 |
| WBGene00003 | lin-36 | -0.6254649049 | 4.8230406702 | -1.2967176993 | 0.0336672285262766 |
| WBGene00003 | lip-1 | -0.711582741C | 1.6292208652 | -1.2341964768 | 0.00558748894629725 |
| WBGene00003 | lsm-1 | -0.3417173615 | 3.9245755791 | -1.0598765006 | 0.008865975072662 |
| WBGene00003 | mec-15 | -0.2295755098 | 4.0297040220 | -0.821595577C | 0.0148951764508183 |
| WBGene00003 | mxl-1 | -0.5388407371 | 7.8275553041 | -0.971179888C | 0.00597326454241204 |
| WBGene00003 | mxl-2 | -0.5393565462 | 3.4676362753 | -1.226897990C | 0.000749009360469706 |
| WBGene00003 | ndx-6 | -1.0319850623 | 1.2816030900 | -1.569495439C | 0.00491233666167212 |
| WBGene00003 | nhr-61 | -0.992514501C | 8.4935118857 | -1.198159527C | 0.0173238639391292 |
| WBGene00003 | npp-16 | -0.4455928347 | 1.8531743469 | -0.8848234443 | 0.0402998074998023 |
| WBGene00003 | ntl-2 | -0.6882065484 | 1.5334262074 | -0.873586107C | 0.0243900038821214 |
| WBGene00003 | pab-1 | -0.3339409934 | 2.7068749034 | -1.3551435446 | 0.00187001525247731 |
| WBGene00003 | pas-1 | -0.2623983344 | 1.0232978342 | -0.9205474162 | 0.00677370412473355 |
| WBGene00003 | pas-2 | -0.6069178869 | 1.3804406865 | -1.0171138248 | 0.0136490014199656 |
| WBGene00003 | pbs-3 | -0.5899410603 | 1.3150710082 | -0.8344276693 | 0.0223806327271981 |
| WBGene00003 | pbs-7 | -0.3535193198 | 6.6217295391 | -0.8821034924 | 0.0156574305595428 |
| WBGene00003 | pdr-1 | -0.5576817916 | 4.0575677010 | -0.7335878719 | 0.0463171456168638 |
| WBGene00004 | plp-1 | -0.7432968811 | 3.8677538039 | -1.160029941C | 0.000267187769893407 |
| WBGene00004 | pms-2 | -0.7669307935 | 4.9123826835 | -0.9934945553 | 0.0207387346240917 |
| WBGene00004 | stam-1 | -1.1463001873 | 3.6613381358 | -0.9420714553 | 0.00725539903713165 |
| WBGene00004 | sin-3 | -1.0532992737 | 3.1038429569 | -10.630972264 | 4.8522884794687e-31 |
| WBGene00004 | pqn-51 | -0.6961635962 | 1.1206666066 | -0.8657252768 | 0.0164796048661349 |
| WBGene00004 | psr-1 | -0.441343271C | 1.1936327305 | -0.8892430936 | 0.00908265097152629 |
| WBGene00004 | puf-5 | -0.6335012321 | 2.4772822100 | -1.0418004738 | 0.00595323709087748 |
| WBGene00004 | puf-10 | -0.4553258804 | 4.9857248266 | -0.8905655287 | 0.0454929174258304 |
| WBGene00004 | rab-6.1 | -0.8632007402 | 2.5581596407 | -1.2989039273 | 0.000126432142977859 |
| WBGene00004 | ran-1 | -0.0970982897 | 0.0085010407 | -1.0555008946 | 0.00985659341951121 |
| WBGene00004 | rbx-1 | -0.5069107471 | 1.1424488300 | -0.8548074721 | 0.0494474158666274 |
| WBGene00004 | rde-4 | -0.2540430478 | 1.7684604548 | -0.7615112145 | 0.0321753128310218 |
| WBGene00004 | rec-1 | -0.754288172C | 4.5165455960 | -1.156391712C | 0.0214213514035014 |
| WBGene00004 | rfc-3 | -0.3573178074 | 1.6872252018 | -0.965357768C | 0.00204518906207917 |

| | | | | | |
|---|---|---|---|---|---|
| WBGene00004 | rfs-1 | -0.8534184100 | 3.9145513497 | -2.3379747838 | 0.0169530127113724 |
| WBGene00004 | rme-2 | -0.4625018934 | 1.5062840506 | -1.3077178694 | 1.85524941163152e-05 |
| WBGene00004 | rla-0 | -0.4083455261 | 8.0077306538 | -1.6401879249 | 0.00108793603484848 |
| WBGene00004 | rla-1 | -0.3932678808 | 2.8992289121 | -2.1394382648 | 1.94680928232503e-05 |
| WBGene00004 | rla-2 | -0.2330882339 | 5.2745005176 | -1.1568857916 | 0.00953480849473906 |
| WBGene00004 | rpl-1 | -0.3804133921 | 1.9729830284 | -1.6932607257 | 0.000689030759008703 |
| WBGene00004 | rpl-2 | -0.3700555942 | 1.2117831212 | -1.2202383874 | 0.0156468961960255 |
| WBGene00004 | rpl-3 | -0.3294480168 | 3.7276067183 | -1.4556824908 | 0.00470154847198345 |
| WBGene00004 | rpl-4 | -0.3622434460 | 7.8281827094 | -1.3902556610 | 0.00369637835521812 |
| WBGene00004 | rpl-6 | -0.2708104065 | 4.6538605452 | -1.4577076530 | 0.00595323709087748 |
| WBGene00004 | rpl-7 | -0.2481825612 | 4.9218361615 | -1.2705487617 | 0.0150329335626997 |
| WBGene00004 | rpl-7A | -0.3247485131 | 2.4313217166 | -1.6514520042 | 0.000651782091240334 |
| WBGene00004 | rpl-9 | -0.6739577912 | 3.5572850255 | -1.8588451135 | 5.23344157137434e-05 |
| WBGene00004 | rpl-10 | -0.4369016013 | 2.5390871050 | -1.4817045911 | 0.00294785822294085 |
| WBGene00004 | rpl-11.1 | -0.3769524823 | 7.3153874686 | -1.5071279023 | 0.00256401156083131 |
| WBGene00004 | rpl-12 | -0.2938576649 | 3.7452473223 | -1.5937970132 | 0.00116343402047996 |
| WBGene00004 | rpl-13 | -0.3016035061 | 2.4729009056 | -1.4344772951 | 0.00645439812875181 |
| WBGene00004 | rpl-14 | -0.4715140939 | 4.2341010851 | -1.5699932357 | 0.000958347340252515 |
| WBGene00004 | rpl-15 | -0.3774457435 | 8.5957987735 | -1.4510327325 | 0.00768576189551562 |
| WBGene00004 | rpl-16 | -0.3318900813 | 7.0063422339 | -1.4690798572 | 0.00539452957835146 |
| WBGene00004 | rpl-17 | -0.3782002740 | 3.0699281401 | -1.4843369007 | 0.00534762234784027 |
| WBGene00004 | rpl-18 | -0.4521054339 | 4.8220196990 | -1.4236226782 | 0.00388591688508462 |
| WBGene00004 | rpl-19 | -0.4021779037 | 6.1949177265 | -1.5351761901 | 0.0024790258954136 |
| WBGene00004 | rpl-20 | -0.3215015728 | 3.2190374072 | -1.5435452679 | 0.00215491874653192 |
| WBGene00004 | rpl-21 | -0.4687972004 | 4.7387629116 | -1.7766481118 | 0.000336985098138496 |
| WBGene00004 | rpl-22 | -0.3808321265 | 2.4817002633 | -1.6960506568 | 0.000133307543909504 |
| WBGene00004 | rpl-23 | -0.1706133376 | 0.0061330386 | -1.4816191824 | 0.00353409625586477 |
| WBGene00004 | rpl-24.1 | -0.3254967732 | 1.8557772896 | -1.5754443836 | 0.00263876395481626 |
| WBGene00004 | rpl-25.2 | -0.5462434042 | 6.6495366415 | -1.9330320763 | 1.20231853332228e-05 |
| WBGene00004 | rpl-26 | -0.2992506342 | 8.4892576284 | -1.4257370654 | 0.0039835275127462 |
| WBGene00004 | rpl-27 | -0.2583045780 | 7.1635550474 | -1.7090125081 | 0.0002890451114692 |
| WBGene00004 | rpl-28 | -0.4788453989 | 2.5745767373 | -1.8621309397 | 3.3054287899841e-05 |
| WBGene00004 | rpl-29 | -0.4322833769 | 2.2930263149 | -1.5425365234 | 0.00185435322591254 |
| WBGene00004 | rpl-30 | -0.3710573047 | 7.1724306738 | -1.4897670125 | 0.00128421156701525 |
| WBGene00004 | rpl-31 | -0.3321938383 | 1.3114516515 | -1.7126013896 | 0.000183374429730536 |
| WBGene00004 | rpl-32 | -0.5333899366 | 4.3810852644 | -1.8468371200 | 0.000118714794143051 |
| WBGene00004 | rpl-33 | -0.3438038436 | 2.9040222508 | -1.4530021316 | 0.00471864363186537 |
| WBGene00004 | rpl-34 | -0.3883493205 | 7.1176623062 | -1.5882137546 | 0.00103691264406009 |

| | | | | | |
|---|---|---|---|---|---|
| WBGene00004 | rpl-35 | -0.4074353098 | 6.5748880996 | -1.6417385048 | 0.000609852746924961 |
| WBGene00004 | rpl-36 | -0.4456718816 | 7.0820972295 | -1.7771999833 | 0.000254238044499091 |
| WBGene00004 | rpl-38 | -0.2928807931 | 1.3737374650 | -1.5577345510 | 0.00193076697382952 |
| WBGene00004 | rpl-39 | -0.5540409542 | 8.5285770963 | -1.5044532819 | 0.00294492304896226 |
| WBGene00004 | rpl-36.A | -0.3725950875 | 2.9015780671 | -1.5846622280 | 0.000762435275597351 |
| WBGene00004 | rpl-43 | -0.4515092511 | 3.0851189726 | -1.6327215266 | 0.000689030759008703 |
| WBGene00004 | rpn-7 | -0.3910406748 | 8.1014571188 | -0.8482889124 | 0.043156163420299 |
| WBGene00004 | rps-0 | -0.3983893109 | 2.9027289726 | -1.7089598964 | 0.000499516337434342 |
| WBGene00004 | rps-1 | -0.3864440134 | 3.5652088315 | -1.3415200368 | 0.00664951028719712 |
| WBGene00004 | rps-2 | -0.4400148164 | 1.1750864907 | -1.4921060663 | 0.00190697006507797 |
| WBGene00004 | rps-3 | -0.3935082217 | 8.6068525655 | -1.5516041935 | 0.00124182782482806 |
| WBGene00004 | rps-4 | -0.4074181351 | 1.7314623305 | -1.5424429826 | 0.00102379428314494 |
| WBGene00004 | rps-5 | -0.4539836897 | 4.6651990542 | -1.6177178026 | 0.00162191545186003 |
| WBGene00004 | rps-6 | -0.3816209657 | 1.9437111799 | -1.4133075781 | 0.00351398392465421 |
| WBGene00004 | rps-7 | -0.3700926017 | 2.8386142415 | -1.5993393193 | 0.00113733838842422 |
| WBGene00004 | rps-8 | -0.6469713667 | 8.6370801570 | -1.9487690363 | 8.48708236411893e-05 |
| WBGene00004 | rps-9 | -0.4484421675 | 1.4216531526 | -1.7762333460 | 0.000234933118014274 |
| WBGene00004 | rps-10 | -0.4155159035 | 5.9055542044 | -1.6358715202 | 0.000867385463938174 |
| WBGene00004 | rps-11 | -0.5508690208 | 5.5401754301 | -1.7993471689 | 0.000116830216264568 |
| WBGene00004 | rps-12 | -0.5309881268 | 3.0980813049 | -1.7633760575 | 5.80777911736705e-05 |
| WBGene00004 | rps-13 | -0.3696235906 | 1.5844551122 | -1.5421879680 | 0.0013203911812113 |
| WBGene00004 | rps-14 | -0.5550313878 | 7.4114390760 | -1.9682518130 | 3.3054287899841e-05 |
| WBGene00004 | rps-15 | -0.4209598649 | 8.6872404037 | -1.6072294472 | 0.000714919184242183 |
| WBGene00004 | rps-16 | -0.5245532836 | 7.0263697042 | -2.0533369478 | 8.44127148142459e-06 |
| WBGene00004 | rps-17 | -0.3909382258 | 4.0914698194 | -1.4830265442 | 0.00283066552464399 |
| WBGene00004 | rps-18 | -0.3274262870 | 4.7402347794 | -1.5580871678 | 0.00189260944960837 |
| WBGene00004 | rps-19 | -0.4574747485 | 8.6105071432 | -1.7090277256 | 0.000163958780800124 |
| WBGene00004 | rps-20 | -0.3425099074 | 3.9755370059 | -1.3695529463 | 0.00230390817754383 |
| WBGene00004 | rps-21 | -0.4514337455 | 9.8552331082 | -1.6694605304 | 0.000369930480878299 |
| WBGene00004 | rps-22 | -0.3821669989 | 2.6237082793 | -1.6267329478 | 0.000394154325503377 |
| WBGene00004 | rps-23 | -0.5358477568 | 7.6000838970 | -1.8391539485 | 0.000145937036737947 |
| WBGene00004 | rps-24 | -0.1467249447 | 0.0302477597 | -1.2249248329 | 0.0161451716418977 |
| WBGene00004 | rps-25 | -0.5691245147 | 2.0886056703 | -1.7808991830 | 0.000340957907800636 |
| WBGene00004 | rps-26 | -0.4147188594 | 1.6636349457 | -1.5717559979 | 0.000622639997514217 |
| WBGene00004 | rps-27 | -0.4214481707 | 2.2890959325 | -1.6557272939 | 0.000269748600157396 |
| WBGene00004 | rps-28 | -0.3812345578 | 1.1784973439 | -1.7849871467 | 0.00032445021679883 |
| WBGene00004 | rps-29 | -0.3751809554 | 4.2843285834 | -1.7376920320 | 0.000446587150055289 |
| WBGene00004 | rps-30 | -0.3284020336 | 1.0573035318 | -1.4027205985 | 0.00132144705165653 |

| | | | | | |
|---|---|---|---|---|---|
| WBGene00004 | rpt-1 | -0.3232675248 | 2.2434666428 | -0.8602071108 | 0.00723677364366282 |
| WBGene00004 | rpt-5 | -0.5067849047 | 2.8439954747 | -0.9514379329 | 0.0223806327271981 |
| WBGene00004 | rars-2 | -0.9299439249 | 3.3252307092 | -0.9874426635 | 0.00539452957835146 |
| WBGene00004 | sip-1 | -0.3189010699 | 0.0031192105 | -1.3400297289 | 0.00573886886712709 |
| WBGene00004 | skr-1 | -0.9997938584 | 3.4997327317 | -1.2008684631 | 0.000145937036737947 |
| WBGene00004 | skr-2 | -0.6761795588 | 1.2320794765 | -1.1811492515 | 0.00559280336363835 |
| WBGene00004 | smn-1 | -0.7898092908 | 5.6880804928 | -1.3756007072 | 5.62711545793835e-05 |
| WBGene00004 | smo-1 | -0.2214775801 | 2.3217466233 | -0.8953538452 | 0.0398296893164544 |
| WBGene00004 | spd-3 | -0.2982160428 | 0.0092015218 | -0.9526573477 | 0.0423296157548056 |
| WBGene00004 | spe-5 | -1.0427953308 | 3.6020906321 | -1.1558171363 | 0.0128213883420574 |
| WBGene00005 | F26F4.8 | -0.8052724264 | 3.1341042812 | -1.4596761892 | 0.000153390482255783 |
| WBGene00005 | sars-1 | -0.5383416301 | 2.6355016642 | -0.9773968581 | 0.00491233666167212 |
| WBGene00006 | tac-1 | -0.2314730787 | 1.9301293696 | -1.1175471383 | 0.000972056743744006 |
| WBGene00006 | fut-3 | -1.5561401514 | 3.5700372990 | -2.5119906050 | 3.38313618532338e-16 |
| WBGene00006 | ant-1.1 | -0.4573381921 | 4.2025527253 | -1.5295122736 | 0.000121181955763698 |
| WBGene00006 | tag-115 | -0.4556054970 | 2.1992705782 | -1.3165816197 | 0.000549694814527833 |
| WBGene00006 | acl-3 | -0.5669494652 | 4.8631364744 | -1.1632204035 | 0.00128421156701525 |
| WBGene00006 | thk-1 | -0.1926838722 | 0.0064046603 | -0.7022660407 | 0.0485777920031503 |
| WBGene00006 | ubc-6 | -0.6359945834 | 4.8006338081 | -1.2214483782 | 0.00497240446106148 |
| WBGene00006 | ubc-9 | -0.5475199363 | 1.3718059033 | -1.1816026646 | 0.000549694814527833 |
| WBGene00006 | ubc-18 | -0.6783898994 | 1.1151895302 | -1.0175219274 | 0.00885639537461736 |
| WBGene00006 | ubh-2 | -0.8049419798 | 6.6138042703 | -1.4292058231 | 0.0002890451114692 |
| WBGene00006 | ubh-3 | -0.9403449242 | 4.0114262131 | -1.3558415712 | 5.80387754337262e-05 |
| WBGene00006 | ubl-1 | -0.3168629102 | 7.3415218088 | -1.4030656053 | 0.00205184539342337 |
| WBGene00006 | ubq-2 | -0.3981012661 | 8.2886201643 | -1.5201320652 | 0.00118945466690309 |
| WBGene00006 | ulp-3 | -0.5316494095 | 2.1587873524 | -1.0867616093 | 0.0485202274827325 |
| WBGene00006 | unc-54 | -0.5328330847 | 0.0401812736 | 1.1335144891 | 0.0176109583884111 |
| WBGene00006 | vig-1 | -0.2959724087 | 1.2492863828 | -1.0832350834 | 0.000649083160055533 |
| WBGene00007 | mdt-8 | -0.4194669582 | 3.3802691479 | -0.8078610486 | 0.0335837708426836 |
| WBGene00007 | mdt-18 | -0.6825107349 | 1.0480560598 | -1.0422739955 | 0.0115552719636781 |
| WBGene00007 | mdt-22 | -0.6541662931 | 2.0807953530 | -1.1897511660 | 0.000179216214333118 |
| WBGene00007 | mdt-29 | -0.4446348723 | 1.2199995383 | -0.8013734554 | 0.0452974243984957 |
| WBGene00007 | B0001.5 | -0.3419594194 | 3.1178245047 | -1.0156659540 | 0.00474556665655936 |
| WBGene00007 | trx-2 | -2.0887552798 | 7.5123985806 | -1.7771565630 | 1.27082934119061e-05 |
| WBGene00007 | B0035.3 | -0.4005754857 | 5.7867479690 | -0.8962216851 | 0.00942429983180476 |
| WBGene00007 | acly-2 | -0.7281197961 | 2.0586707836 | -0.7133770607 | 0.0365753756603351 |
| WBGene00007 | C05C10.2 | -0.9433879290 | 1.1803831387 | -0.9168286285 | 0.00718173560036619 |
| WBGene00007 | cdc-48.1 | -0.1687610200 | 7.2998564705 | -0.7739779846 | 0.0296206393086034 |

| | | | | | |
|---|---|---|---|---|---|
| WBGene00007 | algn-6 | -0.7971684283 | 6.803414071 | -1.0952687003 | 0.00640912669776512 |
| WBGene00007 | pptr-2 | -0.737522535 | 2.686101588 | -1.0430103727 | 0.00100173661445288 |
| WBGene00007 | C14B1.8 | -0.3155857172 | 1.763559452 | -1.0902775933 | 0.0021394879293794 |
| WBGene00007 | C18D11.3 | -0.119501102 | 0.043083692 | -0.920056041 | 0.040615685318781 |
| WBGene00007 | tomm-40 | -0.5264361143 | 1.725686983 | -0.7628456574 | 0.033754881545758 |
| WBGene00007 | tram-1 | -0.807693891 | 2.217290624 | -1.1239410128 | 0.0093802098399202 |
| WBGene00007 | C24F3.2 | -1.4858708171 | 7.286320784 | -1.9443490351 | 6.32866611528839e-08 |
| WBGene00007 | qns-1 | -1.2266151514 | 2.535269209 | -1.396569400 | 1.66278899885084e-05 |
| WBGene00007 | clec-87 | -0.6356347213 | 1.131384657 | -1.1714386254 | 0.00501386450942347 |
| WBGene00007 | mrpl-34 | -0.190403415 | 0.003913219 | -0.9012533153 | 0.0398296893164544 |
| WBGene00007 | C25G4.2 | -0.441247743 | 1.694588003 | -1.407944148 | 0.000770969507008345 |
| WBGene00007 | spe-44 | -1.2522617747 | 1.864671946 | -1.4861207563 | 0.000651782091240334 |
| WBGene00007 | C26D10.3 | -0.7826703895 | 1.218165554 | -1.7833490351 | 0.000903237471222072 |
| WBGene00007 | C27B7.5 | -0.4372256134 | 8.579153508 | -1.008114718 | 0.00677370412473355 |
| WBGene00007 | tofu-1 | -0.3059916797 | 1.813112900 | -0.977924971 | 0.00449720293366652 |
| WBGene00007 | C29A12.1 | -0.9908218767 | 1.180930649 | -1.3729037361 | 3.54156571913312e-05 |
| WBGene00007 | C30H6.9 | -0.7039602655 | 7.752751516 | -1.1431172231 | 0.00636986770451487 |
| WBGene00007 | C35A5.8 | -0.7765746188 | 5.667198569 | -0.9112682975 | 0.0100151084487745 |
| WBGene00008 | sfxn-1.4 | -1.7512250898 | 7.498696631 | -2.8136619861 | 0.000116830216264568 |
| WBGene00008 | C49C3.7 | -1.1269090558 | 1.419218156 | -1.4695566147 | 0.000353928006283995 |
| WBGene00008 | ril-1 | -0.4906487918 | 4.390982728 | -0.842255779 | 0.033754881545758 |
| WBGene00008 | C55A6.1 | -0.9387095893 | 3.207127031 | -1.6594352234 | 1.75400422059934e-08 |
| WBGene00008 | C56A3.4 | -0.681182332 | 3.024101924 | -1.0556104617 | 0.00103219645283455 |
| WBGene00008 | D1054.3 | -0.6038385572 | 2.716422867 | -1.7361372844 | 3.07788940615176e-09 |
| WBGene00008 | F01G4.5 | -0.4388919831 | 1.353898626 | -1.3745725484 | 0.00289114599560019 |
| WBGene00008 | F02E9.5 | -0.7619556453 | 4.844036005 | -1.726582445 | 1.46519193149496e-05 |
| WBGene00008 | gfat-1 | -0.5611847781 | 2.386749316 | -1.2153827655 | 0.00319524488216138 |
| WBGene00008 | F07H5.10 | -0.688572903 | 1.174890582 | -1.3157201525 | 0.000124181031519663 |
| WBGene00008 | dsb-1 | -0.578313194 | 8.928742810 | -0.772280032 | 0.0463171456168638 |
| WBGene00008 | F10B5.2 | -0.3811282175 | 5.830274265 | -0.953894727 | 0.0258679652651789 |
| WBGene00008 | pch-2 | -1.5270260432 | 7.480997252 | -1.9511252445 | 1.66367469370274e-07 |
| WBGene00008 | dylt-1 | -0.3437059623 | 1.164972901 | -0.905452654 | 0.013608461639879 |
| WBGene00008 | eef-1G | -0.3998451517 | 2.259772002 | -1.5021924325 | 0.000100727501279383 |
| WBGene00008 | elli-1 | -0.435563414 | 2.687862278 | -0.841594797 | 0.0154523121407801 |
| WBGene00009 | hint-1 | -0.591527680 | 1.011396001 | -0.8744866967 | 0.0353087416934608 |
| WBGene00009 | pfd-6 | -0.3224391163 | 1.354337310 | -1.229607152 | 0.000322350607850862 |
| WBGene00009 | gfat-2 | -0.4594620784 | 5.102951905 | -1.0754912597 | 0.00643204948490271 |
| WBGene00009 | F22B5.10 | -0.269176346 | 1.890164347 | -0.843491255 | 0.0169233036117202 |

| | | | | | |
|---|---|---|---|---|---|
| WBGene00009 | F22D6.2 | -0.2892825258 | 3.1296373623 | -0.695002632 | 0.0437544571524835 |
| WBGene00009 | F23B12.4 | -1.1894529641 | 3.6820746903 | -2.1880784225 | 0.0456233515442048 |
| WBGene00009 | ndk-1 | -0.564115442 | 8.9133529412 | -1.6848575933 | 0.000205564210550367 |
| WBGene00009 | F25H9.7 | -0.5492533349 | 1.9798320856 | -1.2691994664 | 0.00124982886905589 |
| WBGene00009 | cox-7C | -0.7851126825 | 6.0010663306 | -1.072660297 | 0.0103104356701034 |
| WBGene00009 | F27D4.1 | -1.2751067877 | 1.6202251059 | -1.2129652291 | 0.000133307543909504 |
| WBGene00009 | F30F8.1 | -0.5432579572 | 9.4971124560 | -1.1101663233 | 0.0161518182402066 |
| WBGene00009 | F31C3.3 | -0.7318656297 | 2.0308078699 | -1.0396672613 | 0.00379708600337224 |
| WBGene00009 | psf-2 | -0.6881635577 | 4.6904891888 | -1.6006291136 | 5.23344157137434e-05 |
| WBGene00009 | F32A7.4 | -0.8973804502 | 3.1668106536 | -1.3305551413 | 0.000340957907800636 |
| WBGene00009 | tbc-3 | -0.9517081694 | 1.3024874677 | -1.1877037643 | 0.00164273934511792 |
| WBGene00009 | F32D8.5 | -0.7103876404 | 1.6847515716 | -1.7315179268 | 2.78971557688454e-07 |
| WBGene00009 | F32D8.14 | -0.4001241104 | 2.4065931368 | -1.0042711448 | 0.00355483757279993 |
| WBGene00009 | marb-1 | -0.6808749329 | 3.7244325448 | -1.0308646768 | 0.0184425133577051 |
| WBGene00009 | F35G12.7 | -1.2718197184 | 2.0309198654 | -1.758355418 | 9.07077462671283e-07 |
| WBGene00009 | F40F8.1 | -0.6557124468 | 4.7605435016 | -1.1572065423 | 0.000450766117250963 |
| WBGene00009 | xbx-6 | -0.9027417787 | 5.7912387622 | -1.5386183602 | 3.24555459924065e-06 |
| WBGene00009 | aagr-3 | -0.4835563219 | 1.0632970816 | -0.8435984851 | 0.02598056971727 |
| WBGene00009 | F43G6.3 | -0.7887784615 | 4.6879683899 | -2.0812738597 | 1.74603546934847e-07 |
| WBGene00009 | egg-3 | -0.3666296442 | 5.9766185642 | 0.7623823118 | 0.0432109890811548 |
| WBGene00009 | F44G4.1 | -0.3344606479 | 7.8765707121 | -0.8723923033 | 0.0124380346774357 |
| WBGene00009 | F44G4.2 | -0.4717225958 | 1.9814144362 | -1.1359373822 | 0.00123881112648002 |
| WBGene00009 | F44G4.3 | -0.9523312889 | 2.5569131256 | -1.4929426417 | 0.00815576152222132 |
| WBGene00009 | F46B6.12 | -0.4203514805 | 5.0941187521 | -0.8707885297 | 0.0262867713947236 |
| WBGene00009 | F49C12.9 | -0.7833726172 | 4.7990975735 | -1.8683892351 | 5.83330330439036e-11 |
| WBGene00009 | F49C12.11 | -0.3270955994 | 2.2045576758 | -1.1958323936 | 0.00294492304896226 |
| WBGene00009 | F49C12.12 | -0.5668485735 | 2.2094101864 | -1.0798222647 | 0.0331096221507528 |
| WBGene00009 | vha-17 | -0.8833398887 | 2.5197517846 | -1.5558250615 | 0.000607456725515455 |
| WBGene00009 | gcsh-2 | -1.1540309769 | 3.8459919775 | -2.4526437322 | 1.66714033278636e-13 |
| WBGene00009 | kbrl-1 | -1.0341766245 | 1.1934406282 | -1.3629047578 | 0.0089347942925171 |
| WBGene00009 | ogr-2 | -2.3004264996 | 1.2169142485 | -2.1859431606 | 0.000213129492314089 |
| WBGene00010 | bcs-1 | -0.3022414348 | 1.1384254462 | -0.989389033 | 0.00506681649534184 |
| WBGene00010 | F54D5.7 | -0.4828423207 | 1.5149573430 | -0.7507310804 | 0.0461749028997023 |
| WBGene00010 | vha-14 | -0.2536654791 | 2.7115152254 | -0.6915195339 | 0.0454929174258304 |
| WBGene00010 | F57A10.4 | -0.8859906303 | 4.2973897456 | -1.1762634227 | 0.0281160137391156 |
| WBGene00010 | pot-2 | -0.7735719677 | 2.1126376028 | -1.3402216854 | 0.000958347340252515 |
| WBGene00010 | F57F5.1 | -1.0765404398 | 0.0003458020 | 2.0650189582 | 0.0364512076140284 |
| WBGene00010 | F58A4.2 | -1.8286745478 | 6.5292526081 | -2.7138007267 | 1.18152069329156e-12 |

| | | | | | |
|---|---|---|---|---|---|
| WBGene00010 | F58A4.6 | -0.611577384C | 7.5806236413[illegible] | -1.3028290016 | 0.000935725912547021 |
| WBGene00010 | rbm-17 | -0.2390840649 | 0.0002616896[illegible] | -0.8905416563 | 0.0332656119558407 |
| WBGene00010 | F58H1.3 | -0.5631400184 | 2.8489624451[illegible] | -1.289809878C | 0.00450375172201545 |
| WBGene00010 | idh-1 | -0.8078370198 | 2.1114896102[illegible] | -0.8880986145 | 0.00895635473259128 |
| WBGene00010 | H19N07.3 | -0.3512539291 | 0.0226437656[illegible] | -1.1886058128 | 0.0161451716418977 |
| WBGene00010 | atp-1 | -0.4419912086 | 8.6622808809[illegible] | -1.0883778038 | 0.00437687905729372 |
| WBGene00010 | enu-3.6 | -1.0323672015 | 9.9527439227[illegible] | -2.0450308938 | 4.58962259640861e-05 |
| WBGene00010 | K01G5.8 | -0.2381704544 | 0.0035929059[illegible] | -0.918496334C | 0.0223151767303369 |
| WBGene00010 | rack-1 | -0.3811215248 | 9.8058012413[illegible] | -1.4001266298 | 0.00360746772975649 |
| WBGene00010 | cdc-48.3 | -0.8701636974 | 2.5796956277[illegible] | -1.2759944052 | 0.00545753427217956 |
| WBGene00010 | ath-1 | -1.0755450059 | 1.3001210330[illegible] | -1.3072188785 | 0.00092440542526763 |
| WBGene00010 | vacl-14 | -1.3365613875 | 3.6261531674[illegible] | -1.0634259299 | 0.0024790258954136 |
| WBGene00010 | scbp-2 | -0.9931831887 | 3.2997836661[illegible] | -1.7971311671 | 0.0118123134770021 |
| WBGene00010 | dut-1 | -0.3659860409 | 8.4687219177[illegible] | -0.7392087473 | 0.0372327044948143 |
| WBGene00010 | K07C5.3 | -0.3888306999 | 3.8060785956[illegible] | -0.7457148888 | 0.0412620104466134 |
| WBGene00010 | dbn-1 | -0.4802692674 | 6.7113012021[illegible] | -0.7674629562 | 0.0286823030911631 |
| WBGene00010 | K08E3.5 | -0.5623750080 | 8.5874517363[illegible] | -0.7943690759 | 0.0336778002262491 |
| WBGene00010 | K08E4.2 | -0.868062353C | 1.1549137611[illegible] | -1.6972746438 | 0.000111854824934051 |
| WBGene00010 | K08F4.3 | -0.4515678516 | 1.9515916629[illegible] | -1.7388219232 | 2.26635683089548e-08 |
| WBGene00010 | aipl-1 | -1.2290300813 | 2.1281209507[illegible] | -1.4586606714 | 1.04848973078758e-06 |
| WBGene00010 | trpp-6 | -0.1901924785 | 0.0112935315[illegible] | -0.7493638886 | 0.0423961764505718 |
| WBGene00010 | sodh-1 | -1.2261338548 | 0.0004697212[illegible] | -2.9936148734 | 0.00157921715493894 |
| WBGene00010 | M01F1.3 | -0.2785660579 | 1.4847781986[illegible] | -1.0441231872 | 0.0119879695899209 |
| WBGene00010 | M110.3 | -0.5447498729 | 5.9766674736[illegible] | -1.1732097581 | 0.0128272093808894 |
| WBGene00010 | R01H10.7 | -0.6394986035 | 7.2681308568[illegible] | -1.341812921C | 0.00168516076425543 |
| WBGene00011 | R05H5.3 | -0.5012004694 | 4.7292367306[illegible] | -1.2135240613 | 0.00979700011909909 |
| WBGene00011 | R05H5.4 | -1.8168783661 | 5.5833775605[illegible] | -1.6145801075 | 0.000653977949136587 |
| WBGene00011 | R05H5.5 | -1.9201798544 | 6.0076349182[illegible] | -1.8614192293 | 0.0027895859221073 |
| WBGene00011 | arrd-13 | -1.2641694458 | 1.1958960015[illegible] | -1.1550032912 | 0.00273177042703495 |
| WBGene00011 | cpg-3 | -0.4146628504 | 1.4738771280[illegible] | -1.5332574385 | 0.000352530084109221 |
| WBGene00011 | adsl-1 | -0.399594539C | 7.5167303738[illegible] | -0.7541825795 | 0.0473193699012478 |
| WBGene00011 | ztf-15 | -0.8017513408 | 5.9570924934[illegible] | -0.8685533008 | 0.0219922527615037 |
| WBGene00011 | R07H5.8 | -1.1497821624 | 8.4958040996[illegible] | -1.3214051167 | 8.44127148142459e-06 |
| WBGene00011 | rbm-3.2 | -0.6690063715 | 3.2936958100[illegible] | -1.776768911C | 0.000457131841034829 |
| WBGene00011 | gkow-1 | -0.6057732963 | 2.6572062572[illegible] | -0.744982576C | 0.0467334672360067 |
| WBGene00011 | R53.4 | -0.2202769992 | 1.1476011525[illegible] | -0.9715645943 | 0.0332656119558407 |
| WBGene00011 | R74.8 | -0.3585322177 | 2.4696934661[illegible] | -0.7525830315 | 0.0325318905054985 |
| WBGene00011 | allo-1 | -1.1335785511 | 6.0357193278[illegible] | -1.3826643408 | 4.49779203958323e-05 |

| | | | | | |
|---|---|---|---|---|---|
| WBGene00011 | R186.3 | -0.4641227786 | 3.0220018998 | -0.9622262687 | 0.00989402234545058 |
| WBGene00011 | hrde-2 | -0.4292890068 | 3.2835665795 | -1.1539056107 | 0.00103219645283455 |
| WBGene00011 | T03F6.3 | -0.8276579464 | 3.3238226568 | -1.5210589026 | 1.6216592557174e-06 |
| WBGene00011 | T05B9.1 | -0.1782414408 | 3.3033027589 | -0.7959909162 | 0.0243138899128504 |
| WBGene00011 | rmd-1 | -0.4135262556 | 3.1138904300 | -1.4213679549 | 9.01310911531528e-06 |
| WBGene00011 | pdha-1 | -0.5212875202 | 3.7102180403 | -1.0025024889 | 0.00292542072350632 |
| WBGene00011 | gpcp-2 | -2.0207236784 | 0 | -1.7656417776 | 3.69496504494446e-08 |
| WBGene00011 | cchl-1 | -0.3563622074 | 9.8647661184 | -0.8058468736 | 0.0204303981800394 |
| WBGene00011 | umps-1 | -1.1072567848 | 1.7410510820 | -1.5608666892 | 9.55535219717875e-06 |
| WBGene00011 | sds-22 | -0.5000434351 | 6.0304350408 | -1.3680361326 | 2.52583743459589e-05 |
| WBGene00011 | T09A5.15 | -0.5000619919 | 0.0012324174 | -1.9719376368 | 0.00897654577886356 |
| WBGene00011 | T10C6.6 | -0.7322621015 | 6.7428429512 | -1.0484980711 | 0.00511247369619112 |
| WBGene00011 | T12G3.4 | -1.3815803598 | 1.0078264132 | -2.1155394815 | 2.01871975550886e-06 |
| WBGene00011 | mrps-18B | -0.8432093915 | 4.3479447137 | -0.9892989212 | 0.0122456032813657 |
| WBGene00011 | gtf-2H2C | -0.4152325148 | 1.3473577757 | -0.8210022570 | 0.0257935778433515 |
| WBGene00011 | T20D3.5 | -0.8133543959 | 1.3055734811 | -0.9979583334 | 0.0154518252417962 |
| WBGene00011 | mrpl-50 | -0.1414246863 | 0.0168981478 | -0.9845180643 | 0.00739247315611174 |
| WBGene00011 | erh-1 | -0.1852913826 | 0.0031173682 | -0.7940099336 | 0.0243138899128504 |
| WBGene00011 | ctf-8 | -0.3930476357 | 0.0003215992 | -1.2087228110 | 0.000207815563577541 |
| WBGene00012 | dkf-2 | -0.2267166726 | 0.0001314799 | 0.9580136230 | 0.00924849189475974 |
| WBGene00012 | ebp-2 | -0.7893579946 | 1.3543078897 | -0.9717962945 | 0.0166422659830224 |
| WBGene00012 | adbp-1 | -0.9405922344 | 2.3656215931 | -1.1864851469 | 0.0161451716418977 |
| WBGene00012 | W01D2.1 | -0.3000410725 | 5.0640328269 | -1.5812734808 | 0.00157921715493894 |
| WBGene00012 | osta-3 | -0.3539266365 | 6.7626162963 | -0.8870789312 | 0.0100767602737566 |
| WBGene00012 | W01G7.4 | -0.3942368688 | 3.2940223166 | -1.2025145249 | 0.000508560303415O5 |
| WBGene00012 | W02B12.11 | -0.5908768659 | 3.1636667851 | -0.8567347953 | 0.0243138899128504 |
| WBGene00012 | W07G4.3 | -1.5475617317 | 4.2705213667 | -1.6089383190 | 4.62190058821177e-05 |
| WBGene00012 | Y11D7A.7 | -2.0072527163 | 4.1093964004 | -1.9051573512 | 0.00100173661445288 |
| WBGene00012 | Y17G7B.21 | -0.5094305294 | 4.8549416187 | -1.3984016707 | 0.0269444824229092 |
| WBGene00012 | Y18D10A.11 | -0.4587301103 | 1.2136594903 | -1.4402627358 | 2.51064728013099e-05 |
| WBGene00012 | car-1 | -0.3747943413 | 0.0167989560 | -0.8464287280 | 0.0291838371309107 |
| WBGene00012 | cec-7 | -0.3695392846 | 1.2427556527 | -1.1238800167 | 0.00276905205630985 |
| WBGene00012 | Y39B6A.37 | -0.2811130407 | 6.8587149274 | -0.8752206036 | 0.0296206393086034 |
| WBGene00012 | Y41E3.8 | -0.5475481823 | 8.6387744843 | -1.1150674933 | 0.0168205811972573 |
| WBGene00012 | Y43E12A.3 | -0.8635984119 | 3.4015784651 | -1.7978704739 | 4.71144492096621e-05 |
| WBGene00012 | mrpl-20 | -0.1421943328 | 0.0475653611 | -1.0816017030 | 0.0336672285262766 |
| WBGene00013 | cept-2 | -1.6047156024 | 1.7986228230 | -1.5998202900 | 0.00217151044409565 |
| WBGene00013 | Y49A3A.3 | -0.9183725840 | 8.3184315627 | -0.9587082900 | 0.0369209930932115 |

| | | | | | |
|---|---|---|---|---|---|
| WBGene00013 | Y53H1B.2 | -0.6233469971 | 0.0284400933 | 2.5840193417 | 1.66488265312191e-05 |
| WBGene00013 | Y54G9A.5 | -0.8114311843 | 1.0343918552 | -1.6189945068 | 2.34011410893036e-07 |
| WBGene00013 | bub-3 | -0.4485346493 | 2.1511795058 | -0.8244226570 | 0.0170958384865374 |
| WBGene00013 | Y57G11C.8 | -1.0192290657 | 7.2160233597 | -2.0966361873 | 0.000422686938373154 |
| WBGene00013 | Y57G11C.36 | -2.4713082915 | 1.4371164363 | -2.6979641932 | 2.59794690993153e-14 |
| WBGene00013 | fars-2 | -0.5171478603 | 1.5557331811 | -1.1215995332 | 0.00139555742596482 |
| WBGene00013 | Y62E10A.13 | -0.3134633615 | 3.7179187708 | -0.8620963397 | 0.0305254502373799 |
| WBGene00013 | ule-5 | -1.6094132039 | 0.0123245118 | 6.2857815922 | 1.80372556890977e-07 |
| WBGene00013 | Y67H2A.2 | -0.2763433167 | 3.2053056244 | -1.3185028101 | 0.000684897301178945 |
| WBGene00013 | micu-1 | -0.6657002657 | 7.5385929469 | -0.9693980331 | 0.0178115489770295 |
| WBGene00013 | kdp-1 | -0.1243183307 | 0.0497230328 | -0.9519931418 | 0.0155184979937502 |
| WBGene00013 | Y75B8A.18 | -0.5579031233 | 2.5032377908 | -1.2262224486 | 0.000205564210550367 |
| WBGene00013 | bath-36 | -0.3370610028 | 0.0061809049 | -1.3187364468 | 0.00134096442093109 |
| WBGene00013 | Y87G2A.18 | -0.9270177866 | 4.9290863993 | -1.0983711178 | 0.0283902055649011 |
| WBGene00013 | Y105E8A.14 | -0.1283727711 | 0.0324539487 | -1.0867969912 | 0.0121827370881007 |
| WBGene00013 | Y106G6H.14 | -0.2292033737 | 0.0056123103 | -0.7647056501 | 0.0357022951838369 |
| WBGene00013 | ZK287.7 | -0.5311457715 | 2.2803548517 | -0.9565891926 | 0.0261894168811472 |
| WBGene00014 | riok-3 | -0.4536584092 | 6.6921295050 | -0.9974083281 | 0.00506681649534184 |
| WBGene00014 | ZK643.2 | -0.7944352241 | 9.0383093569 | -0.9629729015 | 0.0110532723345955 |
| WBGene00014 | gdh-1 | -0.6325820352 | 5.2541991511 | -0.7441263183 | 0.0445570152615297 |
| WBGene00014 | chpf-1 | -0.4935422765 | 3.9537966699 | -1.0286710904 | 0.00598048819140576 |
| WBGene00014 | ZK856.11 | -0.5018604379 | 1.5825793629 | -1.2363233887 | 0.00177514678549764 |
| WBGene00014 | clpp-1 | -0.7412954311 | 6.7615649591 | -1.1187636939 | 0.00276901369474864 |
| WBGene00014 | ZK1010.2 | -0.1647328013 | 0.0164627960 | -0.9916063361 | 0.00474556665655936 |
| WBGene00014 | ZK1320.9 | -0.9668023110 | 1.2570705420 | -0.9271411037 | 0.0145923950011764 |
| WBGene00014 | C06A1.4 | -0.2676950401 | 8.6545931295 | -0.7837600362 | 0.0200811112092543 |
| WBGene00014 | F56D5.4 | -0.8396566090 | 8.3449638413 | -1.5403556601 | 0.0261252586228313 |
| WBGene00014 | ZK637.6 | -1.3613854990 | 1.3625881610 | -1.9055148611 | 1.78334142028818e-06 |
| WBGene00015 | B0205.8 | -0.6276551006 | 0.0001122942 | -0.9306068179 | 0.0312371879423459 |
| WBGene00015 | B0238.9 | -0.8303227933 | 2.4170906324 | -1.0098649568 | 0.00530316637230017 |
| WBGene00015 | natc-2 | -0.4078630107 | 9.1325745551 | -0.9026140805 | 0.00908195216272628 |
| WBGene00015 | cpg-2 | -0.4851018241 | 7.6793526966 | -1.0898282627 | 0.0465138372032017 |
| WBGene00015 | B0304.2 | -0.6894900423 | 3.7581304138 | -1.0899299474 | 0.00216338447887046 |
| WBGene00015 | arle-14 | -0.3742523276 | 3.7258906575 | -0.8677645976 | 0.0179408245694232 |
| WBGene00015 | B0336.13 | -0.2505110654 | 0.0004146922 | -1.1111787063 | 0.000871900460474218 |
| WBGene00015 | cdc-26 | -0.3326188450 | 5.9579735573 | -0.8938169368 | 0.0242354245041834 |
| WBGene00015 | C01B12.8 | -1.3002002397 | 1.9295358771 | -1.6039153104 | 7.58581909918408e-05 |
| WBGene00015 | ddi-1 | -0.4491623878 | 4.7370211059 | -0.9044096528 | 0.00807211411071431 |

| | | | | | |
|---|---|---|---|---|---|
| WBGene00015 | C01G8.1 | -0.2532326861 | 9.0129725192 | -1.2687337757 | 3.31185720168559e-05 |
| WBGene00015 | ivd-1 | -0.7684977694 | 2.0228172663 | -0.8806047842 | 0.0129349286410967 |
| WBGene00015 | snpn-1 | -0.5810797893 | 4.3510958734 | -1.0255233726 | 0.0106033449406399 |
| WBGene00015 | C04E6.11 | -0.6237402425 | 3.4571785473 | -0.7735318417 | 0.0481286646478953 |
| WBGene00015 | C06A5.6 | -0.5771773245 | 3.9786672047 | -1.0700424020 | 0.00103219645283455 |
| WBGene00015 | sams-3 | -0.2049445593 | 0.0003992857 | -1.5625705931 | 6.3694466229515e-07 |
| WBGene00015 | C07H6.2 | -1.0320199470 | 6.0331066472 | -1.4063290517 | 0.0337368089204015 |
| WBGene00015 | C09E7.7 | -0.7844950501 | 6.3471138734 | -1.4545368804 | 0.00841726100453489 |
| WBGene00015 | C11D2.7 | -0.8427442863 | 8.8913192249 | -1.1930563039 | 0.000903237471222072 |
| WBGene00015 | C13F10.5 | -0.2361045210 | 0.0024699689 | -0.9327570622 | 0.0189396424832857 |
| WBGene00015 | C16A11.3 | -0.2826908479 | 0.0001262755 | -0.9443555945 | 0.0303131756710447 |
| WBGene00015 | C18F10.7 | -1.3677162581 | 8.8355375992 | -1.2492222052 | 0.000111676552334899 |
| WBGene00016 | mdt-26 | -1.0359444605 | 9.5858728167 | -1.2617699943 | 0.00459640898935717 |
| WBGene00016 | ift-43 | -1.0658862161 | 9.3487214147 | -2.6902234421 | 0.00200107317870346 |
| WBGene00016 | elpc-4 | -0.4299420326 | 8.3028137838 | -1.0403901053 | 0.0069037543072462 |
| WBGene00016 | mesp-1 | -0.6004598443 | 8.4517580399 | -1.1396533208 | 0.00413441580878045 |
| WBGene00016 | mob-4 | -0.7312094227 | 1.6641796538 | -0.7775439472 | 0.0275903896523023 |
| WBGene00016 | hpo-27 | -1.4699032366 | 3.7487844279 | -0.8769432641 | 0.0303866056542812 |
| WBGene00016 | C35D10.5 | -1.0231797336 | 9.8996408826 | -1.0187475154 | 0.0145584411311045 |
| WBGene00016 | C35D10.10 | -0.8315724446 | 3.0235153332 | -1.2339766910 | 0.00102379428314494 |
| WBGene00016 | C37A2.7 | -0.2553533484 | 2.9098997045 | -1.4463545132 | 0.00256792105080881 |
| WBGene00016 | C42C1.8 | -0.7196205922 | 4.3388568296 | -1.3954806364 | 1.97337966950321e-05 |
| WBGene00016 | C42C1.13 | -0.4259750861 | 4.1393904798 | -0.9520135187 | 0.0262867713947236 |
| WBGene00016 | arch-1 | -0.7781922496 | 9.6632862755 | -2.0195852158 | 1.19273295195317e-06 |
| WBGene00016 | C45G9.2 | -0.4135397183 | 2.1988532744 | -0.9232624068 | 0.0248382742211052 |
| WBGene00016 | C50C3.1 | -0.7597625340 | 3.4621437273 | -0.9347972731 | 0.0139236874261562 |
| WBGene00016 | sucg-1 | -0.8704790301 | 1.5895386659 | -0.7982933948 | 0.0490785725796147 |
| WBGene00016 | C53H9.2 | -0.2482098824 | 0.0004330242 | -1.5186378033 | 0.000168371087637383 |
| WBGene00016 | madf-6 | -0.8624516816 | 2.2368226433 | -1.4300746172 | 0.000549694814527833 |
| WBGene00016 | C56C10.7 | -0.3181187517 | 6.0355191396 | -0.7539504091 | 0.0301985736874914 |
| WBGene00016 | epg-5 | -0.2742774638 | 0.0065683188 | -1.2074408635 | 0.0050242906408362 |
| WBGene00017 | D1007.10 | -0.6056355356 | 1.6091862858 | -1.1086419909 | 0.0155992455340687 |
| WBGene00017 | D2024.5 | -0.2379770213 | 1.4269042861 | -0.7611120184 | 0.0353087416934608 |
| WBGene00017 | glrx-3 | -0.2472883938 | 5.8775078037 | -0.9713493614 | 0.00263774656986395 |
| WBGene00017 | ppfr-2 | -0.6506856480 | 2.3460285903 | -0.9657036521 | 0.0093802098399202 |
| WBGene00017 | maco-1 | -1.5346000423 | 6.0589154009 | -1.5100107941 | 0.000147132736403807 |
| WBGene00017 | D2096.7 | -0.4206611208 | 5.4487988672 | -1.2212749808 | 0.00630463673704035 |
| WBGene00017 | cyc-2.1 | -0.4758094814 | 1.1393820378 | -1.1052489013 | 0.00754173239372364 |

| | | | | | |
|---|---|---|---|---|---|
| WBGene00017 | tofu-6 | -0.5172414119 | 2.4784098253 | -0.8559678048 | 0.00738368517951859 |
| WBGene00017 | EEED8.3 | -2.5823481574 | 1.7712071507 | -1.9747339387 | 2.44286985405309e-11 |
| WBGene00017 | EEED8.13 | -3.7626922224 | 5.3810701440 | -3.0282752567 | 0.000209131983178942 |
| WBGene00017 | EEED8.14 | -4.0138917672 | 0 | -2.3474867547 | 2.99239547764308e-08 |
| WBGene00017 | E_BE45912.2 | -0.4759190904 | 0.0003151156 | -1.3762077954 | 0.0248342126466234 |
| WBGene00017 | aldo-2 | -0.5223016864 | 4.2009827982 | -1.5184324892 | 1.0460442912599e-06 |
| WBGene00017 | F07F6.7 | -1.1689430578 | 6.2399129713 | -2.0241874922 | 0.00284025348926687 |
| WBGene00017 | F10E9.7 | -0.9259628709 | 3.2072592882 | -1.0757615958 | 0.0129154452124318 |
| WBGene00017 | F14D2.11 | -1.2352313426 | 6.1341969984 | -1.5947267373 | 0.00500289730536609 |
| WBGene00017 | F14D2.14 | -0.4459178221 | 0.0002452130 | -1.2144385871 | 0.0299521577536132 |
| WBGene00017 | pid-1 | -0.5618108612 | 6.7743509363 | -1.1324845978 | 0.00180557581903888 |
| WBGene00017 | tofu-2 | -0.6968863461 | 1.0073263270 | -1.4932528361 | 1.03073966602003e-05 |
| WBGene00017 | polk-1 | -0.5203977585 | 1.7036326236 | -0.8549399543 | 0.017057526997178 |
| WBGene00017 | F23H11.2 | -0.8565466207 | 5.5269953287 | -1.2398302825 | 0.0116427887769126 |
| WBGene00017 | sucl-2 | -0.4717186082 | 2.1162413262 | -0.8287484405 | 0.0152204464533136 |
| WBGene00017 | crls-1 | -0.5325977458 | 1.2156240391 | -1.0053996340 | 0.00656268717028767 |
| WBGene00017 | F25B5.5 | -0.3598070224 | 1.6950730382 | -1.0978318275 | 0.00126254757190248 |
| WBGene00017 | F25B5.6 | -0.7193508970 | 6.8035957872 | -1.1551945298 | 0.00290486678286092 |
| WBGene00017 | F26F4.6 | -0.2198832675 | 0.0019356570 | -0.8918022728 | 0.0139440228697523 |
| WBGene00017 | F26F4.9 | -0.4743290257 | 3.8445976997 | -1.2188659593 | 0.000390481151327125 |
| WBGene00017 | F26F4.12 | -0.1583246121 | 0.0276809189 | -1.2053229015 | 0.000665917845674207 |
| WBGene00017 | kbp-2 | -0.4215871829 | 6.5363924856 | -1.1464823870 | 0.00677370412473355 |
| WBGene00017 | neg-1 | -1.0112136620 | 1.3055679924 | -0.8917007140 | 0.0235556494463057 |
| WBGene00017 | F32D1.7 | -0.6114758747 | 2.0908766677 | -0.9161214487 | 0.0127285525894454 |
| WBGene00017 | cec-4 | -0.8616108182 | 1.8345740239 | -0.9254739856 | 0.02598056971727 |
| WBGene00018 | ztf-28 | -2.6918189450 | 1.5574853044 | -1.3969454271 | 0.000510854549622812 |
| WBGene00018 | F37A4.1 | -0.5118467469 | 2.2721417406 | -0.8827341897 | 0.0162618944506573 |
| WBGene00018 | F37A4.2 | -0.3993221749 | 7.7950641154 | -1.4939265971 | 2.05276210072155e-05 |
| WBGene00018 | mpdu-1 | -0.8863482592 | 1.4476331501 | -1.6911840149 | 5.34559173032201e-05 |
| WBGene00018 | F40A3.3 | -0.7968116548 | 8.7926462320 | -1.0561377905 | 0.00895635473259128 |
| WBGene00018 | mage-1 | -0.5816986056 | 1.0680470461 | -0.8966694168 | 0.0244783720413201 |
| WBGene00018 | F41G3.6 | -0.7286689297 | 1.0226732734 | -1.0637679187 | 0.000618548436802115 |
| WBGene00018 | exos-8 | -0.2774625425 | 5.7862861999 | -1.5089575233 | 1.91616686475813e-06 |
| WBGene00018 | F42A9.8 | -0.8830911502 | 4.8372195160 | -1.3065539445 | 0.0311577370372484 |
| WBGene00018 | F43E2.1 | -0.1755549273 | 0.0390474422 | -0.9943257334 | 0.0342244440990075 |
| WBGene00018 | F44E7.5 | -1.7377625503 | 1.2066239095 | -2.8122259267 | 2.73911497924593e-08 |
| WBGene00018 | skpt-1 | -0.5261001567 | 1.0129536449 | -1.6474786750 | 7.92918249265342e-08 |
| WBGene00018 | oef-1 | -0.2448715984 | 6.4377461919 | -1.0630626866 | 0.00262856802218715 |

| | | | | | |
|---|---|---|---|---|---|
| WBGene00018 | F49E8.6 | -0.4406310050 | 0.0038908557 | -1.1658817004 | 0.0170106070434896 |
| WBGene00018 | F52C12.2 | -0.6207762716 | 2.2250700557 | -1.1206214577 | 0.0106033449406399 |
| WBGene00018 | vha-18 | -1.2016439116 | 2.7034577052 | -1.5341265991 | 6.3694466229515e-07 |
| WBGene00018 | F53F10.2 | -0.7561187290 | 1.1463408225 | -0.9372137102 | 0.011456584182127 |
| WBGene00018 | pst-2 | -0.2137414934 | 0.0021612732 | -0.9129991224 | 0.0465138372032017 |
| WBGene00018 | eef-1B.1 | -0.5565578931 | 8.0229617196 | -1.6739623293 | 0.000112574325495644 |
| WBGene00018 | wago-2 | -0.7553149902 | 5.1300861478 | -0.8695607857 | 0.0430134412966845 |
| WBGene00018 | zhp-1 | -0.6865486532 | 5.1705761591 | -1.0695830344 | 0.0115028859114922 |
| WBGene00018 | mrps-16 | -0.2617870155 | 0.0004314056 | -1.1688860834 | 0.0186127178510814 |
| WBGene00018 | F56D2.2 | -0.6850955922 | 1.6987358451 | -0.8910479806 | 0.0188072328511341 |
| WBGene00019 | F59A7.8 | -2.2129447879 | 3.2823070246 | -2.9887330124 | 9.20261700414605e-12 |
| WBGene00019 | H14A12.3 | -1.7966953316 | 4.6367347521 | -2.8872273145 | 1.27673630128777e-11 |
| WBGene00019 | ahcy-1 | -0.3877513302 | 6.5638795562 | -1.0531435476 | 0.0169441344162188 |
| WBGene00019 | K03B4.1 | -0.4552153373 | 1.5030115417 | -1.0911489177 | 0.00221361802153402 |
| WBGene00019 | K03B4.2 | -0.4717855419 | 1.0740528858 | -0.8832892220 | 0.00803374610352617 |
| WBGene00019 | K04C2.3 | -0.5437330760 | 7.8214175509 | -1.0321290382 | 0.00531767608699746 |
| WBGene00019 | acdh-3 | -0.5455772758 | 4.0136417742 | -0.9139812773 | 0.0145082457050912 |
| WBGene00019 | K06B9.2 | -0.8820112005 | 0.0249916222 | -1.8607686114 | 0.0300271386046165 |
| WBGene00019 | tos-1 | -0.1770434150 | 0.0326837299 | -0.8644775163 | 0.0243138899128504 |
| WBGene00019 | K07E8.7 | -1.5902362944 | 2.1723891267 | -2.0522272036 | 6.53319000782299e-06 |
| WBGene00019 | K07H8.9 | -0.4164960250 | 1.6964361743 | -1.8135996804 | 1.53330988297995e-06 |
| WBGene00019 | K08D12.3 | -0.3688577055 | 0.0026993517 | -1.1797341002 | 0.00136467290642108 |
| WBGene00019 | K09H9.7 | -1.8051930443 | 1.8955680879 | -1.7705279521 | 0.00388591688508462 |
| WBGene00019 | clec-88 | -0.2675263518 | 6.0108546947 | -0.7447068959 | 0.0223806327271981 |
| WBGene00019 | K11D12.12 | -1.2663432387 | 6.6192163892 | -1.4380369434 | 0.0233323892972349 |
| WBGene00019 | K12H4.2 | -0.3979009347 | 8.9391365003 | -1.1561678363 | 0.00933478753881291 |
| WBGene00019 | K12H4.3 | -0.2153549296 | 0.0002550847 | -1.1446190828 | 0.00050285466026984 |
| WBGene00019 | spcs-3 | -0.3580867218 | 4.2986850776 | -1.5472129541 | 1.09514906507189e-06 |
| WBGene00019 | M02B7.2 | -0.5197910489 | 5.4602354559 | -1.0821189290 | 0.0337165143815552 |
| WBGene00019 | algn-14 | -0.2376397582 | 0.0001921181 | -1.2042446637 | 0.0312371879423459 |
| WBGene00019 | egg-2 | -0.4201297815 | 2.2699829141 | -0.7533867050 | 0.0461749028997023 |
| WBGene00019 | R02D3.8 | -1.1659362112 | 1.1678434627 | -1.4744426610 | 0.000824068172363266 |
| WBGene00019 | R02F2.1 | -0.6920139976 | 1.6079060319 | -0.9314457990 | 0.0078656359497159 |
| WBGene00019 | R05F9.6 | -1.1135775774 | 4.7186398914 | -1.2321732561 | 0.000144314865322779 |
| WBGene00019 | sti-1 | -0.7538059190 | 3.0986634845 | -1.2561571441 | 0.00708274462531474 |
| WBGene00020 | R12C12.7 | -1.0369314668 | 3.3745052656 | -1.6336109945 | 0.000123049391069684 |
| WBGene00020 | egg-5 | -0.8737967164 | 1.4896645444 | -1.2313654263 | 0.00503912163069302 |
| WBGene00020 | kbp-1 | -0.7711880060 | 6.0111546462 | -1.2074522070 | 0.00553924671861108 |

| | | | | | |
|---|---|---|---|---|---|
| WBGene00020 | T02G5.7 | -0.2361901302 | 8.9372720838( | -0.7009713638 | 0.044022646510514 |
| WBGene00020 | T02G5.11 | -3.9975466995 | 1.3484377734: | -3.0698517492 | 0.0166430563819311 |
| WBGene00020 | atp-4 | -0.3471443367 | 1.1821120435( | -0.8575003875 | 0.033154286463503 |
| WBGene00020 | tag-261 | -0.1997171901 | 0.0040813730! | -0.9876120596 | 0.0079717549655951 |
| WBGene00020 | T08B2.11 | -2.1672401826 | 5.7006926803: | -3.0029656029 | 1.52254313225766e-08 |
| WBGene00020 | cerk-1 | -0.5368392228 | 6.7634405082: | -0.9949977564 | 0.00268598678908732 |
| WBGene00020 | T12F5.2 | -0.7640267506 | 2.8116559468! | -1.5990052372 | 4.45383950996644e-05 |
| WBGene00020 | ssup-72 | -0.3632443555 | 1.4504837956( | -0.9202512830 | 0.0298449485909544 |
| WBGene00020 | mrps-18.C | -0.2455631598 | 0.0020192569: | -0.9309296633 | 0.031153239395208 |
| WBGene00020 | T14B4.3 | -0.3613826303 | 1.3383810150! | -0.8964511321 | 0.00623810332288005 |
| WBGene00020 | T19H12.2 | -0.4095072133 | 3.5798294672 | -1.0626690956 | 0.00291156215625154 |
| WBGene00020 | mrrf-1 | -1.4154546489 | 8.2064263184! | -1.6801258388 | 0.0212807746566505 |
| WBGene00020 | pygl-1 | -0.9071629588 | 1.2439698522: | -1.1610329024 | 0.000110414404197634 |
| WBGene00020 | T23B3.1 | -0.9847039625 | 8.7068543477! | -1.0588938134 | 0.00351398392465421 |
| WBGene00020 | ucr-2.3 | -0.4799078956 | 7.0212863748: | -0.8018193383 | 0.0373779614830297 |
| WBGene00020 | T24C4.5 | -0.5698066004 | 6.0711399199! | -1.2527880006 | 0.0011759418423118 |
| WBGene00020 | T24H7.3 | -0.4655912422 | 1.2941182879: | -0.8905520342 | 0.0158736979154305 |
| WBGene00020 | T26A5.8 | -0.4569180313 | 7.7299605048! | -1.4971795460 | 4.21085398307813e-05 |
| WBGene00020 | W02D7.6 | -0.7245636682 | 6.8269444828: | -1.6067232274 | 2.55212689430524e-07 |
| WBGene00020 | sna-1 | -0.4807256916 | 1.4989440040: | -1.0187295844 | 0.00212159820835683 |
| WBGene00021 | W03F9.2 | -0.4000957333 | 8.1510657613: | -1.0490169675 | 0.0334676409985324 |
| WBGene00021 | W03G9.2 | -0.7139553255 | 6.2343209601( | -0.9325040228 | 0.0203341362737953 |
| WBGene00021 | W07E6.5 | -0.3087877870 | 0.0322551602! | -1.6974311070 | 1.27082934119061e-05 |
| WBGene00021 | zip-11 | -1.7604287586 | 4.9287971001: | -3.1856886391 | 8.29822266521554e-10 |
| WBGene00021 | cdc-37 | -0.8686190271 | 8.7432378082! | -1.0638616748 | 0.00119903939876447 |
| WBGene00021 | W09B6.4 | -0.4890737669 | 1.9753514855( | -1.5853284451 | 0.0136099493059425 |
| WBGene00021 | W09C3.4 | -0.5490803337 | 2.2041509794( | -0.9172647172 | 0.0185909049680459 |
| WBGene00021 | Y17G9B.4 | -0.7966602214 | 5.7569944155: | -1.4261303013 | 0.00106497933438308 |
| WBGene00021 | Y23H5A.2 | -1.0945016717 | 9.1152197231! | -2.3811841452 | 7.36218723704568e-09 |
| WBGene00021 | Y34F4.5 | -0.7026731061 | 0.0001722193: | -2.6436506051 | 1.51565881698569e-06 |
| WBGene00021 | Y37E3.8 | -0.4188115407 | 1.0771766830: | -1.7154742692 | 0.000787164040166247 |
| WBGene00021 | Y37E11B.6 | -0.4059580613 | 0.0003323120! | -1.3571301192 | 0.000715657494382445 |
| WBGene00021 | trap-3 | -0.4540699505 | 4.5844389568: | -0.9620114642 | 0.0445570152615297 |
| WBGene00021 | Y41D4A.6 | -0.4801431771 | 1.1721179223( | -1.4579073104 | 0.000320486715448534 |
| WBGene00021 | Y41D4B.4 | -0.4876519151 | 1.8273480328! | -1.0537044472 | 0.0309502660531166 |
| WBGene00021 | Y48G1A.1 | -0.8682530282 | 9.1116342104: | -1.3234509649 | 0.0344894948241989 |
| WBGene00021 | Y48G1A.2 | -0.9116787850 | 6.4610882719: | -1.6867620249 | 9.62941544957265e-08 |
| WBGene00021 | Y48G8AL.7 | -1.2318560494 | 3.8151022526( | -1.8470530799 | 0.000647797193650177 |

| | | | | | |
|---|---|---|---|---|---|
| WBGene00021 | Y51F10.2 | -0.1532859989 | 0.0047143709 | -0.8393745030 | 0.0137536777166226 |
| WBGene00021 | Y52E8A.2 | -0.6999644707 | 6.8495039883 | -0.9175905085 | 0.033401244700199 |
| WBGene00021 | fbxc-45 | -0.8995547434 | 2.0419500404 | -1.0971471779 | 0.0423961764505718 |
| WBGene00021 | Y57E12AL.1 | -0.5591954646 | 1.0236591696 | -1.0460755530 | 0.000805580688094815 |
| WBGene00021 | Y57E12AM.1 | -0.5515624020 | 1.0567917582 | -1.1167966137 | 0.00332591574427685 |
| WBGene00022 | Y65B4A.8 | -1.4296755999 | 3.7040115636 | -1.9891685626 | 5.5309454276613e-05 |
| WBGene00022 | icd-2 | -0.2508764850 | 1.3906534786 | -1.2167154033 | 0.0229703867155608 |
| WBGene00022 | Y67D2.7 | -0.5588214542 | 4.7079067502 | -0.8959477818 | 0.0178975239152386 |
| WBGene00022 | Y69A2AR.32 | -1.2348929808 | 5.4285726732 | -1.8395945715 | 9.92604209534234e-06 |
| WBGene00022 | Y71F9B.6 | -0.7634766859 | 7.1537967464 | -1.6117175204 | 0.00224760130662034 |
| WBGene00022 | chaf-2 | -0.5596415546 | 1.8002338798 | -0.7479035899 | 0.0387574681577992 |
| WBGene00022 | Y71H2AM.9 | -0.5325021017 | 1.5674117072 | -1.0008822446 | 0.0298449485909544 |
| WBGene00022 | Y73B3A.9 | -1.0145573303 | 7.9767176066 | -2.1327823672 | 0.00352998412119056 |
| WBGene00022 | sqd-1 | -0.3193374761 | 4.5309513101 | -0.7617651899 | 0.0203341362737953 |
| WBGene00022 | acp-6 | -0.9318708008 | 0.0001363725 | 3.7927656358 | 0.0252140795219324 |
| WBGene00022 | Y73B6BL.29 | -0.9171191916 | 2.1986679830 | -1.2065648803 | 0.0405802317934255 |
| WBGene00022 | Y73E7A.1 | -1.0584553818 | 2.0705818541 | -1.0611264213 | 0.00935324407502791 |
| WBGene00022 | Y82E9BR.3 | -0.4838325719 | 5.2288316424 | -0.8514628158 | 0.0441661636313187 |
| WBGene00022 | Y110A2AR.1 | -0.4685410720 | 6.8519581500 | -1.0821439495 | 0.00101619099710174 |
| WBGene00022 | ZC317.7 | -4.6509867740 | 6.3553188505 | -1.3963910584 | 0.0288845729948923 |
| WBGene00022 | ZC477.3 | -0.3984987408 | 9.1617541909 | -0.8517179520 | 0.0384223473069214 |
| WBGene00022 | rde-8 | -0.6572012435 | 3.0126591718 | -0.9859816522 | 0.00904663019082651 |
| WBGene00022 | acl-9 | -0.5279475497 | 1.0892980662 | -0.7044703950 | 0.0413361410622778 |
| WBGene00022 | npp-24 | -0.4596797763 | 1.1633101005 | -0.7731626199 | 0.0236839577576897 |
| WBGene00022 | ZK177.8 | -0.6690989055 | 5.9954305698 | -0.8732796890 | 0.00974430804973334 |
| WBGene00022 | tofu-7 | -0.2492342207 | 4.4728742118 | -0.8982578877 | 0.0274793045827988 |
| WBGene00022 | fubl-2 | -0.4771055795 | 1.5189125214 | -0.7636612748 | 0.0437482643927757 |
| **WBGene00022** | **oaz-1** | **-0.8578390169** | **1.2179384805** | **-1.0105455302** | **0.00210421422518981** |
| WBGene00022 | ZK484.6 | -1.1154321047 | 0.0033693276 | -2.7831903664 | 0.000787164040166247 |
| WBGene00022 | tag-307 | -0.6022710765 | 3.9173490138 | -0.8932012418 | 0.0175943731149859 |
| WBGene00022 | fahd-1 | -1.2928599743 | 2.3324809550 | -1.6424922525 | 0.0120983277433306 |
| WBGene00022 | ZK688.5 | -1.6504803602 | 0 | -2.0009152804 | 2.09669906748304e-11 |
| WBGene00022 | ZK688.9 | -0.4941641307 | 1.2946681575 | -0.9561099286 | 0.00839079332258292 |
| WBGene00022 | cin-4 | -1.4693017344 | 4.1095712411 | -2.3880927951 | 1.3523787616686e-12 |
| WBGene00022 | ZK1236.1 | -1.2390463702 | 3.0194472831 | -0.9154054185 | 0.0376430255149683 |
| WBGene00022 | ZK1248.11 | -0.4493983985 | 5.7641061861 | -1.2892168142 | 0.000169389591623116 |
| WBGene00022 | ZK1248.15 | -1.8058982048 | 1.4119587009 | -1.3091731401 | 0.000953639281157841 |
| WBGene00022 | ZK1248.19 | -1.4539176580 | 1.0120775760 | -1.1718843545 | 0.0376430255149683 |

| | | | | | |
|---|---|---|---|---|---|
| WBGene00022 | CD4.10 | -0.3866483261 | 0.0001036294 | -1.2878317745 | 0.0298449485909544 |
| WBGene00023 | rpl-41.2 | -0.5033065577 | 5.8456832969 | -1.7026958246 | 0.000386142291953241 |
| WBGene00023 | F14D2.15 | -0.8324231182 | 1.8835511312 | -1.0739673328 | 0.0261252586228313 |
| WBGene00043 | F32E10.9 | -2.4700672180 | 7.2781643624 | -3.8509759364 | 1.89325550357446e-05 |
| WBGene00044 | T01G1.4 | -0.3616939399 | 9.1604914267 | -0.8245783421 | 0.0437482643927757 |
| WBGene00044 | tag-229 | -0.5232140351 | 4.0573888788 | -0.9895855705 | 0.0162618944506573 |
| WBGene00044 | Y57G11C.51 | -1.1961131799 | 6.5822152055 | -1.7772982527 | 6.37005321944363e-05 |
| WBGene00044 | rad-8 | -0.5764270118 | 7.4641976544 | -0.8657215025 | 0.0103104356701034 |
| WBGene00044 | tag-280 | -0.8622471201 | 7.0380928104 | -1.6327550150 | 0.00276901369474864 |
| WBGene00044 | ZK688.11 | -1.5478345391 | 1.6804275106 | -2.7258451743 | 1.67298835968004e-13 |
| WBGene00077 | C27H6.9 | -0.7956612734 | 6.0736623312 | -1.5921547068 | 4.0176869867396e-05 |
| WBGene00138 | F58H1.8 | -0.5762370040 | 3.0961815311 | -1.1189594131 | 0.0103104356701034 |
| WBGene00185 | K11H3.8 | -0.4920384274 | 4.8700661770 | -0.9967414023 | 0.0408117353667565 |
| WBGene00194 | C53A5.16 | -0.6998274363 | 3.3035033059 | -1.3100937155 | 0.0249786768895591 |
| WBGene00206 | Y80D4G.1 | -1.2127096203 | 1.9591239151 | -1.9344894204 | 0.00474556665655936 |
| WBGene00219 | Y54G2A.73 | -0.5989044867 | 3.8306542635 | -1.7555539182 | 0.0039092944050503 |
| WBGene00219 | C06A1.12 | -0.3262577191 | 0.0011566124 | -0.8386970234 | 0.0430134412966845 |
| WBGene00255 | B0511.19 | -0.1989622757 | 0.0027421370 | -0.9970915119 | 0.00864605053815648 |
| WBGene00271 | T10E9.14 | -0.9849243213 | 1.7412500742 | -1.4656060100 | 0.0233558884336998 |
| WBGene00271 | D2089.8 | -0.8304200042 | 3.7709873629 | -2.0074685114 | 0.0161451716418977 |
| WBGene00000 | akt-2 | 1.1020782965 | 6.0569455632 | 2.1060266075 | 0.0284191845169195 |
| WBGene00000 | amx-1 | 0.5181813609 | 1.1943404238 | 1.6795571830 | 0.00783279954694634 |
| WBGene00000 | anc-1 | 0.8599211381 | 1.0425489310 | 1.2067675374 | 0.0442907004844807 |
| WBGene00000 | apl-1 | 0.7621579618 | 1.7219104179 | 1.4940605770 | 0.0199340180016796 |
| WBGene00000 | ark-1 | 0.4888538738 | 7.2176177241 | 1.0489781159 | 0.00971459101435518 |
| WBGene00000 | atf-5 | 0.4334275041 | 0.0472530754 | 1.3852991760 | 0.00953480849473906 |
| WBGene00000 | atf-6 | 0.5826578405 | 4.7775910231 | 1.5383363077 | 0.024159501943747 |
| WBGene00000 | ccg-1 | 0.7636046523 | 0.0001564242 | 1.4478239687 | 0.00413441580878045 |
| WBGene00000 | cdc-14 | 0.1244535507 | 0.0078732571 | 0.8122329540 | 0.0170106070434896 |
| WBGene00000 | cdh-4 | 0.5790201601 | 2.1232065475 | 1.2888998807 | 0.00231355529884879 |
| WBGene00000 | cpi-1 | 1.1687228057 | 6.7671410689 | 3.6304190998 | 3.57146439038755e-14 |
| WBGene00000 | cnt-1 | 0.3543465844 | 1.3886276833 | 1.3165892741 | 0.0377483985724088 |
| WBGene00000 | crn-4 | 0.7747891883 | 2.1879337965 | 1.1361653164 | 0.024165383077767 |
| WBGene00000 | dao-3 | 0.3853441770 | 0.0384945609 | -1.6559192391 | 0.0260030250996233 |
| WBGene00000 | dhs-14 | 0.6900431078 | 0.0327499559 | 4.7622859434 | 0.00380541493777915 |
| WBGene00000 | dhs-21 | 1.6635270496 | 2.9372371314 | 3.0643484766 | 0.000499516337434342 |
| WBGene00001 | dnj-27 | 0.6381905054 | 6.6931193980 | 2.0617480318 | 0.000689030759008703 |
| WBGene00001 | dpy-23 | 0.3173328664 | 0.0079796211 | 1.7483295446 | 0.00806352271954898 |

| | | | | | |
|---|---|---|---|---|---|
| WBGene00001 | dyn-1 | 0.7946768754 | 4.3435777240 | 1.1917978854 | 0.0280659064190795 |
| WBGene00001 | eef-1A.2 | 0.8471211998 | 1.2596234793 | 1.0234278863 | 0.0365902084867971 |
| WBGene00001 | emb-9 | 1.2093383378 | 1.2776827267 | 1.4730159083 | 0.00058745790955677 |
| WBGene00001 | fat-3 | 0.3923349060 | 0.0193264740 | 1.2589506364 | 0.0332656119558407 |
| WBGene00001 | fis-2 | 0.9406447966 | 1.0407023163 | 1.1953368697 | 0.0408014524631908 |
| WBGene00001 | fmo-4 | 1.5134110708 | 6.8744037111 | 1.2741311587 | 0.0235191322117101 |
| WBGene00001 | fog-1 | 0.4528502320 | 7.3857760379 | 0.8766318808 | 0.0135007178184262 |
| WBGene00001 | frm-1 | 0.6197274998 | 1.8594565846 | 0.8928104767 | 0.0156574305595428 |
| WBGene00001 | frm-7 | 0.2069901301 | 0.0112645075 | 1.1999455233 | 0.041945720655523 |
| WBGene00001 | gei-3 | 0.5729573051 | 0.0030175465 | 2.2094863555 | 0.00952187994619803 |
| WBGene00001 | meg-3 | 0.5047173490 | 7.3589237397 | 1.0474402171 | 0.00675876427329768 |
| WBGene00001 | gex-2 | 0.2370651289 | 0.0005885743 | 0.9415144250 | 0.0236748756689071 |
| WBGene00001 | gfi-2 | 0.1139333087 | 0.0387001779 | 0.9494083428 | 0.0205556531925389 |
| WBGene00001 | gst-4 | 1.7491597214 | 4.5530832215 | 2.1620422880 | 0.00058745790955677 |
| WBGene00001 | gst-24 | 0.9610852768 | 0.0312778821 | 4.0980694573 | 2.16364064048971e-13 |
| WBGene00001 | his-6 | 0.2369145399 | 0.0499369819 | -3.576591830 | 5.16268683725816e-10 |
| WBGene00001 | his-32 | 0.8524415767 | 0.0001078234 | -2.8387689854 | 2.32860711994388e-08 |
| WBGene00001 | his-38 | 0.9668533783 | 5.4122867797 | -2.7398611333 | 0.00942429983180476 |
| WBGene00001 | his-40 | 0.8531248123 | 8.0007743116 | -2.748731964 | 1.91616686475813e-06 |
| WBGene00001 | his-68 | 0.4334701702 | 7.1660634183 | -1.931781951 | 2.3345764972507e-07 |
| WBGene00002 | iff-2 | 0.4262601333 | 0.0022119600 | 0.9671727964 | 0.0346144563133377 |
| WBGene00002 | ina-1 | 0.5485535817 | 7.9386470797 | 1.3337286933 | 0.0215500955508471 |
| WBGene00002 | klp-4 | 0.9749674725 | 9.4821792151 | 1.7914495605 | 0.0454862984367192 |
| WBGene00002 | ksr-2 | 0.1643737165 | 0.0220234202 | 1.1011870901 | 0.00912555288157131 |
| WBGene00002 | lbp-1 | 0.4766272516 | 0.0387644302 | 1.3208429888 | 0.0129154452124318 |
| WBGene00002 | lbp-6 | 0.7660815735 | 5.1561536973 | 2.2302903144 | 7.51483618292984e-05 |
| WBGene00002 | lec-4 | 0.6294373339 | 9.4102901997 | 1.5000879812 | 0.0126937340895077 |
| WBGene00002 | lec-5 | 1.6310457707 | 2.0052454575 | 1.7036697539 | 0.00332096721720804 |
| WBGene00002 | let-2 | 1.1078401582 | 3.2147533148 | 1.5192123128 | 0.00014270063037508 |
| WBGene00002 | let-19 | 0.3001239561 | 0.0121407077 | 0.9173581014 | 0.0417295549414075 |
| WBGene00002 | let-413 | 0.3961836633 | 0.0008876045 | 1.5422284911 | 0.0387527791729912 |
| WBGene00002 | lim-7 | 1.2670508533 | 1.1198750434 | 1.8120165782 | 0.0373746614319323 |
| WBGene00003 | lys-7 | 1.0781872375 | 0.0149976420 | 5.4893042006 | 0.00035042184114175 |
| WBGene00003 | mes-1 | 0.3907052066 | 0.0024582393 | 1.3333098507 | 0.0351358120738652 |
| WBGene00003 | mex-3 | 0.1216477797 | 0.0147642849 | 0.8251514042 | 0.0381958853095823 |
| WBGene00003 | mig-15 | 0.5400429648 | 6.3814593635 | 1.4827463641 | 0.0373057477802643 |
| WBGene00003 | mlp-1 | 1.2230446537 | 4.8380444796 | 2.2501269948 | 0.00423697759441933 |
| WBGene00003 | mom-4 | 0.2407131092 | 0.0027333514 | 1.0842086041 | 0.00738368517951859 |

| | | | | | |
|---|---|---|---|---|---|
| WBGene00003 | msi-1 | 0.4153494187 | 0.0131650980 | 1.9024940238 | 0.0382061898732465 |
| WBGene00003 | dcap-2 | 0.3339309069 | 1.4231022224 | 0.8421081021 | 0.0403033346544568 |
| WBGene00003 | nfm-1 | 0.4488069335 | 0.0031128661 | 2.0435178213 | 0.0236922101583485 |
| WBGene00003 | nhr-84 | 1.3282030475 | 2.5515479479 | 2.1787788327 | 0.0461749028997023 |
| WBGene00003 | nmy-1 | 0.7925745031 | 1.6961405178 | 2.1869338375 | 2.01021136054315e-07 |
| WBGene00003 | nrf-5 | 1.2799694833 | 0.0009027124 | 2.7353800804 | 0.0021394879293794 |
| WBGene00003 | ost-1 | 1.0019045250 | 3.4466669398 | 1.4275097852 | 0.000361350013647058 |
| WBGene00003 | oxi-1 | 0.4159118594 | 2.0031981499 | 0.7450173178 | 0.0380826555033644 |
| WBGene00004 | parp-1 | 0.2091303938 | 0.0001292103 | 0.7840821778 | 0.0409304072927436 |
| WBGene00004 | tank-1 | 0.5993196355 | 2.6302174597 | 0.9990958363 | 0.0115327950954228 |
| WBGene00004 | pqn-21 | 1.2889561465 | 3.7766943790 | 1.7096740971 | 1.13012248214922e-06 |
| WBGene00004 | rbc-1 | 0.4795812995 | 8.7178881144 | 1.1851551629 | 0.0278609393263951 |
| WBGene00004 | rig-4 | 0.0791522264 | 0.0440569854 | 0.9656275374 | 0.0142933257088891 |
| WBGene00004 | rpl-25.1 | 1.0775420603 | 1.3737582916 | 1.4106870852 | 0.0011759418423118 |
| WBGene00004 | sdc-1 | 0.3816557010 | 0.0012860292 | 1.4204719485 | 0.00360746772975649 |
| WBGene00004 | sek-1 | 0.2667761238 | 0.0409804530 | 1.7809654540 | 0.0161518182402066 |
| WBGene00004 | sel-12 | 0.3681359068 | 0.0084995592 | 1.7240654712 | 0.0162911066951935 |
| WBGene00004 | sem-5 | 0.3064468854 | 0.0016679820 | 1.3554399906 | 0.0236774564002575 |
| WBGene00004 | sex-1 | 0.4891986842 | 0.0079109773 | 1.4100049199 | 0.00654898433652986 |
| WBGene00004 | skn-1 | 0.5214204275 | 2.2824070998 | 0.8661441212 | 0.02598056971727 |
| WBGene00004 | sli-1 | 0.5991511127 | 7.9630116664 | 1.2181060187 | 0.0169233036117202 |
| WBGene00004 | sop-2 | 0.3158238540 | 0.0235066792 | 1.3967132607 | 0.0269718755027512 |
| WBGene00005 | sqv-5 | 0.3242427981 | 2.6414720545 | 0.8824171479 | 0.0235551549873585 |
| WBGene00006 | sto-1 | 1.0204654622 | 4.6199150169 | 1.6172428994 | 0.00884188169677257 |
| WBGene00006 | sur-5 | 0.3671542929 | 0.0018364661 | 1.2139509065 | 0.0243138899128504 |
| WBGene00006 | taf-3 | 0.3051112326 | 0.0084362023 | 1.6678763919 | 0.00230390817754383 |
| WBGene00006 | lim-9 | 0.6152247776 | 0.0003650217 | 2.2689809460 | 0.0321132663638764 |
| WBGene00006 | kcc-1 | 0.3435803516 | 5.8444184639 | 1.3585557318 | 0.00293491880207218 |
| WBGene00006 | madd-3 | 0.3049574257 | 0.0021131258 | 1.4756061626 | 0.00924849189475974 |
| WBGene00006 | tap-1 | 0.6949040940 | 0.0006142091 | 1.5933529432 | 0.0325987156046246 |
| WBGene00006 | tba-4 | 0.5584689229 | 0.0090356193 | 1.5789510592 | 0.0146508527910821 |
| WBGene00006 | tbh-1 | 0.7251336081 | 0.0006364944 | 1.1708372198 | 0.0170106070434896 |
| WBGene00006 | tbx-33 | 0.6575210046 | 0.0008952139 | 1.7837740676 | 0.0039835275127462 |
| WBGene00006 | tol-1 | 0.1444669127 | 0.0266683322 | 0.9599028768 | 0.0176109583884111 |
| WBGene00006 | tpi-1 | 0.5276973764 | 6.3993603286 | 0.8781067163 | 0.02598056971727 |
| WBGene00006 | tps-1 | 1.1259097391 | 2.6997696875 | 1.6156719637 | 0.0152204464533136 |
| WBGene00006 | tra-3 | 0.2702735429 | 0.0134497469 | 1.1494122775 | 0.0483415143198372 |
| WBGene00006 | unc-26 | 0.3793229631 | 1.0528215683 | 1.0691617507 | 0.0103974794594557 |

| | | | | | |
|---|---|---|---|---|---|
| WBGene00006 | unc-32 | 0.1121139803 | 0.0120072060 | 0.8949553809 | 0.0199340180016796 |
| WBGene00006 | unc-33 | 0.9109148154 | 3.5923258856 | 1.2486295083 | 0.0133347993206383 |
| WBGene00006 | unc-40 | 0.1297922115 | 8.1161359133 | 0.8995140255 | 0.0273190189871856 |
| WBGene00006 | unc-53 | 0.8633214681 | 1.4464204574 | 2.4562105035 | 0.00329286963378462 |
| WBGene00006 | unc-112 | 0.3095816488 | 0.0026502287 | 1.5810699310 | 0.0140686438598502 |
| WBGene00006 | ver-3 | 0.8570502655 | 2.1338770379 | 1.7032544860 | 0.0176395758101266 |
| WBGene00007 | chd-7 | 0.3363363038 | 0.0187059812 | 1.1874800952 | 0.00714776898588186 |
| WBGene00007 | eri-12 | 0.4192792853 | 1.2379237232 | 1.2821320708 | 0.0273190189871856 |
| WBGene00007 | lron-3 | 1.2900750657 | 4.7689848235 | 1.8437274704 | 0.0173303157041122 |
| WBGene00007 | C04F12.1 | 1.1584337048 | 1.0790784888 | 1.6261412287 | 1.0420981813283e-06 |
| WBGene00007 | C05C9.3 | 1.9484588482 | 3.8182158113 | 2.9549318403 | 0.000386142291953241 |
| WBGene00007 | C07E3.9 | 1.4640896666 | 5.1617181525 | 1.8769537337 | 0.0337614792690729 |
| WBGene00007 | C18D4.6 | 1.5636699640 | 1.6269486134 | 2.7252933719 | 6.54708260381763e-07 |
| WBGene00007 | C27C12.1 | 0.5716901341 | 1.0778387487 | 1.5250410832 | 0.00138194782651652 |
| WBGene00007 | rsu-1 | 0.5102634587 | 6.3570560354 | 1.4318451088 | 0.000386626463845555 |
| WBGene00008 | C38D9.2 | 1.8033320585 | 2.8265765194 | 2.8930501871 | 0.00367463331249773 |
| WBGene00008 | din-1 | 0.4437217376 | 4.1012488928 | 0.9671483442 | 0.0173374876563467 |
| WBGene00008 | F09C8.2 | 0.5389750637 | 0.0010598161 | 1.9733798400 | 0.000320737008007391 |
| WBGene00008 | git-1 | 0.3945079038 | 0.0006054479 | 2.0132948372 | 0.0189327497567198 |
| WBGene00008 | F15D4.5 | 0.6700864052 | 9.8547651615 | 2.6313872651 | 0.0010297446474987 |
| WBGene00008 | F19C6.2 | 0.4554045069 | 0.0012039714 | 1.7794634835 | 0.0175693620017325 |
| WBGene00009 | aptf-4 | 1.4227287385 | 0.0015409089 | 4.0137447110 | 0.00100375272686938 |
| WBGene00009 | F28H6.4 | 0.6703458781 | 2.0807953530 | 1.3989312271 | 0.0006373069915679 |
| WBGene00009 | rict-1 | 0.2953738629 | 8.1739458901 | 1.5658813285 | 1.91616686475813e-06 |
| WBGene00009 | F31B9.3 | 1.3258710799 | 4.7343226144 | 1.0325071514 | 0.0490785725796147 |
| WBGene00009 | F31F6.2 | 0.7202802402 | 4.7111905268 | 1.3768204148 | 0.0382061898732465 |
| WBGene00009 | fasn-1 | 1.1335667482 | 5.6232297988 | 1.0608733914 | 0.0148101023668697 |
| WBGene00009 | F34D10.3 | 1.8420298928 | 6.1824464657 | 1.5733457243 | 0.0136686664329052 |
| WBGene00009 | F35E12.10 | 1.0701460995 | 1.5367281055 | 1.3640512559 | 0.0122456032813657 |
| WBGene00009 | cyld-1 | 0.1258898799 | 0.0095581195 | 1.1207093028 | 0.00169046382872165 |
| WBGene00009 | F45G2.10 | 0.5489376735 | 9.3178470525 | 1.1046589778 | 0.00294785822294085 |
| WBGene00009 | ipla-2 | 0.5679574341 | 5.2658786400 | 1.3038727090 | 0.0253967667973545 |
| WBGene00009 | gpdh-1 | 1.9926279060 | 0.0002617228 | 2.5902428540 | 0.0367410153257638 |
| WBGene00009 | F49E2.2 | 0.8084672018 | 0.0004822015 | 2.0816482482 | 0.0011759418423118 |
| WBGene00009 | F49E2.5 | 1.0197166399 | 8.4233344210 | 1.8081715942 | 0.00279156019689652 |
| WBGene00009 | swan-2 | 0.3527391712 | 0.0077700109 | 1.3507052536 | 0.0166892020772943 |
| WBGene00009 | swan-1 | 0.6467840640 | 1.3867627826 | 1.3527721812 | 0.0146508527910821 |
| WBGene00010 | F58G1.2 | 0.7446103090 | 9.5283866061 | 1.0400792769 | 0.0125197162631088 |

| | | | | | |
|---|---|---|---|---|---|
| WBGene00010 | copz-1 | 0.70306654258 | 2.06653745258 | 1.10656343279 | 0.00485831041281645 |
| WBGene00010 | ttr-51 | 1.89411691280 | 1.44246613590 | 2.04498577130 | 0.00179868565857806 |
| WBGene00010 | K02B7.3 | 2.22602914590 | 7.91771022350 | 3.63870332948 | 3.25218146319758e-05 |
| WBGene00010 | meg-1 | 0.55811545913 | 3.37796911293 | 0.84605368940 | 0.0235191322117101 |
| WBGene00010 | meg-2 | 0.53926106011 | 2.93943828462 | 1.52058647357 | 0.00646272191772102 |
| WBGene00010 | mak-1 | 0.52966579865 | 1.14052566155 | 1.04029953011 | 0.0108630392395372 |
| WBGene00010 | cysl-2 | 1.03455665340 | 2.07326015018 | 1.51784358470 | 0.00318451241078498 |
| WBGene00010 | M117.1 | 1.61688355664 | 1.12320642527 | 2.37942696680 | 0.00715469312478028 |
| WBGene00010 | ndfl-4 | 1.97939221551 | 1.70642880535 | 1.77143953489 | 0.00339211007142601 |
| WBGene00010 | nduo-1 | 1.12721260665 | 5.88275182028 | 1.53013463524 | 0.00652823330727302 |
| WBGene00010 | nduo-2 | 1.22421534760 | 4.17485084522 | 2.50826463213 | 2.7556999531596e-06 |
| WBGene00010 | nduo-5 | 1.11232030461 | 4.96962163674 | 2.66100593619 | 0.0261252586228313 |
| WBGene00011 | R10E4.1 | 0.47278635173 | 3.57450389030 | 1.19727835240 | 0.0227992157023324 |
| WBGene00011 | pck-2 | 0.31086048682 | 3.08006039889 | 1.01299887027 | 0.00558942081528504 |
| WBGene00011 | aakg-5 | 0.17658848633 | 0.00025702137 | 1.33512932420 | 0.000762435275597351 |
| WBGene00011 | T01E8.1 | 0.69708761749 | 0.00078599032 | 1.94384002078 | 0.0423296157548056 |
| WBGene00011 | T02G6.5 | 1.10453569433 | 1.22243641229 | 0.92980181752 | 0.0435121597973575 |
| WBGene00011 | aldo-1 | 0.49277911957 | 0.02835663242 | 1.47976180084 | 0.026111779600276 |
| WBGene00011 | nfya-1 | 0.32242162027 | 0.02864669409 | 1.12339157730 | 0.0296444447123829 |
| WBGene00011 | T12G3.2 | 1.07367580800 | 2.11161286512 | 1.63319537307 | 0.0242354245041834 |
| WBGene00011 | T18D3.1 | 0.52049522083 | 8.63274018717 | 1.32365151600 | 0.028088235706623 |
| WBGene00011 | prmt-9 | 0.34154805191 | 8.41267569057 | 0.86752562834 | 0.0199340180016796 |
| WBGene00012 | ucr-2.1 | 0.75368789060 | 1.93355723808 | 1.08471772824 | 0.0324317813872723 |
| WBGene00012 | W08E3.2 | 0.49757839705 | 4.46336262375 | 1.58603763975 | 0.024159501943747 |
| WBGene00012 | ztf-25 | 1.01925341183 | 1.03683593257 | 0.80058047020 | 0.0491753380503124 |
| WBGene00012 | mxt-1 | 1.27993492282 | 9.02364374398 | 1.93535053780 | 0.0178115489770295 |
| WBGene00012 | fbxb-7 | 1.40488554277 | 2.19285956904 | 1.76781621729 | 2.95814160843953e-06 |
| WBGene00012 | Y37D8A.4 | 0.85529543290 | 5.44765646883 | 1.06792443197 | 0.0261252586228313 |
| WBGene00012 | Y38E10A.14 | 0.43778690720 | 0.04266151890 | 1.84932855635 | 0.000156610483865346 |
| WBGene00012 | fbxa-150 | 0.67586123515 | 0.00063086514 | 3.38662872531 | 0.0180174722761054 |
| WBGene00012 | Y39B6A.10 | 0.80298488640 | 3.05684059280 | 1.31353393079 | 0.0332656119558407 |
| WBGene00012 | Y43F4B.7 | 0.31117286445 | 3.14789997463 | 0.98410034917 | 0.017768707167929 |
| WBGene00012 | cpt-1 | 0.50288283579 | 2.30670322155 | 0.99696587533 | 0.00798378887188276 |
| WBGene00012 | efhd-1 | 0.37417394150 | 0.02260136470 | 1.12469022815 | 0.0210972515941964 |
| WBGene00013 | cup-14 | 0.66041295800 | 3.12778539088 | 0.89217126524 | 0.00919825680043573 |
| WBGene00013 | Y67H2A.10 | 0.32785636992 | 4.69693510925 | 1.59611832445 | 3.07340127788268e-05 |
| WBGene00013 | fbxa-90 | 2.05071644948 | 5.75517402072 | 2.92729647002 | 0.0461766943311455 |
| WBGene00013 | Y105C5A.25 | 2.16448832317 | 1.05128459585 | 2.29775553018 | 0.0156574305595428 |

| | | | | | |
|---|---|---|---|---|---|
| WBGene00013 | fbxa-115 | 2.0545160461 | 9.5817824713 | 2.4641162925 | 0.0136093037324985 |
| WBGene00013 | cdk-11.2 | 0.4059267992 | 0.0009027645 | 1.1737998102 | 0.0238630248620487 |
| WBGene00014 | ZK637.14 | 0.5996520167 | 0.0009584524 | 1.1408823250 | 0.00803374610352617 |
| WBGene00014 | ZK930.2 | 0.8644844621 | 1.3023626859 | 1.2351205068 | 0.0212807746566505 |
| WBGene00014 | ZK1073.1 | 0.4155524252 | 0.0340224154 | 1.3607396105 | 0.0432109890811548 |
| WBGene00014 | C53C7.5 | 0.9784438661 | 6.6207727340 | 3.2767027055 | 0.000134374079802595 |
| WBGene00014 | F07H5.3 | 0.7537578641 | 1.1763356634 | 1.2370920401 | 0.000352530084109221 |
| WBGene00014 | F07H5.5 | 0.5063104977 | 0.0021022833 | 1.1415828903 | 0.00889060020941088 |
| WBGene00015 | B0041.8 | 0.3024315681 | 2.0131038616 | -0.888163767 | 0.0079488840503556 |
| WBGene00015 | B0212.3 | 0.8234954801 | 2.8946747264 | 1.2825375825 | 0.00336074959689006 |
| WBGene00015 | cpna-2 | 0.5620392579 | 0.0002928305 | 1.9745697262 | 0.0261252586228313 |
| WBGene00015 | B0281.5 | 1.0808803252 | 2.3584924808 | 1.6896086249 | 0.00230390817754383 |
| WBGene00015 | C03F11.4 | 1.9145820610 | 4.2396363891 | 2.7792824743 | 0.00530316637230017 |
| WBGene00015 | sor-3 | 0.5618736140 | 0.0010468180 | 1.5509455536 | 0.0136093037324985 |
| WBGene00015 | hyls-1 | 0.2896519729 | 3.5025753360 | 0.9166352248 | 0.0116603908719766 |
| WBGene00015 | C14F11.4 | 1.5573679684 | 5.2206147749 | 1.4179561873 | 0.00685635971153677 |
| WBGene00015 | kynu-1 | 1.4015246397 | 5.1495106167 | 2.8010962882 | 0.00180557581903888 |
| **WBGene00016** | **hpo-15** | **1.6351775913** | **8.5568999822** | **2.6239548208** | **0.0145584411311045** |
| WBGene00016 | cht-3 | 1.2804520491 | 6.2361699858 | 1.9380297248 | 2.00441266324475e-05 |
| WBGene00016 | eri-9 | 0.3938061711 | 5.3892439287 | 1.2080674141 | 0.000121099974662245 |
| WBGene00016 | sec-15 | 0.5403839422 | 1.3485508447 | 1.3558384208 | 0.0169233036117202 |
| WBGene00016 | lntl-1 | 0.9276228811 | 2.7638269972 | 1.6717889112 | 0.012934290340192 |
| WBGene00016 | C41A3.2 | 2.0914493918 | 5.0337900127 | 1.9373206059 | 0.00700749018329233 |
| WBGene00016 | C50A2.3 | 1.3598406989 | 6.0798567894 | 2.4824724765 | 0.00895635473259128 |
| WBGene00016 | C52A10.1 | 1.3135614864 | 2.1029461996 | 3.1357909313 | 0.00027638248706594 |
| WBGene00016 | fbxb-97 | 2.5561436961 | 1.2337751817 | 3.3029607351 | 7.81915863353169e-09 |
| WBGene00016 | pgph-3 | 0.9036737474 | 0.0052401598 | 3.5063134129 | 0.000294564619108989 |
| WBGene00016 | C53D5.1 | 0.5997740538 | 5.7275657295 | 1.3885259125 | 0.0210972515941964 |
| WBGene00016 | ztf-3 | 0.6019206436 | 1.9384068470 | 1.2502917755 | 0.0231050428490513 |
| WBGene00017 | utx-1 | 0.3574561832 | 0.0050208384 | 1.1303059110 | 0.0427430240124337 |
| WBGene00017 | D2092.4 | 1.5030120600 | 2.8277288436 | 1.8538478470 | 0.0409432326310322 |
| WBGene00017 | DC2.5 | 1.0165742999 | 0.0011937167 | 2.2166145158 | 0.043156163420299 |
| WBGene00017 | F08F3.6 | 0.4517034169 | 1.5808031407 | 0.8528662468 | 0.0486610012200231 |
| WBGene00017 | F10D7.5 | 0.5247011521 | 8.3556257475 | 1.3107213476 | 0.0147603758236877 |
| WBGene00017 | F10E9.3 | 0.4078067396 | 1.7195026412 | 1.3049738142 | 0.0102085934165896 |
| WBGene00017 | F13B9.1 | 0.4307572560 | 0.0219568710 | 0.9695644307 | 0.0331096221507528 |
| WBGene00017 | F13C5.2 | 0.6082839509 | 9.1577072274 | 1.7146066582 | 0.00378143298932292 |
| WBGene00017 | F21F3.7 | 0.3774112640 | 0.0030830686 | 1.2228232221 | 0.0120983277433306 |

| | | | | | |
|---|---|---|---|---|---|
| WBGene00017 | pals-34 | 0.7805840029 | 0.0361944361 | 1.6790448410 | 0.0200550783827705 |
| WBGene00017 | F31A3.5 | 0.5045535032 | 0.0005006892 | 1.3154228487 | 0.0437482643927757 |
| WBGene00018 | F35A5.1 | 1.2439540213 | 1.9515916629 | 1.2943279496 | 0.0341266246156233 |
| WBGene00018 | F37C4.5 | 0.6563427027 | 1.1465693083 | 1.3325123229 | 0.00219544124564106 |
| WBGene00018 | F44A2.5 | 0.8290884941 | 3.6215948515 | 2.6980777555 | 0.00236951555240148 |
| WBGene00018 | acs-1 | 0.9191818786 | 2.2812111557 | 1.2894551882 | 0.00873390708500994 |
| WBGene00018 | F46H5.7 | 1.0669302986 | 2.6691753500 | 1.8991408397 | 1.65613772220892e-05 |
| WBGene00018 | F55A3.2 | 0.4205818707 | 1.4192102637 | 0.9443436871 | 0.02598056971727 |
| WBGene00018 | glit-1 | 0.5669433884 | 0.0375093539 | 1.6877189260 | 0.0188912239323777 |
| WBGene00018 | F56C9.10 | 0.2945001941 | 4.9809896909 | 1.0131658022 | 0.0116616553137507 |
| WBGene00018 | F56F10.2 | 1.3491055956 | 0.0044984893 | 2.3146783044 | 0.0261252586228313 |
| WBGene00019 | spg-20 | 1.1178046173 | 6.8577737097 | 1.5857171403 | 3.02424394986255e-05 |
| WBGene00019 | ttr-47 | 1.2322912443 | 2.1568360237 | 2.0255358466 | 0.0463960888159386 |
| WBGene00019 | K10B3.5 | 0.5436248665 | 1.9090274909 | 1.7396351965 | 0.014976618573615 |
| WBGene00019 | K11H12.8 | 0.6597983185 | 3.6784005144 | 0.9583047392 | 0.0110737583840972 |
| WBGene00019 | sago-1 | 0.8940616817 | 4.4078655734 | 2.1244961038 | 1.60096930108931e-05 |
| WBGene00019 | ipla-1 | 0.3149637233 | 2.8831160736 | 1.1232218126 | 0.0029769490410596 |
| WBGene00019 | aass-1 | 0.9811200050 | 6.1885960338 | 1.1490908569 | 0.0328572059873037 |
| WBGene00020 | eppl-1 | 1.5628300609 | 1.0012690280 | 1.2423211592 | 0.0456233515442048 |
| WBGene00020 | aak-2 | 0.3260659054 | 0.0085479218 | 1.3117435793 | 0.020583124723461 |
| WBGene00020 | T05A12.4 | 2.4925234216 | 4.6075391976 | 1.1615911420 | 0.0378303338083639 |
| WBGene00020 | pho-14 | 1.3374497893 | 3.0851446357 | 1.4336333177 | 0.00836031682478536 |
| WBGene00020 | vha-15 | 0.4804442789 | 2.2045131269 | 1.0985605346 | 0.0175943731149859 |
| WBGene00020 | immt-1 | 0.5171500136 | 5.6255060500 | 1.1930149168 | 0.0192633305708344 |
| WBGene00020 | T19D12.6 | 1.0777058619 | 4.0972019846 | 1.2395130524 | 0.0418772570002025 |
| WBGene00020 | T20F5.6 | 0.1692285673 | 0.0001316756 | 0.7882935546 | 0.0320229663395672 |
| WBGene00020 | aakg-2 | 0.7588602058 | 6.5473592937 | 1.4140028116 | 0.0268879211098008 |
| WBGene00020 | T22B7.7 | 1.2659827739 | 0.0008273608 | 5.5910603232 | 1.11213158547634e-17 |
| WBGene00020 | T24A6.20 | 1.1111044771 | 6.5754832345 | 1.4127283623 | 0.0442907004844807 |
| WBGene00021 | W06A11.4 | 1.3959527756 | 2.3576110642 | 2.8133093480 | 0.00230390817754383 |
| WBGene00021 | W09C3.7 | 0.4340536318 | 0.0020717344 | 1.2248622182 | 0.00199066835822691 |
| WBGene00021 | Y18H1A.9 | 0.9541297430 | 0.0410039269 | 2.3262135856 | 0.0147603758236877 |
| WBGene00021 | Y34B4A.2 | 0.4314043037 | 0.0094753183 | 1.8909940080 | 0.00893643434105514 |
| WBGene00021 | Y37E11AM.2 | 0.4911930111 | 1.6220485069 | 1.1273616874 | 0.0437482643927757 |
| WBGene00021 | ppgn-1 | 0.4437222688 | 3.7371878296 | 0.9999571255 | 0.0236176868995733 |
| WBGene00021 | Y47G6A.3 | 2.4219423250 | 1.8016558058 | 2.5298054965 | 0.0112012834910723 |
| WBGene00021 | mis-12 | 0.5330174500 | 1.2159326745 | 1.1767577979 | 0.00545930197412032 |
| WBGene00021 | Y53G8AM.8 | 0.4674208152 | 4.2969694600 | 1.4942395288 | 0.0093802098399202 |

| | | | | | |
|---|---|---|---|---|---|
| WBGene00021 | Y54F10AM.5 | 0.58584358659 | 5.34504914125 | 0.78101315842 | 0.0422344819764757 |
| WBGene00021 | samt-1 | 1.43415325328 | 1.49992162069 | 2.01425824352 | 0.00529282784196875 |
| WBGene00022 | mppa-1 | 0.52331257233 | 4.53204629775 | 1.07090118545 | 0.0238630248620487 |
| WBGene00022 | Y71H10B.1 | 1.28794741320 | 9.22964550960 | 1.22282142040 | 0.0101966804763955 |
| WBGene00022 | Y119D3B.21 | 0.70249826671 | 0.01185987799 | 2.00615174911 | 0.0178187746756022 |
| WBGene00022 | ZC53.1 | 0.65959532588 | 4.26899082579 | 1.23651632682 | 0.0175693620017325 |
| WBGene00022 | cyy-1 | 0.22814392379 | 0.00053767458 | 1.07189961480 | 0.00372524059373357 |
| WBGene00022 | ZK484.3 | 0.21388896419 | 0.00063579302 | 0.93524810829 | 0.0349915165568052 |
| WBGene00022 | cth-2 | 1.19304099374 | 6.64049620668 | 1.32656092312 | 0.00809816817866127 |
| WBGene00022 | ZK1290.5 | 1.75363033069 | 2.85712787664 | 2.12792740450 | 0.000840298870140819 |
| WBGene00023 | rpl-41.1 | 1.66863310752 | 1.08933322308 | 1.15738685920 | 0.0292474849082205 |
| WBGene00023 | ZC513.7 | 0.70767595980 | 0.00011409820 | 2.47025725079 | 0.0017573618466768 |
| WBGene00023 | swm-1 | 2.99705042997 | 6.65800690132 | 2.14794120141 | 0.00675876427329768 |
| WBGene00023 | lin-15B | 0.98647883919 | 8.27207047620 | 1.68549450057 | 2.3795541151114e-06 |
| WBGene00023 | lin-15A | 0.89040262467 | 2.12940434790 | 1.18105647330 | 0.0166389848337867 |
| WBGene00044 | Y32F6A.6 | 1.09850364290 | 2.31994629520 | 1.24692839380 | 0.0286155137890765 |
| WBGene00044 | R04A9.7 | 1.90950128191 | 1.89538212298 | 4.72759863027 | 0.00027638248706594 |
| WBGene00045 | aho-3 | 0.24841414810 | 6.79966309779 | 1.02300960290 | 0.00754173239372364 |
| WBGene00045 | C08A9.11 | 1.25099483952 | 0.00195321734 | 3.11313829952 | 4.4302094283508e-06 |
| WBGene00077 | Y71A12B.19 | 0.66679886062 | 0.02141090889 | 2.55180211164 | 0.00244032702664022 |
| WBGene00219 | F56D6.16 | 0.93206319263 | 0.01958029900 | 3.68462114913 | 4.14751167725426e-05 |
