## Supplemental Table 2 for "SIN-3 transcriptional coregulator maintains mitochondrial homeostasis and polyamine flux"

**Table S2. List of gene commonly misregulated in *sin-3(tm 1276)* and *sin-3(syb2172)* mutants with SIN3 binding at promoter region**

| id_gene | gene_name |
|---|---|
| WBGene00022159 | mppa-1 |
| WBGene00021350 | Y37E3.8 |
| WBGene00020625 | mrrf-1 |
| WBGene00009161 | cox-7C |
| WBGene00007712 | mrpl-34 |
| WBGene00010419 | atp-1 |
| WBGene00009305 | metl-17 |
| WBGene00018961 | mrps-16 |
| WBGene00011759 | mrps-18B |
| WBGene00007686 | tomm-40 |
| WBGene00011273 | R53.4 |
| WBGene00011527 | cchl-1 |
| WBGene00012992 | mrpl-20 |
| WBGene00021849 | Y54F10AM.5 |
| WBGene00003967 | pdr-1 |
| WBGene00017121 | cyc-2.1 |
| WBGene00016844 | sucg-1 |
| WBGene00010317 | idh-1 |
| WBGene00013462 | micu-1 |
| WBGene00020275 | atp-4 |
| WBGene00009139 | F25H9.7 |
| WBGene00020511 | immt-1 |
| WBGene00012158 | ucr-2.1 |
