## Supplemental Table 3 for "SIN-3 transcriptional coregulator maintains mitochondrial homeostasis and polyamine flux"

Table S3

**Positive Mode Gradient**

| Time (mir | %A | %B |
|---|---|---|
| 0 | 2 | 98 |
| 3 | 2 | 98 |
| 11 | 30 | 70 |
| 12 | 40 | 60 |
| 16 | 95 | 5 |
| 18 | 95 | 5 |
| 19 | 2 | 98 |
| 25 | 2 | 98 |

**Negative Mode Gradient**

| Time (min) | %A | %B |
|---|---|---|
| 0 | 4 | 96 |
| 2 | 4 | 96 |
| 5,5 | 12 | 88 |
| 8,5 | 12 | 88 |
| 9 | 14 | 86 |
| 14 | 14 | 86 |
| 17 | 18 | 82 |
| 23 | 35 | 65 |
| 24 | 35 | 65 |
| 24,5 | 4 | 96 |
| 30 | 4 | 96 |
