## Supplemental Tables 4 for "SIN-3 transcriptional coregulator maintains mitochondrial homeostasis and polyamine flux"

| | A | B | C | D | E | F | G | H | I | J | K | L | M | N | O | P | Q | R | S | T | U | V | W | X | Y |
|---|---|---|---|---|---|---|---|---|---|---|---|---|---|---|---|---|---|---|---|---|---|---|---|---|---|
| 1 | Table S4 | | | | | | | | | | | | | | | | | | | | | | | | |
| 2 | | | | | | | | | | | | | | | | | | | | | | | | | |
| 3 | Metabolite Information | | | | QC | | | N2 | | | | | | | | *sin-3* | | | | | | | Statistics | | |
| 4 | | | | | | | | | | | | | | | | | | | | | | | | | |
| 5 | Name | formula | m/z, [M-H]- | RT | QC_001 | QC_002 | QC_003 | N2_001 | N2_002 | N2_003 | N2_004 | N2_005 | N2_006 | N2_007 | N2_008 | sin-3_002 | sin-3_003 | sin-3_004 | sin-3_005 | sin-3_006 | sin-3_007 | sin-3_008 | QC RSD % | log2 foldchange | p-value |
| 6 | alpha-keto-glutaric acid | C5H6O5 | 145,014 | 12,871 | 7047,11005 | 7560,5554 | 7519,6389 | 3084,28846 | 7530,61319 | 5775,513 | 8262,96591 | 6918,91004 | 7504,16663 | 4856,5724 | 5765,72243 | 5067,46432 | 12608,0186 | 5460,01317 | 5125,58683 | 6964,1791 | 9969,3826 | 2617,85566 | 3,9 | 0,14 | 0,6729 |
| 7 | succinic acid | C4H6O4 | 117,019 | 13,37 | 269573,54 | 193724,328 | 161026,78 | 232149,156 | 277230,574 | 275819,95 | 228816,228 | 266235,584 | 238396,713 | 179963,28 | 187676,311 | 159264,774 | 230636,059 | 229056,145 | 204421,768 | 127923,28 | 202819,14 | 248383,134 | 26,8 | -0,23 | 0,1154 |
| 8 | fumaric acid | C4H4O4 | 115,004 | 15,11 | 179158,036 | 173282,506 | 163298,79 | 139838,877 | 160023,717 | 183851,01 | 186557,559 | 157707,463 | 158577,951 | 109190,43 | 126795,747 | 169524,358 | 177257,015 | 102748,105 | 130980,291 | 108711,87 | 139163,9 | 126508,535 | 4,7 | -0,16 | 0,2710 |
| 9 | malic acid | C4H6O5 | 133,014 | 14,958 | 479555,382 | 431896,85 | 409168,68 | 362976,625 | 393252,498 | 518809,83 | 451428,902 | 425031,838 | 421302,944 | 278386,1 | 348688,852 | 419447,994 | 479178,495 | 265413,976 | 375062,023 | 296952,22 | 376239,47 | 352853,455 | 8,2 | -0,13 | 0,3852 |
| 10 | glucose-6-P | C6H13O9P | 259,022 | 19,756 | 54813,0516 | 60590,0108 | 71238,804 | 95148,2945 | 104832,748 | 144148,53 | 81081,0886 | 100731,901 | 100927,349 | 89459,131 | 55927,3117 | 49395,1641 | 95582,9309 | 88091,7371 | 89739,458 | 87138,503 | 170503,58 | 66461,501 | 13,4 | -0,06 | 0,8119 |
| 11 | glucose-1-P | C6H13O9P | 259,022 | 18,569 | 127191,493 | 131939,79 | 134144,3 | 106545,367 | 161334,981 | 220544,1 | 218989,766 | 164283,449 | 185612,186 | 158587,44 | 130122,591 | 82913,1706 | 119363,751 | 122131,423 | 117066,402 | 86429,6 | 150610,63 | 110516,078 | 2,7 | -0,58 | 0,0060 |
| 12 | Fructose-1,6-bis-P | C6H14O12P2 | 338,989 | 24,137 | 41918,1122 | 41004,6937 | 45264,501 | 30144,4863 | 50440,9609 | 65050,489 | 63504,8439 | 54278,1203 | 53881,4079 | 48226,792 | 41656,4202 | 25250,9967 | 41336,1502 | 45445,6276 | 41230,4289 | 33087,675 | 49101,522 | 39644,0813 | 5,2 | -0,37 | 0,0384 |
| 13 | trehalose | C12H22O11 | 341,109 | 8,312 | 607284,506 | 645171,169 | 732235,27 | 620674,097 | 802741,805 | 936072,95 | 1089218,75 | 755652,826 | 853015,225 | 851981,44 | 633826,178 | 484448,88 | 636988,219 | 1083398,92 | 678338,94 | 606937,89 | 901777,72 | 1067882,43 | 9,7 | -0,07 | 0,7254 |
| 14 | trehalose-6-phosphate | C12H23O14P | 421,075 | 21,441 | 87843,2286 | 97306,135 | 101212,5 | 83434,0841 | 84759,4153 | 140985,24 | 138356,825 | 100222,587 | 114432,527 | 95783,124 | 69462,2969 | 85281,7256 | 106007,881 | 95078,5202 | 99726,236 | 90702,633 | 145301,67 | 84636,4286 | 7,2 | -0,03 | 0,8421 |
| 15 | UDP-Glucose | C15H24N2O17P2 | 565,048 | 13,653 | 391237,591 | 448666,907 | 467662,22 | 291033,391 | 406465,598 | 488308,67 | 536713,693 | 410260,082 | 456003,729 | 398038,45 | 315754,394 | 352045,11 | 527458,834 | 544418,27 | 513138,354 | 436395,11 | 578133,84 | 487671,255 | 9,1 | 0,25 | 0,0770 |
| 16 | | | | | | | | | | | | | | | | | | | | | | | | | |
