## Supplemental Table 5 for "SIN-3 transcriptional coregulator maintains mitochondrial homeostasis and polyamine flux"

Table S5

| Metabolite Information | | | | QC | | | N2 | | | | | | | | *sin-3* | | | | | | | Statistics | | |
|---|---|---|---|---|---|---|---|---|---|---|---|---|---|---|---|---|---|---|---|---|---|---|---|---|
| Name | Formula | m/z, [M+H]+ | RT | QC_001 | QC_002 | QC_003 | N2_001 | N2_002 | N2_003 | N2_004 | N2_005 | N2_006 | N2_007 | N2_008 | sin-3_002 | sin-3_003 | sin-3_004 | sin-3_005 | sin-3_006 | sin-3_007 | sin-3_008 | QC RSD % | log2 foldchan | p-value |
| Alanine | C3H7NO2 | 90,055 | 10,466 | 18215,327 | 18382,5563 | 17832,0267 | 33196,1628 | 28760,9483 | 17401,0161 | 19480,1886 | 20498,0311 | 16053,2709 | 18000,0473 | 31542,9659 | 22682,5149 | 19618,8601 | 30379,6161 | 35988,3898 | 31985,5644 | 42146,0045 | 42520,3941 | 1,6 | 0,48 | 0,0505 |
| Arginine | C6H14N4O2 | 175,119 | 13,226 | 2661428,7 | 2700931,07 | 2733846,84 | 4590222,17 | 4918271,85 | 3446331,2 | 3307446,54 | 3653622,99 | 2979197,93 | 2977768,13 | 4088952,3 | 3799374,8 | 2956089,89 | 3947242,42 | 4219531,01 | 4137696,97 | 4600589,63 | 5257123,8 | 1,3 | 0,14 | 0,3173 |
| Asparagine | C4H8N2O3 | 133,061 | 11,063 | 5687,0887 | 6534,12577 | 6127,44896 | 11103,1789 | 11942,8727 | 6101,53018 | 7926,74142 | 8918,7258 | 6371,91071 | 6885,62543 | 12482,1531 | 7384,86078 | 6231,55771 | 12523,7409 | 12403,7401 | 11201,4306 | 14504,5378 | 17770,9525 | 6,9 | 0,39 | 0,1479 |
| Aspartic acid | C4H7NO4 | 134,045 | 12,106 | 4931,25129 | 5182,16197 | 4868,41125 | 6383,75063 | 10945,5183 | 6312,23599 | 7952,90832 | 6819,25934 | 8110,3417 | 7013,39668 | 8502,17505 | 5434,92044 | 4943,12697 | 4153,33683 | 4781,20929 | 4817,89291 | 5310,1037 | 5791,0606 | 3,3 | -0,62 | 0,0011 |
| Glutamine | C5H10N2O3 | 147,076 | 10,997 | 86861,8525 | 91770,6196 | 92816,1998 | 164162,9 | 176186,136 | 102315,183 | 121186,652 | 119597,205 | 91153,6278 | 101026,331 | 173798,174 | 125940,522 | 100371,385 | 172848,797 | 178932,726 | 175305,276 | 205743,833 | 231209,976 | 3,5 | 0,37 | 0,0887 |
| Glutamic acid | C5H9NO4 | 148,060 | 11,498 | 200987,716 | 210580,013 | 218247,262 | 376237,43 | 442531,41 | 240005,839 | 285650,921 | 318553,875 | 245064,619 | 241929,059 | 453781,267 | 303345,956 | 215180,491 | 400589,758 | 430968,173 | 385595,245 | 454525,494 | 554175,362 | 4,1 | 0,27 | 0,2229 |
| Histidine | C6H9N3O2 | 156,077 | 12,909 | 508906,068 | 521410,824 | 512659,604 | 707646,326 | 766268,272 | 639874,319 | 644119,207 | 614850,625 | 585593,209 | 549302,669 | 709335,213 | 606308,142 | 511964,727 | 567332,781 | 644665,795 | 623933,759 | 705151,97 | 734290,151 | 1,2 | -0,06 | 0,5364 |
| Isoleucine | C6H13NO2 | 132,102 | 8,324 | 61468,3089 | 62792,862 | 59150,46 | 102754,217 | 113173,723 | 79967,2527 | 64141,413 | 75505,1033 | 58322,0448 | 65472,2442 | 88783,0659 | 66109,3907 | 61854,4837 | 76584,4115 | 84072,1084 | 73971,2658 | 98975,7579 | 107970,466 | 3,0 | 0,01 | 0,9710 |
| Leucine | C6H13NO2 | 132,102 | 7,978 | 97706,7043 | 93881,3965 | 93089,6849 | 140626,725 | 164000,169 | 119112,248 | 96167,651 | 116177,791 | 93441,819 | 102475,422 | 142354,009 | 102429,721 | 103507,707 | 127498,281 | 138948,61 | 115821,036 | 175341,717 | 159593,802 | 2,6 | 0,11 | 0,4782 |
| Lysine | C6H14N2O2 | 147,113 | 13,488 | 170922,004 | 176531,806 | 180605,536 | 304628,845 | 321659,544 | 178699,167 | 183588,025 | 222019,173 | 158212,357 | 169570,874 | 305370,338 | 285611,706 | 206884,542 | 335422,269 | 386522,932 | 358911,428 | 442186,78 | 452991,499 | 2,8 | 0,61 | 0,0119 |
| Methionine | C5H11NO2S | 150,058 | 8,639 | 23988,6926 | 23624,5 | 24993,0287 | 38246,3225 | 41965,7144 | 34088,3409 | 27847,3298 | 28663,0428 | 27502,1143 | 26643,4083 | 35255,6668 | 20996,4108 | 24860,5219 | 30270,6912 | 29341,8533 | 22159,6871 | 41021,3349 | 29045,2108 | 2,9 | -0,20 | 0,2112 |
| Proline | C5H9NO2 | 116,071 | 9,285 | 266764,008 | 265874,102 | 272575,841 | 433345,01 | 488349,203 | 264362,703 | 228342,915 | 340564,104 | 200580,63 | 285018,337 | 396966,386 | 344884,696 | 322017,561 | 409418,999 | 495632,684 | 464265,916 | 583903,377 | 611632,159 | 1,4 | 0,49 | 0,0345 |
| Serine | C3H7NO3 | 106,050 | 11,066 | 4484,54364 | 4785,15546 | 5119,85135 | 7998,66681 | 7574,74326 | 4515,54207 | 5599,75922 | 5331,63532 | 4173,63853 | 4986,76272 | 9129,24465 | 7426,89881 | 5848,7913 | 12449,8247 | 12588,2573 | 11910,5573 | 12646,942 | 18990,7199 | 6,6 | 0,92 | 0,0128 |
| Threonine | C4H9NO3 | 120,066 | 10,561 | 5619,70737 | 6236,16641 | 6652,52568 | 9002,02358 | 11103,8532 | 5863,24646 | 7001,8818 | 7386,52561 | 5830,39644 | 6381,81572 | 10972,5478 | 8324,27443 | 6488,82826 | 11384,7599 | 13693,1245 | 11194,7789 | 15471,0269 | 18316,0419 | 8,4 | 0,61 | 0,0382 |
| Tryptophan | C11H12N2O2 | 205,097 | 8,055 | 95259,5576 | 102486,634 | 107182,599 | 98774,4556 | 117149,528 | 119855,95 | 138448,17 | 113048,181 | 112841,155 | 124011,215 | 129365,683 | 101509,338 | 97032,3489 | 113078,6 | 120060,282 | 127795,868 | 157329,022 | 148139,097 | 5,9 | 0,05 | 0,6575 |
| Tyrosine | C9H11NO3 | 182,081 | 8,972 | 171594,586 | 175684,058 | 141391,982 | 196164,598 | 206437,293 | 200590,146 | 179113,205 | 189374,514 | 156301,85 | 170352,925 | 176335,087 | 171238,894 | 161319,784 | 146379,262 | 178972,775 | 132264,1 | 173400,628 | 188684,118 | 11,5 | -0,16 | 0,0600 |
| Valine | C5H11NO2 | 118,086 | 8,548 | 6044491,87 | 5880055,62 | 5736347,3 | 8899125,29 | 9042259,87 | 7358026,51 | 5193066,55 | 7608835,78 | 5025599,88 | 5952129,11 | 7289489,48 | 9110593,06 | 7319331,21 | 6273516,06 | 7743425,12 | 8316083,34 | 8246326,33 | 10106200,3 | 2,6 | 0,21 | 0,1448 |
| GSH | C10H17N3O6 | 308,091 | 13,187 | 28891,1022 | 27940,8811 | 28539,4191 | 57536,8015 | 65428,8287 | 49803,6488 | 51017,5382 | 54138,8324 | 23554,6534 | 18358,0878 | 52802,6805 | 53306,4681 | 36896,1553 | 48680,0123 | 73487,3308 | 62001,2465 | 35067,7207 | 58595,5283 | 1,7 | 0,17 | 0,4576 |
| GSSG | C20H32N6O1 | 613,159 | 14,173 | 287152,651 | 308663,846 | 313130,145 | 775143,471 | 911324,723 | 371343,968 | 491976,406 | 623566,764 | 392134,267 | 441680,485 | 999426,364 | 406963,218 | 327090,308 | 620591,246 | 629619,239 | 749774,462 | 903651,445 | 842115,8 | 4,6 | 0,03 | 0,9065 |
| Ornithine | C5H12N2O2 | 133,097 | 13,562 | 8737,51998 | 8241,59094 | 9003,45149 | 17182,9169 | 17840,79 | 9935,54456 | 10152,7226 | 13270,1689 | 9129,47979 | 10163,533 | 17172,6025 | 13230,1526 | 9322,2725 | 13714,9557 | 15451,5794 | 15107,1895 | 16811,6075 | 17949,5858 | 4,5 | 0,15 | 0,4238 |
| Citrulline | C6H13N3O3 | 176,103 | 11,555 | 59772,4554 | 70200,6881 | 78408,1143 | 65917,8345 | 60346,611 | 40928,5709 | 36857,8755 | 46184,5767 | 36159,9539 | 47852,6159 | 55460,8195 | 123308,924 | 90618,5332 | 139717,046 | 168449,892 | 159424,418 | 171871,274 | 208254,05 | 13,4 | 1,64 | 0,0002 |
| Dimethyl argin | C8H18N4O2 | 203,150 | 12,232 | 18351,174 | 18703,6731 | 18112,2555 | 23022,403 | 22919,7242 | 16006,939 | 17942,0533 | 17509,1992 | 13576,1674 | 18045,3652 | 21727,7131 | 24110,9661 | 19432,9137 | 26781,2278 | 29492,7004 | 28394,2934 | 37904,2723 | 40038,9337 | 1,6 | 0,64 | 0,0075 |
| Sarcosine | C3H7NO2 | 90,055 | 10,683 | 1211,32239 | 1831,30296 | 1786,96192 | 2418,01284 | 2071,01829 | 955,2936 | 1552,29044 | 2125,38046 | 1479,39139 | 1476,31138 | 3810,13267 | 2087,22927 | 1652,87636 | 4105,86965 | 4772,89011 | 3253,02839 | 5295,27052 | 5579,17748 | 21,5 | 0,94 | 0,0209 |
| CoA | C21H36N7O1 | 768,123 | 8,815 | 1758,16349 | 3431,86225 | 3230,23985 | 4265,26097 | 4040,89264 | 2552,42763 | 3525,53537 | 3400,04645 | 1813,66968 | 1775,80351 | 2546,88584 | 6964,37063 | 4910,85486 | 5739,38934 | 3364,02936 | 3757,36494 | 2465,79068 | 4915,81149 | 32,6 | 0,62 | 0,0379 |
| CoA 2:0 | C23H38N7O1 | 810,133 | 12,767 | 14467,046 | 15122,2542 | 13806,4296 | 20657,1414 | 38327,1664 | 17450,6532 | 21250,7368 | 29579,3591 | 24374,1396 | 20983,1928 | 44951,7662 | 18968,894 | 12908,3767 | 29740,714 | 36703,3386 | 25892,0262 | 33789,2623 | 45216,9341 | 4,5 | 0,09 | 0,7387 |
| CoA 3:0 | C24H40N7O1 | 824,149 | 12,196 | 3192,40523 | 3249,68535 | 2947,38622 | 4729,98483 | 7930,07228 | 3198,42805 | 4112,95283 | 4388,9957 | 3815,94251 | 4113,26396 | 8299,56568 | 3822,54273 | 2456,33275 | 4434,84425 | 5089,17352 | 5886,13965 | 5713,03539 | 6311,03044 | 5,1 | -0,08 | 0,7676 |
| CoA 4:0 | C25H42N7O1 | 838,164 | 11,721 | 3710,4865 | 3651,89347 | 3543,84042 | 6960,3289 | 9627,76809 | 3622,56429 | 4584,80164 | 6668,52256 | 5046,92107 | 4006,271 | 10313,7262 | 4631,14124 | 4035,37031 | 9180,7322 | 9269,73446 | 8045,23958 | 9793,34137 | 13023,0212 | 2,3 | 0,38 | 0,2165 |
| CoA 5:0 | C26H44N7O1 | 852,180 | 11,332 | 15900,649 | 14990,7667 | 14019,31 | 28458,5933 | 32009,8764 | 15349,2411 | 16645,621 | 21397,1188 | 15342,2819 | 13596,0895 | 29289,2877 | 27402,1 | 19450,9774 | 39304,6017 | 37788,2121 | 37315,5355 | 37222,0996 | 49734,7998 | 6,3 | 0,72 | 0,0095 |
| Tiglyl-CoA | C26H42N7O1 | 850,164 | 11,534 | 9230,68191 | 8612,17641 | 8740,30476 | 14738,3246 | 21256,1648 | 11153,6472 | 12214,3118 | 13056,6976 | 12471,883 | 10846,1343 | 19282,2262 | 16349,5867 | 11860,8042 | 21630,4733 | 20366,5157 | 20930,2962 | 17222,4922 | 29091,0835 | 3,7 | 0,45 | 0,0550 |
| SAM | C15H22N6O5 | 399,145 | 13,361 | 760954,986 | 778463,181 | 767318,08 | 1282426,96 | 1469450,14 | 971330,083 | 1135167,74 | 1161346,5 | 906641,333 | 947169,145 | 1223209,35 | 863078,78 | 617048,233 | 750555,96 | 858239,846 | 864189,876 | 930022,265 | 1008629,2 | 1,2 | -0,43 | 0,0038 |
| SAH | C14H20N6O5 | 385,129 | 10,681 | 9517,48064 | 10450,9558 | 9991,53995 | 21748,8045 | 24697,4934 | 13949,6462 | 15693,7056 | 16532,2089 | 11621,9297 | 12666,3671 | 22500,5775 | 11470,9652 | 9658,02581 | 15309,0661 | 17877,8261 | 14458,4729 | 20102,2078 | 22231,4224 | 4,7 | -0,13 | 0,5352 |
| Cystathionine | C7H14N2O4S | 223,075 | 13,255 | 704570,4 | 715318,857 | 725363,285 | 1480454,2 | 1776751,75 | 849329,427 | 1026500,8 | 1185490,45 | 857766,417 | 875989,895 | 1686992,03 | 1083511,67 | 773232,37 | 1472286,82 | 1574723,65 | 1423594,92 | 1680265,6 | 1923054,99 | 1,5 | 0,22 | 0,3281 |
| S-Adenosylme | C14H23N6O3 | 355,155 | 14,168 | 5480,78994 | 5474,69932 | 5207,0464 | 10151,8161 | 10826,7069 | 6763,58757 | 7328,95001 | 11379,5457 | 10067,6082 | 11662,6729 | 10502,0011 | 2884,71215 | 2170,63904 | 2307,28836 | 2385,76041 | 3275,04723 | 2349,07248 | 3075,02128 | 2,9 | -1,90 | 0,0000 |
| 5'-Methylthio: | C11H15N5O3 | 298,097 | 13,359 | 40606,3406 | 40907,1755 | 40645,6548 | 69239,1106 | 79782,3152 | 51935,2017 | 58465,5326 | 64511,8583 | 48198,6555 | 49972,1335 | 66801,5261 | 47737,3636 | 34220,1647 | 43389,3716 | 49348,39 | 48971,5898 | 50690,0013 | 55571,8884 | 0,4 | -0,37 | 0,0110 |
| Succinyl-coa | C25H40N7O1 | 868,139 | 13,861 | 1528,59377 | 1769,7061 | 1470,12612 | 2403,55066 | 7915,64501 | 2261,90898 | 3371,55828 | 3274,32712 | 2902,26641 | 3848,47842 | 5350,64266 | 2818,36257 | 2788,22592 | 5108,82945 | 4575,81369 | 5580,77781 | 3692,32985 | 7864,90132 | 10,0 | 0,24 | 0,4634 |
| Folic acid | C19H19N7O6 | 442,147 | 11,001 | 2120,63183 | 3199,21995 | 2809,28986 | 4768,79866 | 4406,60481 | 3120,9102 | 3263,37194 | 2635,62295 | 1896,87311 | 2142,11755 | 3304,86673 | 3578,1291 | 2778,53543 | 2760,08681 | 3593,01543 | 3470,86703 | 3388,35186 | 4076,95996 | 20,2 | 0,08 | 0,6502 |
| 5-methyl tetral | C20H25N7O6 | 460,194 | 11,071 | 2148,99577 | 2961,31492 | 2407,88103 | 5559,30151 | 6496,10956 | 2943,67112 | 1934,18766 | 3548,72066 | 2274,94435 | 2872,5004 | 5643,1422 | 3637,90655 | 2447,47738 | 4680,00495 | 4442,42271 | 4755,7894 | 5445,02428 | 7346,9857 | 16,6 | 0,26 | 0,3762 |
| Dihydrofolic a | C19H21N7O6 | 444,163 | 12,911 | 12879,1477 | 9679,51134 | 8296,37975 | 8948,04492 | 11272,5145 | 8854,0216 | 10144,445 | 9488,98353 | 10477,1499 | 7472,68722 | 9408,01282 | 9995,17336 | 10086,807 | 7103,83494 | 8015,9789 | 7556,18995 | 7865,07968 | 9215,2777 | 22,9 | -0,15 | 0,1417 |
| Carn 2:0 | C9H17NO4 | 204,123 | 8,301 | 805367,307 | 826284,95 | 787852,792 | 948193,139 | 1320194,42 | 948559,603 | 863967,836 | 1116101,11 | 861126,218 | 1046198,86 | 1452350,16 | 1179098,82 | 967034,781 | 1134951,62 | 1348211,16 | 1456032,2 | 1031494,63 | 1910938,31 | 2,4 | 0,27 | 0,1562 |
| Carn 3:0 | C10H19NO4 | 218,139 | 6,615 | 413049,665 | 411022,625 | 411507,117 | 607302,85 | 762442,332 | 434615,921 | 429424,189 | 524004,045 | 384426,798 | 514064,688 | 675027,164 | 537722,581 | 479689,322 | 605595,59 | 663186,393 | 591811,226 | 552155,691 | 1076975,99 | 0,3 | 0,25 | 0,2737 |
| Carn 4:0 | C11H21NO4 | 232,154 | 5,121 | 341525,343 | 339732,624 | 335636,414 | 496724,005 | 552651,796 | 319657,177 | 324144,514 | 414898,521 | 290341,504 | 359078,797 | 548640,88 | 351640,425 | 322963,509 | 597865,197 | 641685,178 | 520791,504 | 670092,456 | 931349,799 | 0,9 | 0,48 | 0,0939 |
| Carn 5:0 | C12H23NO4 | 246,170 | 4,149 | 628906,055 | 635604,505 | 638117,445 | 954763,051 | 1019029,03 | 637725,086 | 648994,118 | 768624,605 | 541485,98 | 648508,682 | 932102,028 | 769616,98 | 701244,142 | 957199,082 | 1000352,36 | 1008485,4 | 1068913,7 | 1385308,45 | 0,8 | 0,36 | 0,0632 |
| Tiglyl-Carnitin | C12H21NO4 | 244,154 | 4,525 | 47611,8769 | 47435,1297 | 47903,9502 | 65421,0842 | 84625,6107 | 54583,0994 | 50643,1521 | 59337,176 | 45971,3777 | 50844,9931 | 74318,7673 | 39566,3726 | 52967,3638 | 68443,6305 | 72734,1667 | 34067,5692 | 85577,264 | 91290,7472 | 0,5 | 0,07 | 0,7758 |
| Riboflavine | C17H20N4O6 | 377,146 | 4,133 | 193243,842 | 200878,167 | 190971,746 | 203210,45 | 239804,571 | 209496,861 | 264989,094 | 214639,309 | 209910,768 | 184449,945 | 243535,334 | 130152,609 | 153631,1 | 188353,965 | 191202,951 | 155198,679 | 209962,719 | 176951,948 | 2,7 | -0,36 | 0,0038 |
| Pantothenic a | C9H17NO5 | 220,118 | 3,17 | 407315,721 | 402670,904 | 399152,07 | 439885,919 | 505295,888 | 312939,979 | 369444,277 | 438263,672 | 308371,518 | 354670,852 | 599646,678 | 515729,828 | 382360,096 | 507395,797 | 673240,102 | 606344,984 | 656146,777 | 925465,653 | 1,0 | 0,55 | 0,0271 |
| N-Acetyl sperm | C9H21N3O | 188,176 | 13,749 | 1328246,04 | 1300711,18 | 1258667,76 | 1995949,76 | 2049425,21 | 1193640,77 | 1237151,84 | 1925244,21 | 1127578,13 | 1417278,85 | 2010105,83 | 1937582,88 | 1379200,59 | 1778273,25 | 2051165,36 | 2245003,4 | 2559793,33 | 2713575,93 | 2,7 | 0,37 | 0,0569 |
| Agmatine | C5H14N4 | 131,129 | 13,608 | 12351,2498 | 12166,4502 | 11179,8977 | 9122,32002 | 10319,8289 | 8916,08924 | 8277,31597 | 10089,6811 | 7691,2395 | 9089,48231 | 10359,88 | 16766,1111 | 14121,4487 | 15364,2098 | 18393,4596 | 15683,0247 | 25399,5071 | 18623,1036 | 5,3 | 0,94 | 0,0007 |
| Cadaverin | C5H14N2 | 103,123 | 13,837 | 3427,66539 | 3234,80975 | 3134,82611 | 3574,85772 | 3385,97443 | 3774,42736 | 2695,55025 | 3971,69413 | 3008,44104 | 2832,3371 | 3089,75233 | 6092,74413 | 3929,94574 | 3533,63996 | 4999,80567 | 4513,02667 | 5414,64261 | 5734,85792 | 4,6 | 0,57 | 0,0032 |
| Putrescine | C4H12N2 | 89,107 | 14,123 | 16355,945 | 15778,6703 | 15739,9191 | 23152,4004 | 22006,8216 | 21200,5933 | 15159,8501 | 18063,1725 | 16959,6567 | 16480,0716 | 15305,4898 | 24547,9069 | 18910,0242 | 15074,7835 | 17647,0415 | 19196,1128 | 18317,2014 | 20825,336 | 2,2 | 0,05 | 0,6738 |
| N-Acetyl putre | C6H14N2O | 131,118 | 8,911 | 57465,8973 | 56251,3295 | 54163,9012 | 120457,422 | 129844,647 | 74167,01 | 60949,7036 | 83481,9714 | 58919,4467 | 75909,6684 | 103255,265 | 68649,7709 | 56128,3428 | 71441,7713 | 84135,3278 | 77371,4227 | 89802,0141 | 118865,13 | 3,0 | -0,13 | 0,5476 |
| N-Acetyl sperm | C12H28N4O | 245,234 | 15,227 | 6836,0567 | 6945,0949 | 7109,31012 | 1869,29785 | 1907,31569 | 1478,98019 | 1439,80773 | 3068,55053 | 1747,18961 | 3703,2866 | 4602,12381 | 14734,6829 | 3819,3599 | 10553,4855 | 15817,0963 | 11362,5373 | 16996,6245 | 22530,5546 | 2,0 | 2,47 | 0,0021 |
| Serotonin | C10H12N2O | 177,102 | 11,562 | 5104,50717 | 6054,79398 | 6932,29965 | 5575,19555 | 4671,11261 | 3139,05937 | 3384,49376 | 3750,93893 | 2675,99142 | 4283,95344 | 4647,08883 | 9987,68696 | 7411,98972 | 11294,3662 | 13059,4665 | 13278,1858 | 13773,5174 | 17103,8691 | 15,2 | 1,61 | 0,0003 |
| Saccharopine | C11H20N2O6 | 277,139 | 13,358 | 41324,0667 | 41702,3704 | 44943,1797 | 73485,7029 | 76408,5898 | 39360,1835 | 48789,5768 | 58961,68 | 44231,394 | 44715,7945 | 80870,2986 | 78629,9104 | 51627,2411 | 102977,177 | 121170,797 | 94484,2296 | 129126,658 | 140493,861 | 4,7 | 0,81 | 0,0081 |
| L-2-Aminoadi | C6H11NO4 | 162,076 | 11,022 | 106777,629 | 113189,451 | 108486,115 | 189149,151 | 216287,564 | 108258,825 | 133824,646 | 154086,233 | 109323,941 | 116848,524 | 250678,216 | 164963,924 | 115692,509 | 232717,323 | 254264,869 | 230533,866 | 286372,651 | 348025,359 | 3,0 | 0,55 | 0,0577 |
| Spermidine | C7H19N3 | 146,165 | 15,408 | 339471,282 | 344154,901 | 344695,016 | 75176,974 | 98926,8377 | 59816,9739 | 50831,3831 | 151308,521 | 82388,873 | 167849,151 | 206096,104 | 761536,673 | 149504,72 | 428390,93 | 526334,479 | 410079,547 | 452880,577 | 836559,015 | 0,8 | 2,19 | 0,0034 |
